# Functional regimes define the response of the soil microbiome to environmental change

**DOI:** 10.1101/2024.03.15.584851

**Authors:** Kiseok Keith Lee, Siqi Liu, Kyle Crocker, David R. Huggins, Mikhail Tikhonov, Madhav Mani, Seppe Kuehn

**Affiliations:** Department of Ecology and Evolution, The University of Chicago, Chicago, IL 60637, USA; Center for the Physics of Evolving Systems, The University of Chicago, Chicago, IL 60637, USA; Department of Engineering Sciences and Applied Mathematics, Northwestern University, Evanston, IL 60208, USA; NSF-Simons Center for Quantitative Biology, Northwestern University, Evanston, IL 60208, USA; USDA-ARS, Northwest Sustainable Agroecosystems Research Unit, Pullman, WA 99164, USA; Department of Physics, Washington University in St. Louis, St. Louis, MO 63130, USA; National Institute for Theory and Mathematics in Biology, Northwestern University and The University of Chicago, Chicago, IL; Center for Living Systems, The University of Chicago, Chicago, IL 60637, USA

## Abstract

The metabolic activity of soil microbiomes plays a central role in carbon and nitrogen cycling. Given the changing climate, it is important to understand how the metabolism of natural communities responds to environmental change. However, the ecological, spatial, and chemical complexity of soils makes understanding the mechanisms governing the response of these communities to perturbations challenging. Here, we overcome this complexity by using dynamic measurements of metabolism in microcosms and modeling to reveal regimes where a few key mechanisms govern the response of soils to environmental change. We sample soils along a natural pH gradient, construct >1500 microcosms to perturb the pH, and quantify the dynamics of respiratory nitrate utilization, a key process in the nitrogen cycle. Despite the complexity of the soil microbiome, a minimal mathematical model with two variables, the quantity of active biomass in the community and the availability of a growth-limiting nutrient, quantifies observed nitrate utilization dynamics across soils and pH perturbations. Across environmental perturbations, changes in these two variables give rise to three functional regimes each with qualitatively distinct dynamics of nitrate utilization over time: a regime where acidic perturbations induce cell death that limits metabolic activity, a nutrient-limiting regime where nitrate uptake is performed by dominant taxa that utilize nutrients released from the soil matrix, and a resurgent growth regime in basic conditions, where excess nutrients enable growth of initially rare taxa. The underlying mechanism of each regime is predicted by our interpretable model and tested via amendment experiments, nutrient measurements, and sequencing. Further, our data suggest that the long-term history of environmental variation in the wild influences the transitions between functional regimes. Therefore, quantitative measurements and a mathematical model reveal the existence of qualitative regimes that capture the mechanisms and dynamics of a community responding to environmental change.

## Introduction

The metabolic activity of soil, marine, and freshwater microbiomes drives carbon and nitrogen transformations that sustain biogeochemical cycles and life in the biosphere [1–3]. These microbiomes are also subjected to environmental perturbations including changes in temperature, pH, moisture, oxygen, and nutrients stemming from natural and anthropogenic events. As such, in order to predict the effect of climate change on global nutrient cycles, it is necessary to understand how microbiome metabolism responds to environmental change in nature.

Determining how environmental change impacts community metabolism has proven vexing because of the complexity of natural microbiomes. This complexity is perhaps most apparent in soils, which possess immense taxonomic diversity [4], spatial heterogeneity [5], and chemically diverse environments [6]. As a result, environmental perturbations can modify collective metabolic activity in many ways, from direct changes in microbial composition, physiology [7], and ecological interactions [8, 9] to indirect modification of nutrient availability [10–12] and spatial organization [5, 13]. Thus, a key question arises: which mechanisms are important for determining the metabolic response of complex microbiomes to environmental change?

Large-scale surveys approach this question by quantifying correlations between environmental variation, community composition, and metabolic processes in the wild [14–25]. Although surveys have revealed robust correlations, they face two challenges in uncovering the mechanisms determining community response to environmental change. First, and most importantly, surveys do not allow control for confounding factors, such as correlated environmental variables, rendering any causal inference infeasible. Second, it is difficult to quantify metabolic dynamics in situ on a large scale in the wild. As a result, surveys have limited power to determine the mechanisms that govern the metabolic response to environmental change in natural communities.

To control for confounding factors and gain mechanistic insights, we use soil microcosms, which remove correlated environmental fluctuations and permit controlled perturbations in the lab. To further control for confounding factors, these soils are sourced from a single site [26, 27] that exhibits large natural pH variation but minimal variability in other environmental factors (e.g., climate, moisture, soil texture, C/N ratio). Leveraging insights from global surveys, we focus on pH – the environmental variable that shows a strong correlation with soil microbiome composition and features of metabolism. [9, 15, 25, 28, 29]. Second, soil microcosms enable high-throughput quantification of metabolic time series in response to environmental perturbations. Our metabolic measurements focus on a key functional process in nitrogen cycling, the anaerobic respiration of nitrate which is ubiquitously performed by complex communities of soil bacteria, in response to natural and applied changes in pH.

Here, we measure nitrate utilization dynamics in >1500 microcosms across a wide range of natural and laboratory-induced pH changes. Next, we develop a judicious mathematical framework that accurately describes nitrate utilization dynamics across all microcosms. Our model shows that changes in functional dynamics in response to pH perturbations can be mechanistically understood by considering just two variables: the quantity of biomass in the community actively utilizing nitrate and the availability of growth-limiting nutrients. These two parameters emerge naturally from our mathematical model using only the community-level nitrate uptake data. The model predicts that changes in pH alter nitrate utilization dynamics by differentially affecting both the quantity of active biomass and the availability of nutrients that limit its growth.

As a result, despite the ecological, chemical, and spatial complexity of soils, we find that the functional response of the soil microbiome to changes in pH can be categorized into three mechanistically distinct regimes demarcated by the levels of these two variables. Each functional regime is defined by which of the two variables exerts greater control over nitrate utilization rates. During moderate pH perturbations, metabolic rates are set by the pH-mediated release of nutrients from soil particles that limit the growth of a large metabolically active biomass (Nutrient-limiting regime, Regime II). When soils are subjected to large basic perturbations, massive nutrient release relieves the nutrient limitation, but the dominant taxa are no longer metabolically active, and metabolism is set by the rapid growth of initially rare taxa (Resurgent growth regime, Regime III). During large acidic perturbations, functional responses are limited by the pervasive death of the active biomass in the community (Acidic death regime, Regime I). The transition between functional regimes can be abrupt (from Regime II to III) or smooth (from Regime I to II) as pH is varied and depends on the long-term pH history of the soil. Thus, while the dynamics and mechanisms of each functional regime are conserved across soils, the transitions between regimes depend on environmental history and community composition. Our study demonstrates a generalizable approach wherein high-throughput soil microcosm experiments coupled with mathematical models can overcome the complexity of natural ecosystems to mechanistically reveal the specific microscopic processes that contribute to the microbiome’s response to environmental change.

## Results

Nitrate (NO_3_*^−^*), which has critical implications for agriculture and climate, is reduced in soils when bacteria use it as an electron acceptor during anaerobic respiration in the absence of oxygen. Both denitrification (NO_3_*^−^ →* NO_2_*^−^ →*..*→* N_2_) or dissimilatory nitrate reduction to ammonia (DNRA, (NO_3_*^−^ →* NO_2_*^−^ →* NH_4_^+^) reduce nitrate to nitrite (NO_2_*^−^*) while consuming organic carbon. Due to the importance of pH in microbial physiology and soil chemistry, decades of studies have examined how pH affects nitrate reduction [30]. However, discrepancies in experimental methods (Table S1) and limited modeling, have made it difficult to find principles governing metabolic responses to pH perturbations [17] (Table S2).

### Metabolite dynamics in soils after short and long-term pH perturbations

To address this problem we measured nitrate utilization dynamics in soil microcosms across a range of native and perturbed pH levels. We sampled 20 top soils with pH from 4.7 to 8.3 (Fig. 1A, Table S3) at the Long-term Agricultural Research Cook Agronomy Farm (CAF) (Pullman, WA, USA). Sampled soils had similar characteristics (Table S3) which minimized the effects of confounding factors that might alter metabolic responses to perturbations. At this site, long-term variation in soil pH arises from local agricultural practices and erosion.

**Figure 1:**
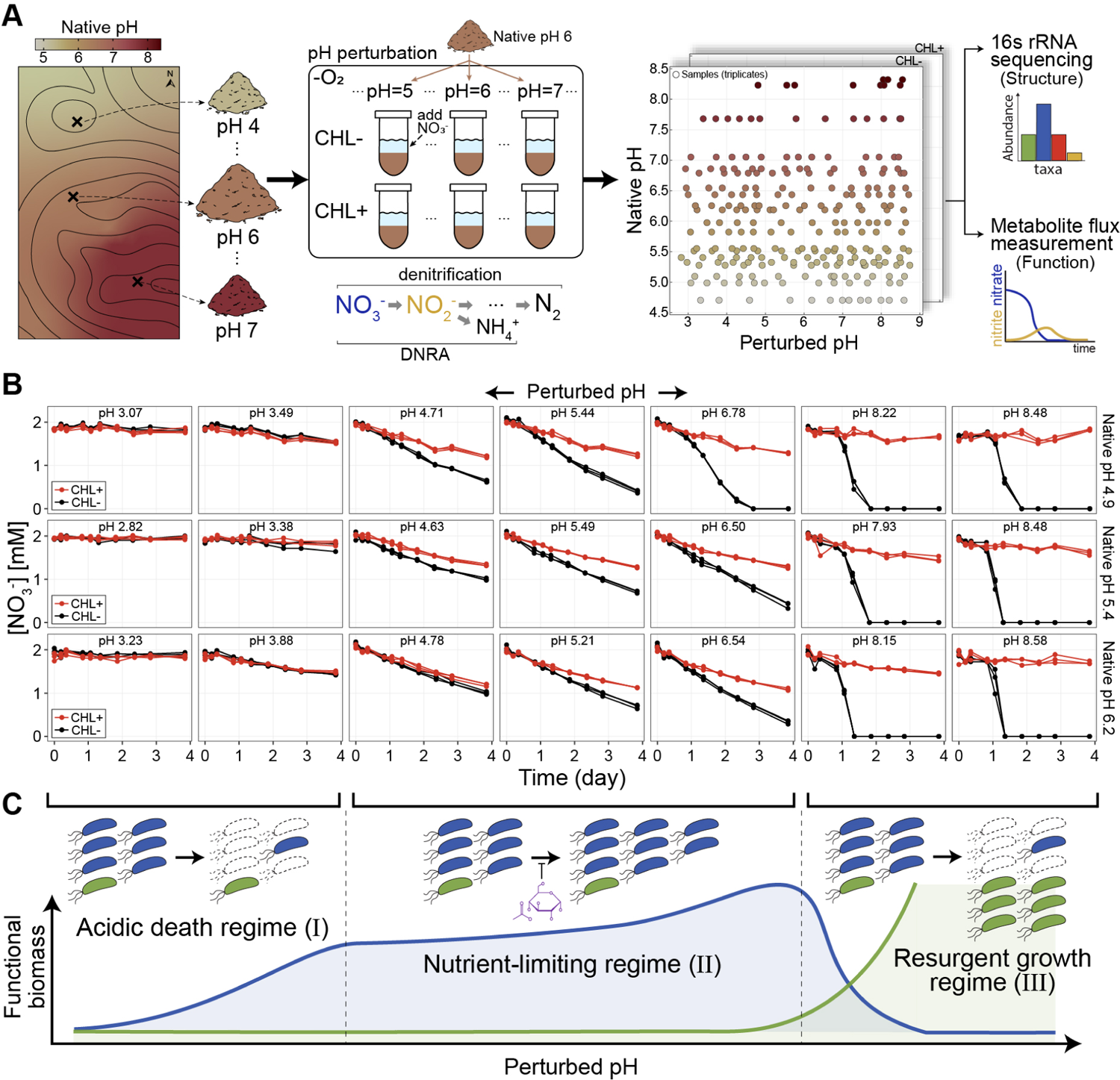
Soil microbiome metabolite and abundance dynamics under long and short-term pH variation. **(A)** Schematic of the field sampling for soils with long-term pH variations (20 soils, pH 4.7 to 8.3, Cook Agronomy Farm, Pullman, WA, USA) and the experimental setup for imposing short-term pH perturbations in laboratory conditions. With each of the 20 soils, we created slurries (1:2 soil:water) amended with 2mM nitrate, adjusted to 13 different pH levels, and treated with (CHL+, no growth) or without chloramphenicol (CHL-, growth). *∼*1,500 microcosms are depicted in a grid of different pH conditions (perturbed pH vs. native pH) each condition in triplicates. Microcosms were incubated anaerobically for a 4-days while nitrate and nitrite were quantified colorimetrically. For metabolic dynamics, we measured nitrate (NO*^−^*) and nitrite (NO*^−^*) flux, the first two intermediates in denitrification and dissimilatory nitrate reduction to ammonium (DNRA). Communities were quantified by 16S rRNA amplicon sequencing before and after slurry incubation. **(B)** A subset of nitrate concentration dynamics (function) during the 4-day anaerobic incubation: three topsoils with different native pH levels (rows) were perturbed to either acidic or basic pH (columns) at the start of the incubation (T_0_), all with (CHL+, red) and without chloramphenicol (CHL-, black) treatments in triplicates (see Methods). The pH indicated inside the panels is the stabilized end-point pH to which the slurries were perturbed (Methods). **(C)** Schematic depicting three different functional regimes that capture how the soil community responds to pH perturbations. With moderate pH perturbations, the functional response can be characterized as the Nutrient-limiting regime (Regime II), where nitrate utilization is performed by dominant taxa (blue) that utilize nutrients released from the soil matrix due to perturbation. Growth is limited by the amount of available growth-limiting nutrients (purple). During strong basic perturbations, growth-limiting nutrients are in excess, and rare taxa (green) rapidly outgrow dominant populations that cannot perform nitrate reduction in basic conditions, hence the Resurgent growth regime (Regime III). Strong acidic perturbations induce cell death that limits metabolic activity, resulting in an inactive state (Acidic death regime, Regime I). Functional biomass of the dominant (blue) and rare (green) taxa are shown by the lines below.

For each soil sample, we created mixtures of soil and water (slurries) with 2mM nitrate and varying levels of strong acid or base to perturb each soil’s native pH to 13 values between 3 and 9 (Fig. 1A). Therefore, our experiment quantifies the effects of short-term pH perturbations, while the differences between soils can inform us about the effects of long-term exposure to high or low pH. We employed slurries to make amendments easier, limit the effects of differential water content, and mimic rain events when most of the anaerobic respiratory nitrate utilization occurs [31, 32].

Soil slurries retained much, but not all, of the complexity of the natural context, including the diversity of the communities, the soil nutrient composition, and the spatial structure due to intact soil grains. The metabolic activity we observed relied only on the natively available carbon (electron donor for nitrate reduction), and thus we did not enrich for specific taxa beyond the nitrate added to the system. To separate the activity of pre-existing nitrate utilizers from growth in each condition [33], we included controls in every pH perturbation treated with chloramphenicol which inhibits protein synthesis (Fig. 1A). The dataset comprised 20 soils, at 13 distinct pH levels, with and without chloramphenicol, each in triplicate.

We measured the dynamics of the relevant metabolites (nitrate, nitrite, ammonium, and watersoluble organic carbon) during the 4-day incubation in anaerobic conditions (Fig. 1A). Focusing on non-gaseous metabolites enabled us to perform high temporal resolution measurements of metabo-lite dynamics across the *∼*1500 microcosms. For 10 of 20 soils, we performed 16S rRNA amplicon sequencing before and after incubations.

We observed three types of dynamics across pH perturbations and soils (Fig. 1B). First, all chloramphenicol-treated (CHL+) conditions exhibited linear nitrate (NO*^−^*) utilization dynamics (red lines, Fig. 1B, Figs. S1, S2). This is expected because, with chloramphenicol, nitrate reducers are unable to grow, and the rate of nitrate reduction remains constant [31]. The slope of nitrate in time in CHL+ conditions quantifies the activity of the pre-existing functional biomass. For large acidic or basic perturbations, we observe little or no nitrate reduction in the CHL+ condition (flat red lines, far left/right columns, Fig. 1B) indicating that there is little pre-existing functional biomass that can reduce nitrate under large pH changes. Second, we observed linear nitrate/nitrite reduction dynamics even in samples without chloramphenicol (CHL-) for pH perturbations around the native pH (black lines, Fig. 1B, Figs. S2, S3). Thus, near the native pH, after some early growth, the functional biomass stays constant even without the growth-inhibiting drug (CHL-), suggesting that the growth of the functional biomass is limited by nutrients other than nitrate (schematic, Fig. 1C). Third, when we perturb the pH above 8, we observe an initial lag of nitrate reduction, followed by an exponential increase in reduction rate (black lines, far right, Fig. 1B). This indicates that an initially rare population grows rapidly reducing all available nitrate.

### Simple consumer-resource model captures metabolite dynamics across all pH perturbations

To describe the nitrate dynamics, we used the consumer-resource model presented in Fig. 2. Crucially, this model subsumes the ecological complexity of the soil microbiome into a single effective biomass rather than explicitly considering the multitude of possible interactions between taxa. The model has three variables: the functional nitrate-utilizing biomass (x), nitrate concentration (A), and the second growth-limiting nutrient (C) whose existence we hypothesized above. The five model parameters include: consumption rates (r*_A_* and r*_C_*), growth rate (γ), and affinities (K*_A_* and K*_C_*). The consumption rate of a resource is determined by the amount of functional biomass (x) and per-biomass consumption rates. The biomass growth rate (x^•^) is set to zero in CHL+ conditions due to chloramphenicol inhibition (γ = 0, x^•^ = 0).

**Figure 2:**
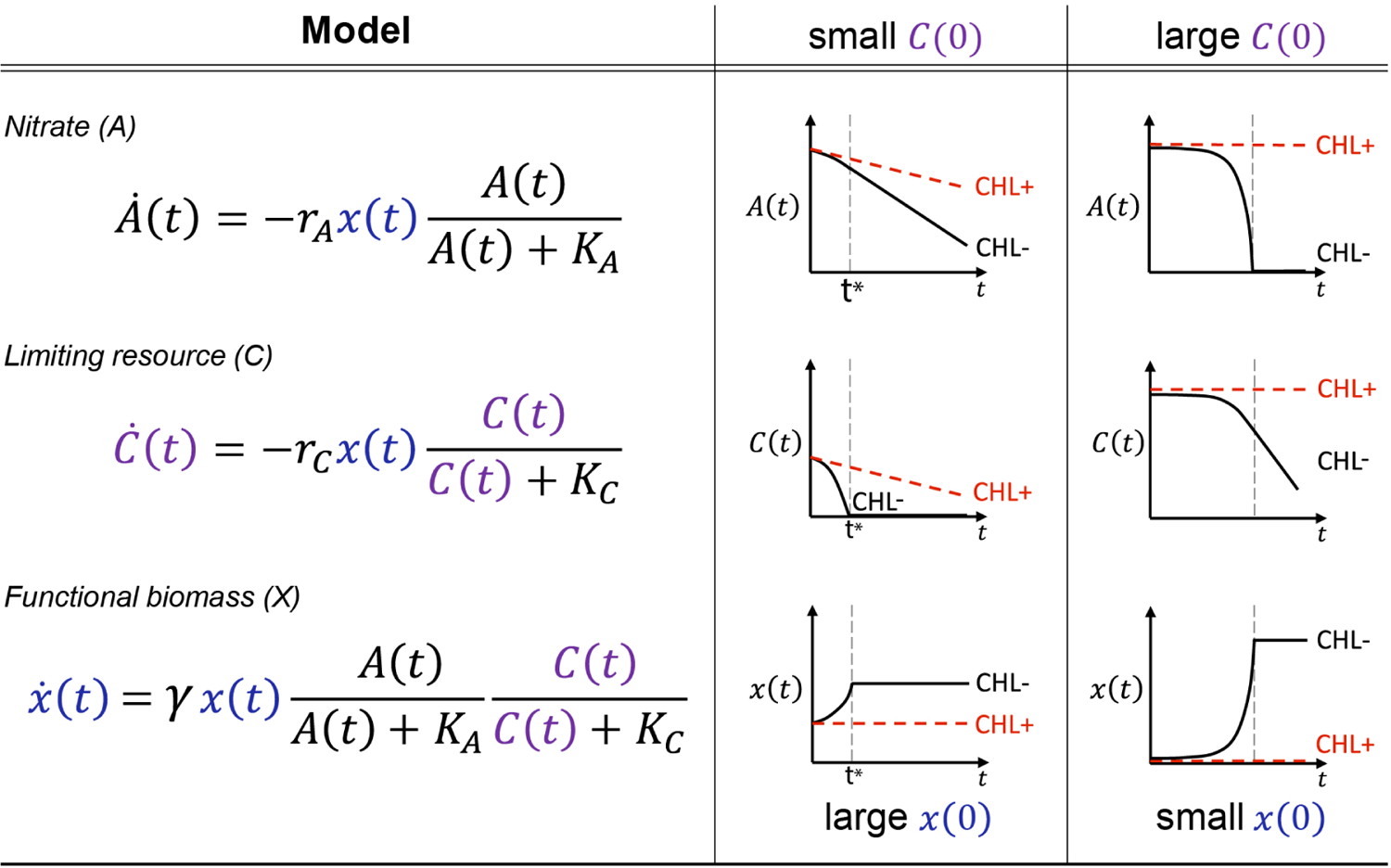
Consumer-resource model captures metabolite dynamics. A mathematical representation of the consumer-resource model to fit the nitrate reduction dynamics of the community (**Model** column). The model describes the community through the total functional biomass (*x*, biomass) which describes the aggregated biomass of species that perform nitrate reduction, nitrate concentration (*A*, mM), and a limiting resource concentration (*C*, mM). Nitrate consumption rate (*A*^•^(*t*)) takes a Monod [34] form with a reduction rate parameter (*r_A_*, mM/biomass/day) and an affinity parameter (*K_A_*, mM). Nitrite, which is reduced from nitrate, is not modeled. To capture linear nitrate dynamics, we include a non-substitutable resource that limits growth (*C*) with Monod consumption function and parameters *r_C_* (mM/biomass/day) and affinity parameter *K_C_* (mM). Growth of functional biomass (*x*•(*t*)) is determined by concentrations of nitrate (*A*) and limiting nutrient (*C*) with biomass growth rate (*γ*, 1/day). Plots in the right two columns show dynamics of *x*(*t*), *A*(*t*), and *C*(*t*) at small *C*(0) and large *x*(0) (middle) and large *C*(0) with small *x*(0) (right). Red and black traces show dynamics with and without growth-inhibiting (*x*•(*t*) = 0) chloramphenicol respectively. Without growth, the nitrate reduction rate is constant and proportional to the functional biomass *x*(0) (red lines, top row for large/small *x*(0)). The **small C(0)** column illustrates how the model captures linear nitrate dynamics in chloramphenicol untreated (CHL-) conditions (black lines). With the small amount of initial limiting resource *C*(0), functional biomass will stay constant after the limiting nutrient is depleted at t*^∗^*, which produces a constant nitrate reduction rate (linear NO*^−^* dynamics, black line, top). The **large C(0)** column shows exponential nitrate depletion dynamics in CHL- conditions (black lines) when there is excess *C*(0) and *x*(0) is small. Functional biomass grows exponentially, resulting in exponential nitrate utilization dynamics (black line, top). The affinity parameters (*K_A_*, *K_C_*) and yield parameter (*γ*) were fixed for all samples (see Methods for rationale and Fig. S8).

If the initial nutrient concentration C(0) is small (Fig. 2, middle column), the nutrient C runs out quickly, arresting biomass growth and resulting in A being consumed at a constant rate from t*^∗^* onwards (dashed line). This recapitulates the late-time linear dynamics in CHLconditions for moderate pH perturbations(Fig. 1B). In contrast, when the initial nutrient concentration C(0) is large (Fig. 2, right column), it is nitrate (A) that runs out first. In this regime, the initial rate of nitrate utilization (determined by x(0), the initial functional biomass) grows exponentially until A runs out. Therefore, a small x(0) and a large C(0) recapitulates the initially slow but exponentially growing dynamics observed for large basic pH perturbations (Fig. 1B).

Our consumer-resource model provided a good fit to the observed nitrate dynamics in all soils (<10% error per data point, Fig. S7). To perform this fitting, we fixed the growth rate γ and the affinity parameters (K*_A_*, K*_C_*), and varied just two rescaled parameters: x^^^(0) = x(0)r*_A_* and γC^^^(0) = γC(0)r*_A_*/r*_C_* (see Methods). These parameters retain the same interpretation: x^(0) reflects the initial functional biomass, and γC^^^(0) the available limiting nutrient, the rescaling corresponds to measuring these quantities in terms of nitrate utilization rates (see Methods).

### Model reveals functional regimes

We plotted x^(0) (pre-existing functional biomass) against γC^^^(0) (available limiting nutrient, Fig. 3A) and identified three regimes of nitrate utilization dynamics (Methods, Fig. S9). Regime I, the Acidic death regime (both x^(0) and γC^^^(0) are low) is observed for pH ≲ 4, and shows little to no nitrate reduction (Fig. 3B, (a) and (d)). Regime II, the Nutrient-limiting regime (x^(0) is large and γC^^^(0) is small) is observed for 4 ≲ pH ≲ 8, and exhibits a relatively large initial nitrate reduction rate that transiently increases and then remains constant (Fig. 3B, (b) and (e)). Finally, Regime III, the Resurgent growth regime (small x^(0), large γC^^^(0)) is observed for pH ≲ 8, and displays a closeto-zero initial utilization rate, followed by an exponential speed-up that continues until nitrate is depleted (Fig. 3B, (c) and (f)).

**Figure 3:**
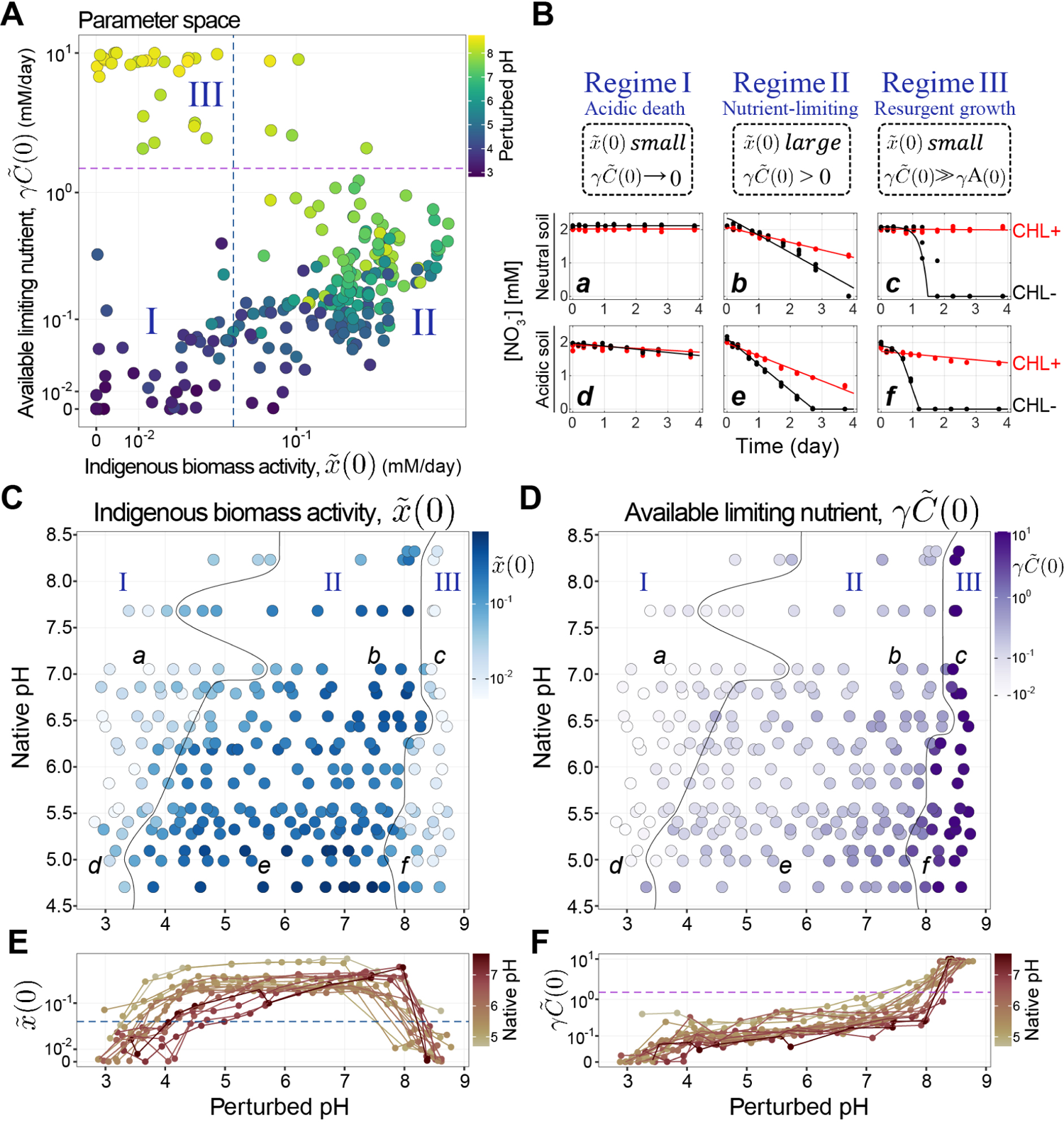
Conserved regimes capture soil’s functional response to pH perturbations. **(A)** Scatterplot of the two model parameters (functional biomass *x*^(0), limiting nutrient concentration *γC*^^^(0)) inferred from nitrate dynamics across all samples. See text and Methods for details of model fitting. Note log-scale. The color of points indicates each sample’s perturbed pH. Three regions separated by dashed lines indicate the distinct regimes of functional response against pH perturbations: the Acidic death regime (Regime I), Nutrient-limiting regime (Regime II), and Resurgent growth regime (Regime III). The locations of the dashed lines were determined by thresholding distributions of *x*^(0) and *γC*^^^(0) (see Methods). **(B)** Example nitrate dynamics for each of the three regimes for a neutral soil (top row) and an acidic soil (bottom row). Red lines are with growth-arresting chloramphenicol and black without. (*a, b, c, d, e, f* correspond to perturbed conditions indicated in panels **C and D**). *a,d* show little nitrate reduction, *b,e* show linear nitrate dynamics with slopes that increase without chloramphenicol (see Fig. 2, middle column), and *c,f* show no activity without growth (red) but exponential nitrate utilization in the absence of the drug (Fig. 2, right column). **(C) and (D)** pH affects indigenous biomass activity *x*^(0) (blue) and available limiting nutrient *γC*^^^(0) (purple). Fitted parameter values are shown with the color (log-scale) in the grid of long-term pH variation (y-axis, Native pH) and short-term pH perturbation (x-axis, Perturbed pH). In all soils from different native pH levels, we observe a conserved set of responses to short-term pH perturbations: Nutrient-limiting regime (region indicated by II) near the native pH, then transitioning to the Acidic death regime (region indicated by I) during acidic perturbation (black line), also transitioning to the Resurgent growth regime (III) for basic perturbations (black line). Long-term (native) pH dictates the pH thresholds of regime boundaries (black line). **(E)** Trends of *x*^(0) (log-scale) across varying perturbed pH values for soils with different native pH levels (native pH indicated by line color), demonstrating consistent transition between regimes and a plateau of high activity within the mid-range pH (Regime II) across all soils. **(F)** *γC*^^^(0) (log-scale) with perturbed pH, showing a rise in limiting nutrients induced by short-term pH increases. Colors indicate native pH. We used the median fitted value of the three biological replicates for all data points of *x*^(0) and *γC*^^^(0).

We observe all three functional regimes across all soils, but the pH at which a transition from one regime to another occurs depends on the native pH of the soil. Figure 3C&D shows the inferred initial functional biomass (x^(0)) and limiting nutrient (γC^^^(0)) across soils of varying native pH (yaxis) and laboratory perturbed pH (x-axis). We next harnessed our model to identify mechanisms underlying these regimes.

### Metabolite dynamics in Regime II are governed by carbon release

In Regime II (the Nutrient-limiting regime), the nitrate reduction rate increases with pH (Fig. 1B). Our model proposes that the mechanism behind this increase is the increasing availability of the growth-limiting nutrient (Fig. 3F), which translates into larger growth of active biomass and hence the increased nitrate reduction rate (Fig. 2). Here, we investigate whether this model prediction is valid by examining how increasing pH leads to higher levels of growth-limiting nutrients and identifying these nutrients.

Previous studies have observed that increasing pH can enhance the availability of organic carbon in soils [35–37]. Studies indicate that this release of nutrients from soil is based on a substitution mechanism at the ion exchange sites within the soil clay particles [38, 39] (Fig. 4B, detailed mechanism in SM). Therefore, we hypothesized that the amount of nutrients released would be proportional to the quantity of either base (NaOH) or acid (HCl) added to the slurry. Based on this assumption, the fold change in nitrate reduction rate, reflecting the growth of active biomass limited by this nutrient, should be proportional to the quantity of acid or base added to the system. In Fig. 4A, we observe precisely this trend across all soils, as evidenced by a data collapse in the increase in nitrate reduction rate with NaOH (light blue region). The trend is specific to Regime II (Fig. S11A), and if the data are plotted against pH, the correlation becomes much weaker (Fig. S10). As further evidence supporting our hypothesis, we measured increases in the absolute abundances via 16S rRNA amplicon sequencing with internal standards (see Methods). Increasing sequencing reads reflect increases in biomass, both at a coarse level (fold change in total biomass, Fig. S11B) and fine level (individual ASVs that responded to the amendment of nitrate, Fig. S11C, see SM for details). Corroborating our hypothesis, we observe a linear relationship between the increase in absolute abundances and the amount of NaOH added to the system (Fig. S11).

**Figure 4:**
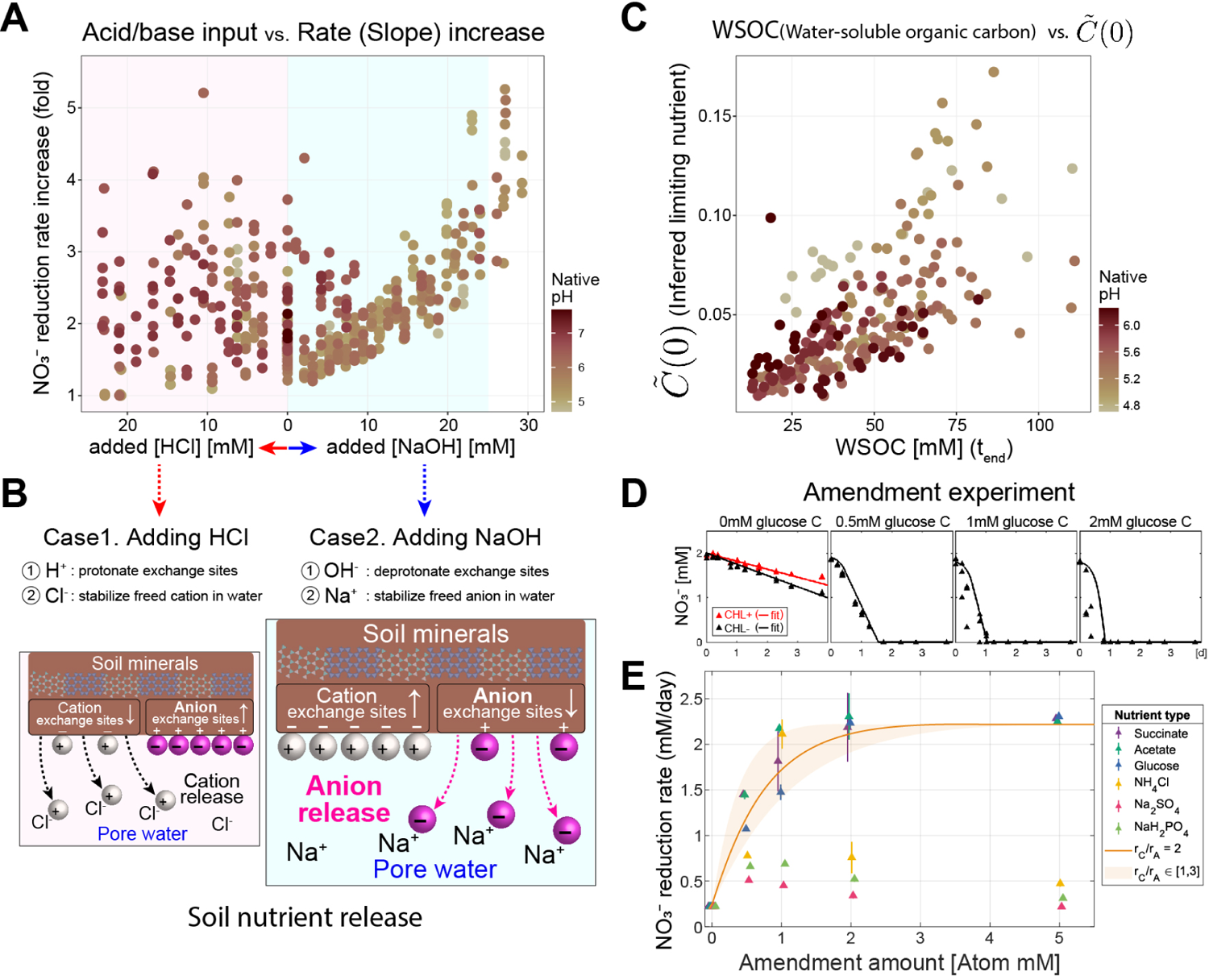
Carbon limits growth in Regime II and is released by ion exchange mechanism. **(A)** The amount of NaOH added to the soil plotted against the fold-change of nitrate reduction rate (ratio of rate with growth (CHL-) and with no growth (CHL+)). Base additions from 0mM to 25mM NaOH correspond to soils in the Nutrient-limiting regime (Regime II, Fig. 3) (light blue background in **A**). No increase in growth was observed for acidic perturbations (*>*0 mM HCl addition, pink region, Fig. S11A). **(B)** Cartoon illustrating the mechanism of soil nutrient release hypothesis; NaOH results in the release of anionic nutrients (magenta-colored spheres) from soil particles (brown region), while the addition of HCl would release cationic nutrients (white spheres) and adsorbs anionic nutrients. Microbes cannot access the nutrients adsorbed in the soil particles but can access the nutrients dissolved in pore water. Added OH*^−^* ions decrease the number of anion exchange sites in the soil particles, releasing anionic nutrients. In concert, Na^+^ ions stabilize the released anions (see text and Fig. S13B for additional details). **A** and **B** suggest the growth-limiting nutrients are anionic (negatively charged). **(C)** Scatterplot of model-inferred *C*^^^(0) (available limiting nutrient) and measured water-soluble organic carbon (WSOC) measured via a chromate oxidation assay (Methods) that is not chemically specific and WSOC likely contains different C compounds, N, P, etc. Data points are from Soil 1–12 samples where there are enough number of points per soil to observe a linear relationship in the light blue region in **A** (0mM–25mM NaOH,). **(D) and (E)** Amendment experiments for soil in the Nutrient-limiting regime (Regime II) at unperturbed pH. **(D)** Panels show nitrate dynamics with different levels of glucose amendments (red: CHL+, black: CHL-, points: data, lines: model fit), where linear dynamics (at 0mM C) transition into exponential dynamics (*≥* 0.5mM C) supports carbon limitation of nitrate utilization. Lines are model predictions. **(E)** Nitrate reduction rates after amending soils with different concentrations of nutrients (three carbon sources, ammonium, sulfate, and phosphate). Points are the mean rates, estimated by linear regression, of triplicates with error bars indicating standard deviation. Carbon (succinate (pK_a_ = 4.2 and 5.6), acetate(pK_a_ = 4.75), and glucose) amendments increased the nitrate reduction rates starting from low concentrations (0.5 CmM)). Carbon compounds are negatively charged when pK_a_ < pH (here, the pH of soil 6 is 5.4). Ammonium, sulfate, and phosphate did not result in a similar increase in nitrate reduction. We cannot independently infer the ratio r*_A_*/r*_C_* (Methods), model predictions are shown for 1 < r*_A_*/r*_C_* < 3 (shaded region) with a line for r*_A_*/r*_C_* = 2 (best fit). This ratio can be interpreted as the nitrate:carbon utilization ratio.

The asymmetric response of the change in nitrate reduction rate upon the addition of NaOH rather than HCl (blue versus pink shaded regions Fig. 4A) provides insight into the identity of the released nutrient. Under the mechanism of ion-exchange-mediated nutrient release, adding ions releases nutrients adsorbed to the clay particles into the pore water, making them accessible to microbes (Fig. 4B). HCl and NaOH will release cationic and anionic nutrients respectively (SM for details, Fig. S13B). Our observation that the limiting nutrient governing Regime II dynamics is released in proportion to the amount of NaOH indicates that the growth-limiting nutrient is anionic, with likely candidates including phosphates, sulfate, or carbon. Notably, measurements of watersoluble organic carbon (WSOC) at the endpoint increased linearly with NaOH added (Fig. 4C). This suggests that some WSOC is negatively charged (anionic) and that the growth-limiting nutrient might be WSOC, or concomitantly released nitrogen (N), sulfur (S), or phosphorus (P).

To further identify the limiting anionic nutrient, we performed an amendment experiment on a representative soil (Soil 6 (pH 5.4), see Methods). We amended a soil slurry without perturbing pH with glucose (neutral), succinate (anion when pH > pK_a_ = 4.2), acetate (anion when pH > pK_a_ = 4.75), phosphate (anion), ammonium (cation), and sulfate (anion) added in varying concentrations (Methods, Fig. S14). We found that the amendment of carbon, but not other N, S, and P sources, immediately increased the nitrate reduction rate, changing the linear dynamics to exponential (Fig. 4D & E), indicating that carbon was the limiting nutrient. With a single free parameter, our model predicted the nitrate utilization dynamics in a soil amended with glucose (Fig. 4D). Similar results are found for other carbon sources, but not sources of N, S, or P (Fig. 4E). The single free parameter is the ratio r*_C_*/r*_A_*, which can be interpreted as a stoichiometry of carbon to nitrate utilization (Fig. 2). We find this ratio to be highest for glucose (2.5) and lowest for acetate (1), suggesting carbon is utilized more quickly relative to nitrate in glucose amendments. The relatively more rapid utilization of glucose may be because glucose can be consumed by anaerobic respiration (requiring nitrate) and fermentation, whereas acetate is not fermentable. The amendment experiment confirms the mechanism predicted by our model, that a nutrient other than nitrate limits reduction dynamics for modest pH perturbations. Critically, this insight emerged naturally from our mathematical description of the nitrate utilization dynamics across pH perturbations.

### Regime III arises from the rapid growth of rare taxa

Under large basic pH perturbations, all soils exhibited a sharp transition from linear to exponential nitrate consumption dynamics. Our model fits suggest interpreting these metabolite dynamics as resulting from a small initial functional biomass (x^(0)) undergoing exponential growth due to excess nutrient γC^^^(0) (Fig. 3C-F). To test this interpretation, we used the sequencing data to investigate the compositional changes that occur after large basic perturbations (Regime III).

Sequencing measurements corroborate our model predictions by revealing a group of rare taxa enriched in Regime III. These are especially clear if ASVs are grouped at the phylum level, revealing that Firmicutes undergo explosive growth in this regime (10-fold enrichment at the aggregate phylum level, and several hundred-fold for individual ASVs, particularly in the Bacilli genus; Fig. S17). We computed the fold change of each phylum’s absolute abundance across treatments relative to the no-growth CHL+ control. Non-negative matrix factorization (NMF) analysis of the growth fold values revealed that most of the variation in these data could be captured with just two axes of variation (Fig. S15B, Methods). Each of these axes was composed of one or two phyla, one included Proteobacteria and Bacteroidota, and the other Firmicutes.

Fig. 5A-B shows growth-folds for the two groups of phyla identified by NMF that dominate growth across all soils and pH conditions. In the Nutrient-limiting regime (Regime II), Proteobacteria and Bacteroidota increased their growth with increased pH, then decreased towards the start of Regime III. This matches the growth behavior of indigenous functional biomass (x^(0)) revealed by the model in Regime II (Fig. 3C). Conversely, Firmicutes did not grow until a critical pH threshold between 7-8.5, which matches the onset of exponential nitrate utilization dynamics in Regime III (Fig. 3F). Importantly, the boundary between Regime II and III derived from the functional dynamics data (Fig. 3 C & D), aligns with the shifts in growth responses of Firmicutes (Fig. 5B) and Proteobacteria/Bacteroidota (Fig. 5A). These growth patterns suggest that the changes in the identity of the phyla responsible for nitrate reduction reflect the functional regimes. A more detailed analysis of the likely metabolic traits of these strains [40] suggests that the transition from Regime II to III is also accompanied by a shift from denitrification to DNRA which agrees with the fact that excess carbon favors DNRA [41](Figs. S18 S19).

**Figure 5:**
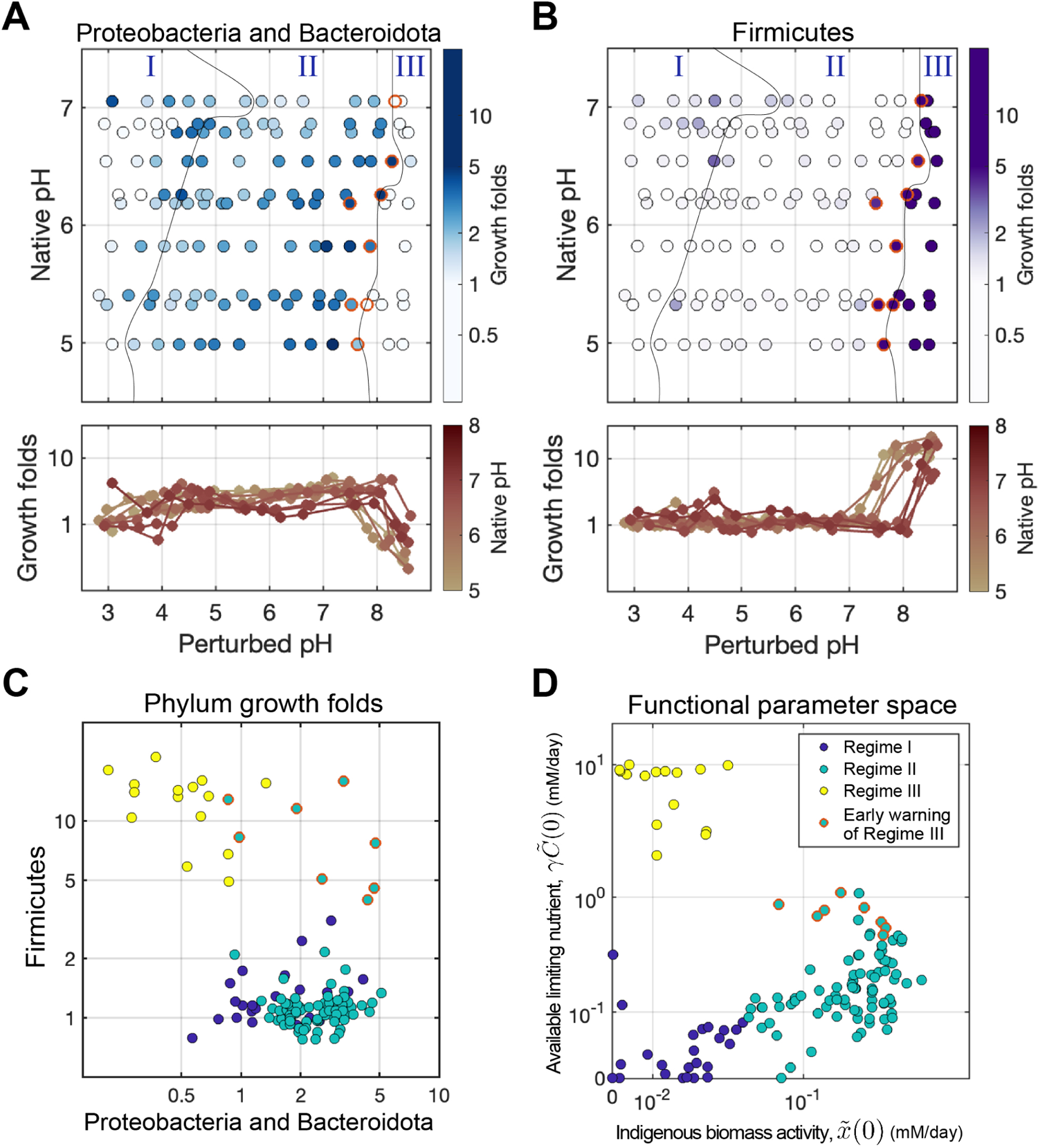
Regime III: Resurgent growth emerges from native population decline and rare taxa expansion. Global trends of growth in the phylum level across perturbed pH levels reveal taxonomic origins of the rapid growth in the resurgent growth regime (Regime III). 16S amplicon sequencing at the end of each incubation was used to identify amplicon sequence variants (ASVs) in CHL+/conditions (Methods). ASVs were aggregated at the phylum level. For each phylum, a growth fold was computed as the ratio of abundances with/without growth (Abs*_CHL−_/*Abs*_CHL_*_+_). A statistical decomposition across all conditions identified three phyla that dominated abundance changes due to growth: Proteobacteria and Bacteriodota with similar changes, and Firmicutes. (see main text and Fig. S15). **(A)** Growth folds for the phyla Proteobacteria + Bacteroidota (combined abundance) indicated by the color for each native and perturbed pH condition. The growth declines at a basic pH threshold, mirroring the patterns observed in the fitted model parameter x^(0) (indigenous biomass activity, Fig 3C) at the Regime II-III boundary). Regime boundary lines are those determined in Fig. 3). Line plots (lower panel) growth folds were plotted in log-scale, color indicating native pH given in color bar. **(B)** Identical to (A) but showing growth folds (color) of Firmicutes increasing during the transition from Regime II to III, where pH perturbations are strongly basic. This mirrors the increase in inferred carbon concentrations γC^^^(0) (Fig. 3). **(C)** Scatter plot of growth folds of Proteobacteria + Bacteroidota against Firmicutes. Points marked in red, associated with Regime II (also red in (D)), exhibit high growth of both Proteobacteria + Bacteroidota and Firmicutes. For red points, Firmicutes abundances are an early-warning indicator of a transition between regimes. **(D)** Same plot as Fig. 3A of x^(0) verses γC^^^(0) with points marked by the regime they belong to and red points indicating Regime II conditions near the boundary between Regime II and III. Note these red circles are in Regime II, but the Firmicutes abundances are high (panel **(C)**).

### Growth is an early-warning indicator of a transition between regimes

Intriguingly, we found that Firmicutes begin increasing at pH levels just below the transition from Regime II to III, thereby acting as ‘early warning indicators’ for the impending transition (red circles, Fig. 5D). Specifically, when we plot the growth folds of the Firmicutes versus Proteobacteria and Bacteriodota, we find that Firmicutes abundances begin to rise prior to the system entering Regime III as defined by nitrate utilization dynamics (Fig. 5C). This finding indicates that compositional data can be used to predict impending functional state transitions during environmental perturbations.

### Acidic perturbations in Regime I reduce functional biomass via death

In response to a short-term decreases in pH, the model indicates a reduction in indigenous functional biomass (x^(0)) and a decrease in the availability of limiting nutrients (Fig. 3). Below a pH value of 3–5, depending on the soil’s native pH, nitrate reduction ceases (Regime I). We tested whether the sequencing data reflects the decreasing trend of functional biomass (x^(0)) with pH. We computed the fold-change in each Phylum’s endpoint absolute abundance in CHL+ conditions relative to abundances at the initial time point T_0_ (‘survival fold’; Fig. S20A). This ratio reflects the change in abundance in the absence of growth, hence we regard this as a proxy for death. For all phyla except the Firmicutes, we observed a consistent drop of survival folds during acidic perturbations (Fig. S20A). Furthermore, we confirmed that the survival folds exhibited an approximately linear relationship with the x^(0) values (Fig. S20B). These observations confirm the decline of biomass in acidic conditions, likely via death and DNA degradation, except in taxa tolerant to short-term pH changes (Firmicutes, Fig. S20A). Thus, we conclude that acidic perturbations lead to widespread death, while basic conditions lead to selective growth (Fig. S15C).

### Long-term soil pH defines regime boundaries

Next, we sought to understand what properties determine the pH at which soils transition between regimes. We observed that the native pH of the soil (long-term pH) determined the pH at which any given soil transitioned between functional regimes (Fig. 3C & D). This result suggests that the soil communities are adapted to their long-term pH conditions [19, 33, 42].

One key property of soils that impacts the pH variation the microbiome experiences is the soil’s pH titration curve: how soil pH changes in response to acid/base additions. The shape of the titration curve was similar across all soils (Fig. 6D, Fig. S23A), showing a plateau at low and high pH with a nonlinearity in between. As a result, acidic soils with native pH near the lower plateau were more strongly pH-buffered than the neutral soils (with native pH around the steepest portion of the nonlinearity; Fig. 6B, Fig. S23A). This observation indicates that at similar levels of acid addition, neutral soils would experience *a larger drop in pH* than acidic soils (ΔpH*^acidic^* < ΔpH*^neutral^*, Fig. 6A, B). We speculate that this makes communities in acidic soils less tolerant of acidic pH fluctuations, as they are less likely to experience large reductions in pH. This reasoning would help explain the observation that acidic soils transition from regime II to I after a smaller perturbation in pH than neutral soils (Fig. 6E, ΔpH*^acidic^* < ΔpH*^neutral^*). As a result, plotting the pH at the regime II to I transition against the native pH gives a line with slope <1 (Fig. 6E, bottom dashed line), where a line of slope 1 would indicate that entry into Regime I requires an acidic pH shift of a constant magnitude.

**Figure 6:**
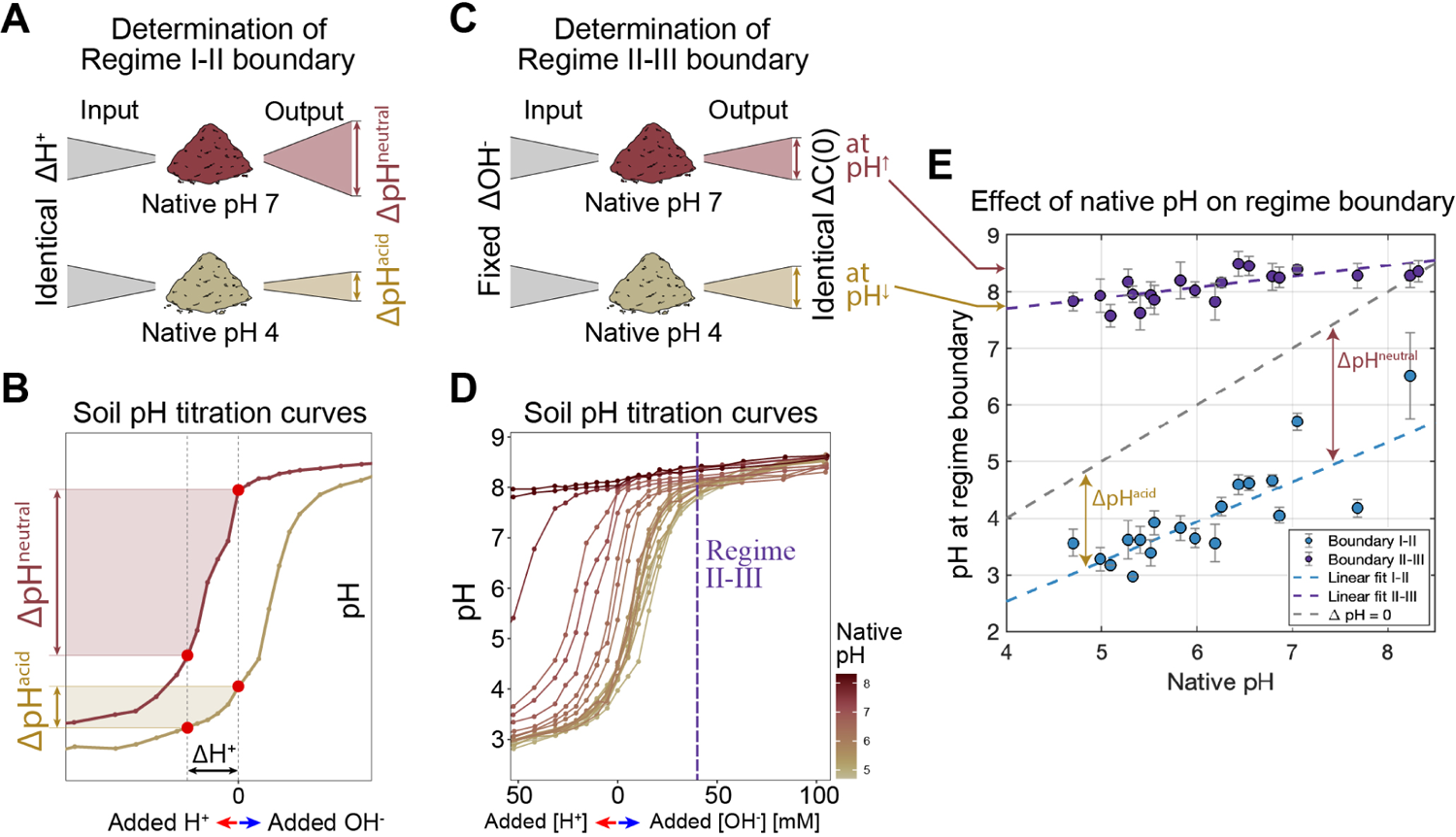
Regime boundaries are determined by long-term pH and history of pH variation. **(A)** Cartoon for how native pH impacts the pH of the Regime I and II boundary. The cartoon depicts how the identical amount of acid perturbations (ΔH^+^) gives rise to larger changes in pH for neutral soils (ΔpH*^neutral^*) than acidic soils (ΔpH*^acidic^*) which arises due to differences in the location of the native pH on the titration curve shown in **B**. **(B)** Shows titration curves where the pH (y-axis) is measured after adding different amounts of acid or base (Methods) for neutral soil (dark brown) and acidic soil (light brown). The dashed vertical line at 0 indicates the pH with no acid/base perturbation. Due to the shape of these curves, if both soils are subjected to the same ΔH^+^ (bottom) the neutral soil experiences a larger change in pH (shaded regions). This suggests that acidic soils experience smaller pH fluctuations and therefore transition to Regime I from II after smaller pH perturbations as shown in (**E**). **(C)** Cartoon for how native pH determines the pH of the Regime II and III boundary. The cartoon depicts how the fixed amount of added base (NaOH) results in an identical amount of released carbon (Δ*C*(0), Fig. 4). Large *C*(0) drives the transition from Regime II to III. For a fixed addition of NaOH, more neutral soils reach higher pH again due to the shape of the titration curves as shown in **D**.). **(D)** Soil pH titration curves (identical to **C**) for all soils with different native pH levels. The vertical dashed line indicates the quantity of NaOH added to move from Regime II-III. More neutral soils (darker colors) reach higher pH values for this fixed quantity of added NaOH. This correlates with the increasing pH at the Regime II-III boundary (purple points, (E)). **(E)** pH levels (y-axis) when transitions between functional regimes occur from Regime II to I (blue points) and from Regime II to III (purple points) for soils from different native pH levels (x-axis). Regime boundaries are determined as the midpoint between the last pH perturbation in Regime I and the first in Regime II. Error bars represent the pH difference between these conditions. An identical strategy was used for Regime II-III. The dashed blue line (Regime I-II boundary) and dashed purple line (Regime II-III boundary) are weighted least squares fits, with the weights inversely proportional to the error of each point. The dashed black line is slope 1, where the change in pH from native to the regime boundary is constant for all soils. Lines with a slope different from 1 indicate that the difference between native pH and pH at the regime boundary depends on the native pH of the soil. The slope of the blue dashed line is 0.7 (95% confidence interval: [0.44, 0.97].

In contrast, we find that soils transition from Regime II to III when carbon is in excess. From Fig. 4, we know that carbon is released in proportion to the NaOH added to the slurry (Fig. 6C). Accordingly, we find that a constant addition of NaOH drives the transition from Regime III to II (Fig. S24). However, due to the shape of the titration curves as seen in Fig. 6D, for a constant base amendment, more neutral soils reach higher pH (dashed line Fig. 6E). Therefore, as expected from the titration curves, more basic soils transition to Regime III at higher pH (Fig. 6E).

Our sequencing data support the idea that variation in regime boundaries with native pH has a basis in the taxonomic composition of the microbiome. In more acidic soils, the Proteobacteria and Bacteroidota show better survival at lower pH (Fig.S21). In contrast, the pH at which Fermicutes begin to grow in Regime III rises with the soil’s native pH (Fig. 5B), in line with the Regime II to III transition observed in functional measurements (Fig.6A). In addition, the native pH of the soil is predictable from the identity of the strains that exhibit growth in regime III (Fig. S17A, S22). These findings suggest that prolonged exposure to a specific pH likely selects for specific taxa, thereby influencing the pH at which the community transitions between functional regimes.

## Discussion

We showed that a simple mathematical model derived from quantitative measurements of metabolite fluxes delineates which mechanisms are relevant for understanding the functional response of the soil to perturbations. Remarkably, the model does not attempt to account for all processes in the soils and instead captures the behavior of the entire community using a single effective biomass subjected to nutrient limitation. From this perspective, we identified functional regimes demarcated by whether active biomass or available nutrients dictate the metabolism of the system. For example, we discovered a nutrient-limiting regime (Regime II), where the indigenous biomass is robust to moderate pH changes [43] and metabolism is governed by carbon limitation. In contrast, metabolism in Regime III is governed by the growth of initially rare taxa perturbations [44].

### Limitations of the study

Our study has several limitations. First, our soil slurries do not capture the full complexity of natural soils. Microcosms experience fixed anaerobic conditions, but nitrate utilization in the wild occurs during fluctuations between aerobic and anaerobic conditions [45]. Second, our more extreme pH perturbations (ΔpH > 2) are larger than is routinely experienced in natural systems. Third, unlike previous studies, we do not quantify nitrous oxide and nitrogen gas (Table S2) production both downstream products of denitrification. Nitrous oxide is of critical interest given its importance as a greenhouse gas, so it will be important to understand how its production varies across functional regimes.

Finally, the simplicity of our model, which describes a single effective biomass, leaves open the question of what role ecological interactions play in determining community metabolism. It is unclear within a given regime whether there are strong interactions between responding taxa or not. At the transition between regimes II and III, we cannot determine if the Firmicutes outcompete the Regime II taxa for carbon or nitrate or whether the physiology of taxa that dominate in Regime II does not permit their growth in more alkaline conditions.

### The significance of functional regimes and their generalizability

Our study establishes the existence of functional regimes where specific chemical, physiological, or ecological processes govern system response. This demonstrates that understanding the community response to perturbation may not require grappling with every metabolic process or interaction in the community, but only with a handful of key features. Our demonstration comes in the context of nitrate utilization and soil pH. However, this study opens the door to asking whether similar functional regimes describe community response to a suite of key perturbations including temperature or xenobiotics. A previous study of the response of soils to temperature revealed dynamics strikingly similar to Regime III at high temperatures and the asymmetric response in the Acidic death regime (Regime I) at low temperatures [46].

### Functional regimes as guides for understanding complex omics data

Sequencing measurements of complex microbiomes result in datasets with thousands of variables taxa, genes, or transcripts. Distilling some understanding from these data presents a huge challenge. The existence of regimes guided our understanding of the dynamics of the >30,000 ASVs in our dataset by directing us to look for specific responses.

More broadly, the last decade has seen an explosion of methods for quantifying community dynamics and metabolism from transcriptomic and metagenomic measurements [47–49] to single-cell metabolomics [50] and quantitative stable isotope probes [51–53]. The challenge is to synthesize these data to achieve insights into dynamics and function. Our work illustrates the promise of an approach where we acquire large-scale quantitative measurements of metabolism at the whole community level, describe these dynamics mathematically, either phenomenologically or potentially new AI-driven methods [54], and then interpret the resulting model mechanistically. For example, in Regime III, we expect native taxa to exhibit stress response and declining metabolic activity, and the converse for Firmicutes. Thus the framework of regimes suggests a route for leveraging new technologies for a deeper understanding of mechanisms in complex microbiomes.

### Physiological insights from constant utilization rates in nutrient-limited environments

The linear dynamics of nitrate utilization observed in Regime II have been previously observed [11, 31, 55–57] and attributed to carbon limitation [31]. Moreover, previous work supports the result that available organic carbon can be the limiting factor for nitrate utilization [10, 36, 55, 58–61]. How can limited carbon lead to a constant rate of nitrate reduction? Carbon is the electron donor for anaerobic respiration of nitrate which is the terminal electron acceptor. If carbon runs out we might expect that cells will run out of reductant to convert nitrate to nitrite, but this is not what we observe. One hypothesis is that cells internally store carbon to regenerate reductant [62]. To test this hypothesis, we incubated individual denitrifying bacterial strains in minimal media supplemented with 2mM NO*^−^* in the absence of exogenous carbon. Similar to the linear dynamics observed in the soil microcosm, we observed linear nitrate reduction dynamics in the carbon-limited monocultures (Fig. S6), revealing that the metabolism of a single strain can mirror the metabolism of the soil microbiome. The energy obtained from a constant rate of nitrate reduction is likely channeled to maintenance rather than growth. More broadly, the discovery of the three functional regimes, including the nutrient-limiting regime, is notable because it reflects a potential duality between the physiology of an ecosystem and the three phases of a cell: growth (Regime III), stationary (Regime II), and stress (Regime I). This duality suggests the possibility that cellular physiology might provide a conceptual framework for understanding the ecosystem.

### Unifying decades of prior work in a quantitative framework

To limit confounding factors, our study focused on 20 samples from a single agricultural site. However, the features of the three functional regimes defined here are present in many previous studies performed on soils from other sites, suggesting that these regimes are a general feature of nitrate utilization and pH perturbations. For example, Nömmik observed metabolite dynamics consistent with a transition from Regime II to III [63]. Parkin *et al.* observed a native pH dependent x^(0) with chloramphenicol applied [33], consistent with all three regimes. Simek observed increasing nitrate utilization rates with time as pH increased, another Regime II to III transition [64]. Anderson *et al.* observed increasing rates of nitrate utilization with increasing pH, and the recruitment of Firmicutes in very basic conditions [35]. These results show that the regimes are potentially general and not specific to our study site or protocol.

### Direct and indirect effects of pH perturbations

In the context of community metabolism, it has been debated whether the indirect effect of pH on nutrient availability is as important as the direct effect of pH on microbial physiology [17]. Our results answer this question because the model enables us to quantify both the pH’s indirect effect on the growth-limiting nutrient (changes in C^^^(0)) and its direct effect on indigenous functional activity that reflects physiology (changes in x^(0)). In the Nutrient-limiting regime (Regime II), the indirect pH effect (changes in C^^^(0)) is more important in determining the nitrate reduction rate because the indigenous functional biomass (x^(0)) is stable in this regime. In Regime III the model suggests physiological responses might be most important since nutrient limitation is relieved but only a small fraction of taxa grow.

### Optimal pH and long-term adaptation

Due to the agricultural importance of nitrate utilization, it has been debated whether soils exhibit an optimal pH for denitrification [64]. Previous studies have demonstrated the pH level associated with the highest rate of denitrification closely aligns with the native soil pH [33, 64] over short timescales (<3 hours, Table S2). Other studies observed a shift of optimum pH to more neutral values on longer timescales [64]. Our study reconciles these outcomes and elucidates the underlying cause. The fastest nitrate utilization occurs near the native pH of the soil on short timescales (Fig. 1B). This is consistent with our results because the pre-existing functional biomass (x^(0)) is the largest near the native pH (Fig. 3C&E). However, basic pH perturbations release carbon (C(0)), driving growth and faster reduction in alkaline conditions. Furthermore, with pH perturbations over 8 and long enough timescales (>12 hours), the growth of rare taxa drives fast nitrate reduction. As a result, the optimal pH depends on the timescale of the measurement.

### Functional regimes and environmental fluctuations: origins of microbial diversity in nature

For large basic perturbations, the abundant native taxa could no longer perform nitrate reduction, while the rare biosphere grew rapidly to reduce nitrate (Regime III), acting as the source of functional resilience in the community. The adaptation of rare taxa in extreme environments suggests that there might be a trade-off between stress resistance and fast growth [65]. Rare taxa may specialize in surviving under extreme stress conditions (e.g., Firmicutes, Fig. S20), but perform little metabolic activity when the environment is near its native state. Conversely, dominant taxa near native environmental conditions (e.g., Proteobacteria) may specialize in faster growth rates when the nutrient becomes available but have limited ability to persist in stressful environments. These observations give rise to a picture where rare taxa are sustained by the presence of environmental fluctuations that transiently provide an opportunity to exploit resources [66].

Soil pH can change daily due to plant exudates (shifts of 0.4 in 12 hrs) [67], seasonally due to changes in rainfall and temperature (1–1.5 pH units) [68], and through agricultural practices [69]. The titration curves gave insights into the amplitude of pH fluctuations the community experiences in nature (Fig. 6B). These observations place taxa’s distinct physiology and environmental fluctuations at the center of understanding the origin and structure of regimes and therefore the metabolism of natural microbial communities. While physiology has experienced a renaissance of late, with quantitative approaches providing key insights [70], we know comparatively little about the role of natural environmental fluctuations in the wild. Our results suggest that understanding the dynamics and origins of these fluctuations could provide deep insights into the responses of complex communities to environmental change.

## Methods

### Sample collection, site description, and soil characterization

Twenty topsoils were sampled across a range of pH values (4.7–8.32) from the Cook Agronomy Farm (Table S3). The Cook Agronomy Farm (CAF, 46.78°N, 117.09°W, 800 m above sea level) is a long-term agricultural research site located in Pullman, Washington, USA. CAF was established in 1998 as part of the Long-Term Agroecosystem Research (LTAR) network supported by the United States Department of Agriculture. Before being converted to an agricultural field, the site was zonal xeric grassland or steppe. CAF operates on a continuous dryland-crop rotation system comprising winter wheat and spring crops. CAF is located in the high rainfall zone of the Pacific Northwest region and the soil type is classified as Mollisol (Naff, Thatuna and Palouse Series) [71]. Sampling occurred from September 8-12, 2022, post-harvest of spring crops, to reduce plant’s impact on soil microbial communities. This period was during the dry season preceding the concentrated autumn rainfall.

Topsoils were collected from the eastern region of the CAF at a depth of 10–20 cm, other than Soil 1 & 2 (depth of 0–10 cm). Eastern CAF practices no-tillage which eliminates soil inversion and mixing of the soil surface to 20 cm. The N fertilizer in this field has been primarily deep banded to depths of approximately 7 to 10 cm during the time of application, which creates a spike of nitrate resource in the soil depth we sampled. Each soil sample was obtained by cutting down through the hardened dry soil with a spade in a circular motion to create a cylindrical cake of soil of radius 10– 20 cm until the desired depth. Each soil sample was not merged from sampling multiple replicates due to differences in pH in different locations. Samples were collected within a diameter of 500 m within the CAF to minimize the variation of edaphic factors other than pH. The large variation in soil pH comes from the long-term use of ammoniacal fertilizers and associated N transformations, which may undergo nitrification resulting in the release of H ions. In combination, spatial pH variation increases with field-scale hydrologic processes that occur under continuous no-tillage superimposed over a landscape that has experienced long-term soil erosion.

To maximize the coverage of sampled native pH, we used a portable pH meter (HI99121, Hanna Instruments, Smithfield, RI, USA) to directly measure and estimate the soil pH without having to make slurry on site to determine whether to sample the soil before sampling. For accurate pH values, pH was measured in the laboratory using a glass electrode in a 1:5 (soil to water w/w) suspension of soil in water (protocol of International organization for standardization, ISO 10390:2021), where 7g of soil was vortexed with 35ml of Milli-Q filtered water, spun down, and filtered with (0.22 µm) pore size. With these pH values, we selected 20 topsoil samples that are well spread across a range of pH from 4.7–8.32 with intervals of 0.1–0.6. Twenty soil samples were sieved (<2 mm), removed of apparent roots and stones, and gravimetric water content was determined (105 *^◦^*C, 24h). The sieved samples were stored in the fridge for 0-3 months until the incubation experiment. For sequencing the initial community, subsamples were stored in *−*80 *^◦^*C until the DNA extraction. The twenty soils were sent to the Research Analytical Laboratory (University of Minnesota, USA) to measure soil texture (soil particle analysis; sand, silt, clay composition), total carbon and nitrogen, and cation exchange capacity. The soils were also sent to Brookside Laboratories, Inc. (New Bremen, OH, USA) for a standard soil analysis package (Standard Soil with Bray I phosphorus). Twenty soils had relatively similar edaphic properties: 5–9% gravimetric water content (g/drysoilg), soil texture of silty clay or silty clay loam with 0% sand and 32–43% clay, and C:N ratio of 12–16 with 1–1.9% total carbon (wt/wt) (Table S3).

### Soil rewetting, constructing soil pH titration curves, and pH perturbation experiments

To mimic the autumn rainfall in the Pacific Northwest region and minimize the effect of spiking microbial activity by rewetting dry soils [72], we rewetted the sieved soil for 2 weeks before the perturbation experiments at room temperature with sterile Milli-Q water at 40% of each soil’s water holding capacity. After resetting, a soil slurry was made by adding 2mM sodium nitrate solution to the soil (2:1 w/w ratio of water to soil). The slurry was then transferred to 48-deep well plates (2.35ml of slurry per well) for incubation under anaerobic conditions (950 RPM, 30 *^◦^*C) for 4 days. Anaerobic incubation was performed in a vinyl glove box (Coy Laboratory Products 7601-110/220) purged of oxygen with a 99%/1% N2/CO2 gas mixture, where the gaseous oxygen concentration was maintained below 50 ppm to prevent aerobic respiration [73].

To perturb the soil pH to desired levels, we constructed each soil’s pH titration curve for the 20 soils with varying native pH to know exactly how much acid or base to add to each soil sample. To do so, separate from the main pH perturbation experiment, we added 23 different levels of HCl (acid) or NaOH (base) in the slurry, final concentrations ranging from 0–100 mM HCl or NaOH.

We additionally tested whether the anion of acid (Cl*^−^*) or the cation of base (Na^+^) had a distinctive effect on the nitrate reduction dynamics, which was not the case (for results, see Fig. S12 and SM). We colorimetrically measured the pH (see section below) immediately after and 4 days after adding each well’s designated amount of acid/base. Due to natural soil’s buffering capacity, it takes 1-2 days to stabilize its pH level. Thus, we used the endpoint (after 4 days) pH measurements for all pH perturbations. We did a spline interpolation on the titration data points, plotting endpoint pH (y-axis) against acid/base input (x-axis), to compute how much HCl and NaOH needs to be added to the soil to obtain 13 different levels of pH with *≈* 0.4 intervals ranging from pH 3 to 9, including the pH level without the addition of any acid or base. For Soil 19 and Soil 20, we had only 7 and 3 perturbed pH levels respectively, because the strong buffering capacity of these soils (native pH over 8) limited the range of pH perturbation.

For the main pH perturbation experiment, the computed levels of concentrated HCl or NaOH were added to the slurry in the 48-deep well plate with and without chloramphenicol treatment for each perturbed pH level in triplicates. The plates were immediately transferred to the shaking incubator (950 RPM in Fisherbrand Incubating Microplate Shakers 02-217-759, 3 mm orbital radius, 30 *^◦^*C) inside the anaerobic glove box and incubated for 4 days. For chloramphenicol-treated (CHL+) samples, we added concentrated chloramphenicol solution to the slurry to obtain a final concentration of 1 g/L. To gauge the effect of the 2mM nitrate, we had no-nitrate controls (0 mM nitrate) for both CHL+/treatments in the unperturbed pH conditions. With antifungal cycloheximide controls (200 ppm) for all 20 soils, we confirmed that fungal activity minimally affects nitrate utilization dynamics (Fig. S4). We also confirmed that abiotic nitrate/nitrite reduction does not occur by measuring metabolic dynamics of autoclaved soil (120 *^◦^*C, 99 minutes, autoclaved 5 times every 2 days) (Fig. S5). To offset the effect of increasing metabolite concentration due to evaporation throughout the 4-day incubation period, we used the wells with just 2mM nitrate, nitrite, and ammonium solutions to correct for evaporation in the slurry samples for every time point. The values of the gravimetric water content of each soil were taken into account to correct for the dilution of 2mM nitrate due to moisture in the soil. After the incubation, the plates were stored in *−*80 *^◦^*C for sequencing endpoint communities.

### Time-series slurry sampling, extraction, and colorimetric assays to measure nitrate, nitrite, ammonium, WSOC, and pH

To obtain the metabolic dynamics, we subsampled 60 µL of the slurry into 96-well plates 10 times throughout 4 days (0, 4, 8, 19, 25, 31, 43, 55, 67, 91 hrs), where the initial time point (T_0_) is the time of pH perturbation and the start of anaerobic incubation. To measure nitrate and nitrite dynamics, extracts were prepared from the sampled slurries by adding and vortexing 2 minutes with 90 µL of 3.33 M KCl solution (final concentration of 2 M KCl) and centrifuging at 4000rpm for 5 minutes. The supernatant was filtered at 0.22 µm with a vacuum manifold to remove soil particles that could interfere with colorimetric assays. Concentrations of nitrate and nitrite in the extracts were determined colorimetrically using the Griess assay [74] and vanadium (III) chloride reduction method, following the protocol outlined previously [73]. We confirmed that 95%–99% of the nitrate in the soil can be accurately retrieved and detected using this method, as verified by nitrate spike-in and extraction experiments in the soil. For a subset of 20 soils (Soil 1, 5, 12, and 17), the ammonium dynamics were measured colorimetrically using the Salicylate-hypochlorite assay from the soil extracts [75]. Chloramphenicol treatments in the samples (CHL+) led to consistent detection of 0.5 mM NH^+^ due to its N-H moiety. The salicylate-hypochlorite assay is also affected by the amount of base (NaOH) in the samples, resulting in slightly lower detection of chloramphenicol in the CHL+ samples (0.45mM NH^+^ in 100mM NaOH perturbations). Taking advantage of these control measurements, we used the constant NH_4_^+^ levels in the controls without 2mM NO_3_*^−^* (No Nitrate controls) in the CHL+ samples for each soil to offset the NaOH effect in the CHLsamples and subtracted NH^+^ levels caused by chloramphenicol in CHL+ samples.

For water-soluble organic carbon (WSOC) measurements, we subsampled 60 µL of the slurry into 96-well plates at T_0_ and endpoint (4 days). Then, soil extracts were prepared by adding, vortexing with 90 µL Milli-Q water, centrifuging at 4000rpm for 5 minutes, and 0.22 µm filtering the supernatant. Concentrations of the organic carbon in the supernatant were measured colorimetrically by the Walkley-Black assay, which uses dichromate in concentrated sulfuring acid for oxidative digestion [76]. We subtracted 0.4 Cmg/ml from the CHL+ samples because chloramphenicol gave rise to a measured value of 0.4 WSOCCmg/ml without additional carbon. For pH measurements, we subsampled 100 µL of the slurry into 96-well plates at T_0_ and the endpoint. Then, soil extracts were prepared by adding, vortexing with 150 µL KCl solution (final concentration of 1 M KCl), centrifuging at 4000rpm for 5 minutes, and 0.22 µm filtering the supernatant. pH of the 120 µL supernatant was determined colorimetrically by adding 4ul of the multiple indicator dye mixture via the protocol described previously [77]. The reason we used 1 M KCl method for pH measurement (ISO 10390:2021) was that, contrary to the KCl method, the H_2_O method (using water instead of 1M KCl) resulted in a highly yellow coloration of the supernatants in strong basic perturbed samples, which interfered with the wavelength of the colorimetric pH assay. For samples of pH outside the range of the assay (below pH 3 and above pH 9), we used a pH micro-electrode micro (Orion 8220BNWP, Thermo Scientific, Waltham, MA, USA).

### DNA extraction with internal standards, library preparation, and 16s rRNA amplicon sequencing

We performed 16S amplicon sequencing on half of all samples: 10 (3, 5, 6, 9, 11, 12, 14, 15, 16, 17; Table S3) out of 20 soils were sequenced before perturbation and at the endpoint in both CHL+ and CHL-conditions, totaling 1,085 amplicon sequencing measurements. Genomic DNA was extracted from 500 µL aliquots in a combined chemical and mechanical procedure using the DNeasy 96 PowerSoil Pro Kit (Qiagen, Hilden, Germany). Extraction was performed following the manufacturer’s protocol, and extracted DNA was stored at *−*20 *^◦^*C. To estimate the absolute abundance of bacterial 16S rRNA amplicons, we added known quantities of genomic DNA (gDNA) extracted from *Escherichia coli* K-12 and *Parabacteroides* sp. TM425 (samples sourced from the Duchossois Family Institute Commensal Isolate Library, Chicago, IL, USA) to the slurry subsamples before DNA extraction. Equal concentrations of gDNA from these two strains were added. Both strains have identical rRNA copy numbers of 7 and comparable genome sizes of approximately 5000 kb. DNA Library preparation was performed using the 16S Metagenomic Sequencing Library Preparation protocol with a 2-stage PCR workflow (Illumina, San Diego, CA, United States). The V3–V4 region was amplified using forward primer 341-b-S-17 (CCTACGGGNGGCWGCAG) and reverse primer 785-a-A-21 (GACTACHVGGGTATCTAATCC) [78]. We confirmed using gel electrophoresis that the negative samples containing all reagents did not show visible bands after PCR amplification. Sequences were obtained on the Illumina MiSeq platform in a 2 *×* 300 bp paired-end run using the MiSeq Reagent Kit v3 (Illumina, San Diego, CA, United States) with 25% PhiX spike-ins. A standardized 10-strain gDNA mixture (MSA-1000, ATCC, Manassas, VA, USA) was sequenced as well to serve as a positive control, which was confirmed to have relatively uniform read counts after assigning taxa.

### Model and fitting

#### Consumer-resource model

Consider a consumer-resource model with one consumer variable (functional biomass x(t), OD/biomass) and two resource variables (nitrate A(t) and carbon-nutrient C(t), mM), which evolves in time (t, day). The ordinary differential equations (ODEs) of the consumer-resource model can be expressed as:

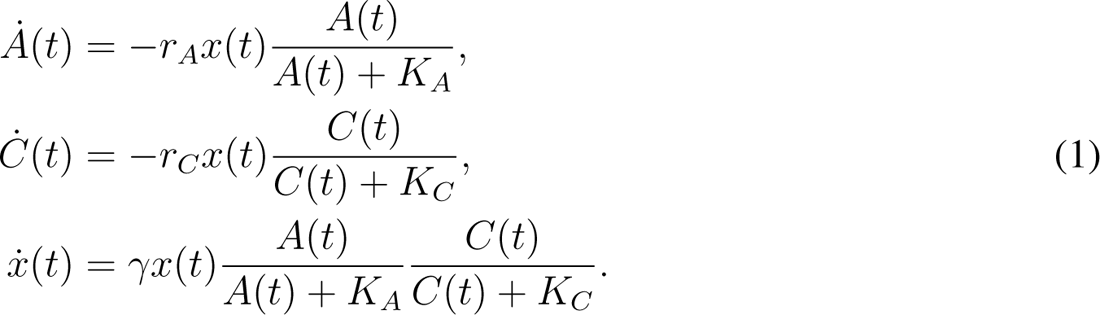

The first two equations of (1) represent the resource consumption rates, which are determined by the functional biomass (x, biomass), the maximum consuming rates per unit biomass (r*_A_* and r*_C_*, mM/biomass/day), and the Monod functions (A/(A + K*_A_*) and C/(C + K*_C_*), dimensionless). Here assume the affinities (K*_A_* and K*_C_*, mM) to be fixed and small. So the Monod functions can be deduced to 1 when A *≫* K*_A_* or C *≫* K*_C_*, and can be deduced to 0 when A *→* 0 or C *→* 0. The third equation represents the growth of functional biomass, which is determined by the maximum growth rate per biomass (γ, 1/day) and the multiplication of two Monod terms indicating the fact that nitrate and carbon are non-substitutable (electron acceptor and donor respectively). The growth is exponential (x(t) = x(0)e*^γt^*) when both A *≫* K*_A_* or C *≫* K*_C_*, but growth stops when either A *→* 0 or C *→* 0. Therefore, in this model, the growth of biomass is limited by both resources, but the consumption of one resource can continue when the other resource runs out and the biomass growth stops. For example, we believe this happens when C *→* 0 in Regime II and the consumption of A continues (Fig. 2).

#### Solution for nitrate dynamics

To find the solution for nitrate dynamics, we rescale the equations by combining parameters: x^ = r*_A_*x, C^^^ = Cr*_A_*/r*_C_*, K^^^*_C_* = K*_C_*r*_A_*/r*_C_*. Therefore, the equations become:

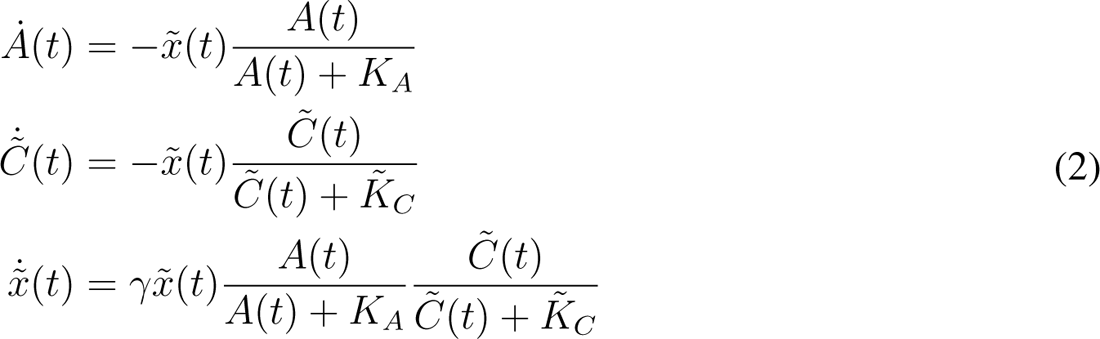

In the rescaled equations (2), the parameters and variables all have units of rates (nitrate per time): [x^] = mM/day and [C^^^] = [K^^^*_C_*] = mM. Therefore, the solution of nitrate dynamics only depends on three parameters (γ, K*_A_*, K^^^*_C_*) and three initial conditions (A_0_, C^^^(0), x^(0)). Since the affinities are very small (K*_A_ ≈* 0.01mM, K^^^*_C_≈* 0.01mM), the solution of biomass approximately equals to x^ = x^(0)e*^γt^* before the time at which growth stops t*^∗^*. So the resource dynamics before t*^∗^* are approximately A = A_0_ *−* x^(0)(e*^γt^ −* 1)/γ and C^^^ = C^^^(0) *−* x^(0)(e*^γt^ −* 1)/γ. Accordingly, the time at which growth stops is given by t*^∗^* = log min(A_0_, C^^^(0))γ/x^(0) + 1 /γ. If C^^^(0) < A_0_, the nitrate dynamics after t*^∗^* and before running out are given by A = A(t*^∗^*) *−* (γC^^^(0) + x^(0))(t *−* t*^∗^*). As a result, the nitrate consumption rate after t*^∗^* is γC^^^(0) larger than the initial rate x^(0).

#### Least-square fitting scheme

To infer the model parameters from the metabolite measurement, we use the least-square fitting scheme to find the closest dynamic curves to the time-series data. Our metabolite measurement including the time points (<u>t</u>*^−^* = [t_2_*^−^*, t_3_*^−^*, …, t_N_*^−^*]) and nitrate amount (<u>a</u>*^−^* = [a_1_*^−^*, a_2_*^−^*, …, a_N_*^−^*]) for each CHL- sample, and the measurements of <u>t</u>^+^ and <u>a</u>^+^ for a corresponding CHL+ sample. We set up the loss function as the mean-squared-error (MSD):

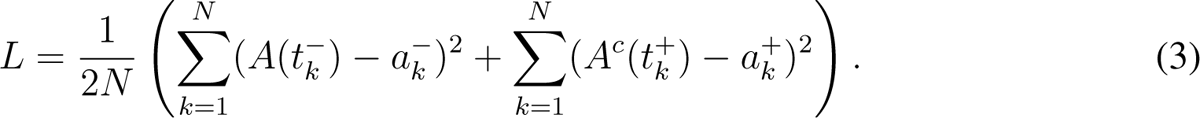

Here the functions A(t) and A*^c^*(t) are theoretical solutions of the consumer-resource model (2) for CHL-/+ conditions, respectively. Because the nitrate dynamics A(t) and A*^c^*(t) are determined by the parameter set Θ = *{*x^(0), C^^^(0), A_0_, A*^c^*, γ, K*_A_*, K^^^*_C_}*, we minimize the loss function L(Θ) to get the optimal model parameters Θ*^∗^*. We note to the readers that three parameters are fixed (γ = 4.8day*^−^*^1^, K*_A_* = K^^^*_C_*= 0.01mM) as justified by the sensitivity analysis in the following paragraph. Note, that these parameters were globally fixed across all the data. We also use different initial nitrate (A_0_ and A*^c^*) in the functions A(t) and A*^c^*(t). The optimization algorithm is the interiorpoint method which is built in the MATLAB <u>fmincon</u> function. The codes and data are available at https://github.com/SiqiLiu-Math/xxx. The fitting errors over all samples are shown in Fig. S7, in which the root-mean-squared-error (RMSE, L(Θ*^∗^*)) and the error per datapoint (*|*A(t*k^−^*) *−* a*k^−^|* or *|*A*^c^*(t*k*^+^) *−* a^+^*|*) are normalized by the input value of nitrate (2mM).

#### Sensitivity analysis on model parameters

γ, K*_A_*, and K^^^*_C_*were globally fixed to one value across all data. Here we justify this decision. We analyzed the sensitivity of γ, K*_A_*, and K^^^*_C_*on simulated dynamic data. To reflect the three typical dynamics (regimes) observed from the measurement, we simulated three nitrate curves by setting up the initial conditions to be x^(0) = 0.01, 0.1, 0.001mM/day and C^^^(0) = 0.005, 0.05, 2mM, respectively. Other parameters are given by A_0_ = A*^c^* = 2mM, K*_A_* = K^^^*_C_*= 0.01mM, γ = 4day*^−^*^1^. We then used different fixed values of parameters to fit the three examples. In the first row of Fig. S8, we used different fixed γ values from γ = 2day*^−^*^1^ to γ = 6day*^−^*^1^ to fit three simulations. We demonstrate very small mismatches (RMSE< 5%) from these variations of parameter values, which are almost invisible in Regime I and Regime II fittings. In the second and the third row of Fig. S8, we use different fixed K*_A_* and K^^^*_C_*values from 10*^−^*^4^mM to 1mM to fit three simulations. When K*_A_* < 0.1mM or K^^^*_C_*< 0.1mM, the mismatches were again very small (RMSE < 1%) and invisible. These results indicate that the fixed values of γ, K*_A_* and K^^^*_C_*are insensitive in large ranges.

#### Determination of regime boundary with model parameters

To define the regime boundaries, we examined the distributions of each parameter’s value. x^(0) had a bimodal distribution (Fig. S9A). This bi-modality becomes more evident when we separately observe its distribution from the left half (perturbed pH < 4) and right half (perturbed pH > 6) of the parameter space displayed in the perturbed pH vs. native pH grid in Figure 3C (Fig. S9B). Therefore, we set the threshold for the x^(0) boundary where these two modes are evidently separated (x^(0) = 0.05). The distribution of γC^^^(0) exhibited a significant mode around 0, prompting us to set the threshold (γC^^^(0) = 1.5) at the tail region, where the γC^^^(0) threshold also separated the Regime III samples in the top-left quadrant of the x^(0) vs. γC^^^(0) scatter plot (Fig. 3A). The separation of Regime I and Regime II data points may not be clear cut in the x^(0) vs. γC^^^(0) scatter plot (Fig. 3A). However, when we plot x^(0) of individual soils from different native pH levels (Fig. S9E), especially in the natively acidic soils, the transition from Regime II (large x^(0)) to Regime I (small x^(0)) is evident going towards more acidic pH perturbations because of the large x^(0) levels sustained over a wide pH range in Regime II.

### Sequence data analysis

#### Sequencing data preprocessing and assigning taxonomy to ASVs with DADA2

Raw Illumina sequencing reads were stripped of primers, truncated of Phred quality score below 2, trimmed to length 263 for forward reads and 189 for reverse reads (ensuring a 25-nucleotide overlap for most reads), and filtered to a maximum expected error of 4 based on Phred scores; this preprocessing was performed with USEARCH ver. 11.0 [79]. The filtered reads were then processed with DADA2 ver. 1.18 following the developers’ recommended pipeline [80]. Briefly, forward and reverse reads were denoised separately, then merged and filtered for chimeras. For greater sensitivity, ASV inference was performed using the DADA2 pseudo-pooling mode, pooling samples by soil. After processing, the sequencing depth of denoised samples was 10^4^–10^6^ reads per sample. Low-abundance ASVs were dropped (≲ 10 total reads across all 1085 samples), retaining 34 696 ASVs for further analysis. Taxonomy was assigned by DADA2 using the SILVA database ver. 138.1, typically at the genus level, but with species-level attribution recorded in cases of a 100% sequence match.

#### Computing absolute abundance with internal standards of each ASV per sample

As an internal control, we verified that the ASVs corresponding to the two internal standard genera *Escherichia-Shigella* and *Parabacteroides* were highly correlated with each other as expected (person correlation ρ = 0.94). These ASVs were removed from the table and combined into a single reference vector of “spike-in counts”. The spike-in counts constituted 8.9 *±* 8.8% of the total reads in each sample. For downstream analysis, the raw ASV counts in a sample were divided by the spike-in counts of the internal standard per sample to obtain the absolute abundance of the ASV in a sample. Total biomass per sample was obtained by dividing the total raw read counts with the spike-in counts of the sample.

#### Differential abundance analysis to identify enriched ASVs

We conducted differential abundance analysis to statistically determine which amplicon sequence variants (ASVs) were significantly enriched in the Nutrient-limiting regime (Regime II) or the Resurgent growth regime (Regime III), respectively. To do so, we identified enriched ASVs for each perturbed pH condition in each native soil comparing endpoint chloramphenicol-untreated (CHL-) samples with endpoint chloramphenicol-treated (CHL+) samples. For each native soil, we then compiled a list of enriched ASVs by aggregating a union set of enriched ASVs across perturbed conditions that belong to Regime II (or Regime III). To remove ASVs that could be false-positive nitrate reducers, we similarly performed differential abundance analysis to identify ASVs that are enriched in no-nitrate controls (nitrate*^−^*) by comparing endpoint chloramphenicoluntreated (CHL& nitrate*^−^*) samples with endpoint chloramphenicol-treated (CHL+ & nitrate*^−^*) samples. This filtering was done when inferring nitrate reducer biomass (Fig. S11C&D) and inferring the Regime III strains (Fig. S17). For each native soil, we only had nitrate*^−^* controls for the condition without acidic/basic perturbation. We assumed that these enriched ASVs in no-nitrate conditions (NNresponders) without acid/base perturbation would also be false-positive nitrate reducers in other acidic or basic perturbation levels. For each native soil, we filtered out these falsepositive NNresponders from the aggregated list of Regime II (or Regime III) enriched ASVs.

To identify the ASVs enriched for each perturbed pH level, it was necessary to determine what change in recorded abundance constitutes a significant change, relative to what might be expected for purely stochastic reasons. The relevant null model would combine sampling and sequencing noise with the stochasticity of ecological dynamics over a 4-day incubation, and cannot be derived from first principles. However, since all measurements were performed in triplicate with independent incubations, the relevant null model can be determined empirically. The deviations of replicate-replicate comparisons from 1:1 line were well-described by an effective model combining two independent contributions, a Gaussian noise of fractional magnitude c_frac_ and a constant Gaussian noise of magnitude c_0_ reads, such that repeated measurements (over biological replicates) of an ASV with mean abundance n counts are approximately Gaussian-distributed with a standard deviation of σ(c_0_, c_frac_) = (c_frac_n)^2^ + c^2^ counts. In this expression, c_frac_ was estimated from moderateabundance ASVs (> 50 counts) for which the other noise term is negligible; and c_0_ was then determined as the value for which 67% of replicate-replicate comparisons are within *±*σ(c_0_, c_frac_) of each other, as expected for 1-sigma deviations. This noise model was inferred separately for each soil and each perturbed pH level, as the corresponding samples were processed independently in different sequencing runs. For example, the parameters in Soil 11 were c_frac_ = 0.21 *±* 0.04 and c_0_ = 4.5 *±* 0.7 counts (Fig. S25).

The model was used to compute the z-scores for the enrichments of absolute ASV abundances in CHLtreatments against CHL+ controls (three independent z-scores from three replicate pairs; rep1-rep1, rep2-rep2, rep3-rep3). The median z-score was assigned to each ASV for each perturbed condition. In consideration of ASVs with 0 read count in either CHL-/+ samples, all raw ASV counts were augmented by a pseudocount of 0.5 and divided by the per-sample spike-in counts, yielding values that can be interpreted as the absolute biomass of each taxon (up to a factor corresponding to the copy number of the 16S operon), measured in units where 1 means as many 16S fragments as the number of DNA molecules in the spike-in. Significantly enriched ASVs were identified in each perturbed condition as those with z-scores greater than z = Φ*^−^*^1^(1 *−* α/2/n*_ASV_*), where Φ*^−^*^1^(x) is the inverse CDF of the standard normal distribution, α = 0.05, and n*_ASV_* as the number of nonzero ASVs in a given sample. This critical z-score (z = 4.2, when n*_ASV_* = 2000 for enriched ASVs and z = 4.3, when n*_ASV_* = 2500 for filtering no-nitrate responders (NNResponders)) corresponds to a two-tailed Bonferroni-corrected hypothesis test at significance level α under the null hypothesis that counts in the CHLand CHL+ conditions are drawn from the same distribution. These analyses were performed using custom MATLAB (Mathworks, Inc) and R scripts, which are available on the GitHub data repository for the present manuscript; for additional technical details, the reader is referred to the detailed comments in these scripts.

#### Non-negative matrix factorization (NMF) analysis on phylum-level growth folds

To analyze the abundance change at the phylum level, we compute the growth fold of each phylum at each condition. For each phylum, we compute the absolute abundance by aggregating the abundances of all ASVs within that phylum. Taking CHL+ abundance Abs^+^ as the reference abundance and CHL- abundance Abs*^−^* as the endpoint abundance (where Abs denotes taxon abundance normalized to internal standard), the logarithm of the growth fold for phylum i and condition j is given by g*_ij_* = log(Abs*_ij_^−^* +10*^−^*^3^)*−*log(Abs*_ij_*^+^ +10*^−^*^3^). Note that we use CHL+ abundance as reference instead of the initial abundance (at T0), to account for any effects on read counts unrelated to growth which would be common between CHL+ and CHL- conditions (e.g. direct effect of acid/base addition), allowing us to focus only on growth-mediated abundance changes. We also set all negative g*_ij_* to 0 since we are focusing on growth. For all 130 conditions (10 soils *×* 13 perturbations) and 40 phyla, the phylum-level growth folds G is a 40 *×* 130 matrix. For each phylum, the row vector ⃗g*_i_* represents how it grows under different conditions (see Fig. S15 for the growth vectors of the first 10 phyla). In order to reduce the dimensionality of the growth matrix and extract the main features of the growth vectors, we use non-negative matrix factorization (NMF) to decompose the growth matrix G = W *∗* H by factor 2. Here the feature matrix H is of size 2 *×* 130, and the two rows ^⃗^h_1_ and ^⃗^h_2_ are two modes of growth vectors (shown in Fig. S15B). Therefore, the growth vector of phylum i is thus decomposed as ⃗g*_i_ ≈* w*_i_*^1⃗^h_1_ + w*_i_*^2⃗^h_2_, while the weights w*_i_*_1_ and w*_i_*_2_ are from the 40 *×* 2 weight matrix W. The weights of all 40 phyla are plotted in Fig. S15B, showing that Firmicutes are mostly composed by the second mode ^⃗^h_2_ and other phyla are mostly composed by the first mode ^⃗^h_1_. Additionally, Bacteroidota and Proteobacteria show the most significance of the first mode. This decomposition keeps 93.36% of the original growth matrix.

#### Genotyping enriched ASVs with PICRUSt2

To understand what traits make Resurgent growth strains unique, we used PICRUSt2 ver. 2.5.2 [40] to infer putative genotypes of the enriched ASVs in the Resurgent growth regime (Regime III) (Fig. S18C). Using the script “place seqs.py” in the pipeline, we matched the representative 16S rRNA sequences of each amplicon sequence variant (ASV) to PICRUSt2’s curated reference genome database (multiple sequence alignment). Then, using the “hsp.py” script and default parameters, we predicted KEGG orthologs (KO) abundance of each ASV with the matched reference genome (hidden-state prediction). To narrow down to KOs/genes related to denitrification and Dissimilatory Nitrate Reduction to Ammonium (DNRA), we focused on nitrate reductase in denitrification (*narG*/K00370, *narH*/K00371, *narI*/K00374, *napA*/K02567, *napB*/K02567) and nitrite reductase to ammonium (*nirB*/K00362, *nirD*/K00363, *nrfA*/K03385, *nrfH*/K15876). To track which KOs were enriched at which pH in the 89 families used in the peak pH analysis (see SM for peak pH analysis), we summed the relative abundance (reads / total reads of each perturbed pH level in CHL-) of the ASVs belonging to each family that possessed at least 1 predicted gene respectively for *narGHI*, *napAB*, *nirBD*, and *nrfAH*. Then, we plotted their relative abundance values across pH for all soils, indicated by the intensity of the point’s colors (Fig. S18).

#### Taxonomy of growing strains in Regime III varies with soil native pH

To further investigate whether the taxonomic identity of Resurgent growth (Regime III) strains varies across natural pH environments, we performed a regularized regression analysis to see if we can predict the native pH level of the source soil from the presence or absence of taxa that grow in Regime III at the ASV, Species, Genus, Family, or higher taxonomic levels. The Resurgent growth strains were determined by the differential abundance analysis as described previously. Should our findings confirm that the prediction of native soil pH is feasible based on the taxonomic variation of these strains, it means that the strains responsible for growth in Regime III *depend on the long-term pH of the soil*. To do so, we used the sequencing data to build a matrix where the rows are samples (including three biological replicates) belonging to the Resurgent growth regime (Regime III), where each row has a corresponding native pH value of the original soil. There are 10 source soils with different native pH levels, and each soil has 3 to 6 pH perturbed samples (replicates) of which metabolite dynamics are classified as the Resurgent growth regime. The matrix’s columns are different taxa belonging to the identified Resurgent growth strains, either in ASV, species, genus, family, or higher levels. Each element of the matrix is 0 if absent and 1 if present in the sequencing data of the sample. Because the presence and absence of taxa can randomly depend on the random sampling depth of each sample, we test varying threshold values (0, 0.001, 0.005 relative abundance) to call the taxa present if their relative abundance is greater than the threshold (effects shown in Fig. S22F).

The regularized regression was performed to predict the native pH of the source soil of the samples from the presence and absence of taxa using only additive terms and LASSO regularization to avoid overfitting [81]. To estimate the regularization hyperparameter, tenfold cross-validation was performed on the samples from ten different soils with different native pH levels. All models were fit using the package glmnet in R version 4.1.4. To make predictions of the native pH, we used two strategies. First, ‘in-sample’ predictions used all available data points to fit the regression coefficients and predicted native pH using those coefficients. Second, to ask whether we can still predict the native pH without the model seeing the samples belonging to that specific native pH level, we implemented a ‘Leave-one-soil-out’ (LOSO) procedure where all the perturbed samples from one native soil were left out as a test set, and the model was trained on the remaining data to fit the regression coefficients. Then, we used the model to predict the native pH of the left-out samples (out-of-sample prediction). The observed versus predicted pH values are shown in the scatter plots (Fig. S22A). The prediction quality (R^2^, coefficient of determination = 1 *−* SS*_E_*/SS*_T_*, sum of squares error over total sum of squares) was computed using the mean predicted and mean observed native pH levels for each soil (for different taxonomic levels and prediction strategies, see Fig. S22 E & F, negative R^2^ values indicate the predictions are worse than just predicting the pH as the mean predicted pH). To ascertain that our high prediction quality was not a random artifact, we randomly permuted the native pH values of our soils 1000 times and then predicted in-sample the native pH to obtain 1000 R^2^ values. We computed the threshold R^2^ value that corresponds to the p-value of 0.05 (top 50th R^2^ value out of 1000 instantiations) and compared it with the R^2^ value that we have obtained with our true native pH predictions (Fig. S22G).

### Testing the effect of different bases and salts on nutrient release

To see the effects of different bases (NaOH and KOH) on nitrate reduction dynamics, we added different concentrations of NaOH and KOH (final concentration of 0, 8, 16, 24 mM in the slurry), following the same protocol previously described (without chloramphenicol), to measure the nitrate and nitrite dynamics (Fig. S12) using Soil 6 (Table S3). In addition, to test the effects of Na^+^, K^+^, Cl*^−^* separately, we added different concentrations of salts (NaCl, KCl) (without chloramphenicol and without adding any acid/base) and measured the metabolite dynamics (Fig. S12).

### Nutrient amendment experiments with slurries

To experimentally determine what nutrient was limiting growth in the Nutrient-limiting regime, we conducted nutrient amendment experiments respectively with glucose, succinate, sodium acetate, ammonium chloride (NH_4_Cl), monosodium phosphate (NaH_2_PO_4_), and sodium sulfate (Na_2_SO_4_) (for results, see Fig. 4D and Fig. S14). Among them, succinate (pK*_a_* = 4.21 and 5.64, 25 *^◦^*C), acetate (pK*_a_* = 4.76, 25 *^◦^*C), and phosphate (pK*_a_* = 2.2, 7.2, and 12.4, 25 *^◦^*C) were strong candidates for the limiting nutrient according to our soil nutrient release hypothesis, due to their anionic nature in mid-range pH (5-7). The incubation was conducted following the same protocol using Soil 6 (Table S3) without chloramphenicol and without adding any acid/base. Concentrations were either in C mM, N mM, S mM, or P mM with final concentrations in slurry varying from 0 to 5 mM, each in triplicates. Because we have previously tested the effect of Na^+^ and Cl*^−^* to be negligible in nitrate dynamics, the effect of these amendments can be attributed solely to C/N/S/P nutrients other than Na^+^ and Cl*^−^*.

## Acknowledgements

We acknowledge Ian Leslie for coordinating and guiding the soil sampling in the Cook Agronomy Farm (CAF, Pullman, WA, USA). We thank Maren L. Friesen and Janice Parks for providing temporary lab space and facility to store sampled soils and conduct pH measurements at Washington State University, USA. We thank Timothy Paulitz, Ian Leslie, and David Huggins for critical advice regarding soils. This research was a contribution from the Long-Term Agroecosystem Research (LTAR) network. LTAR is supported by the United States Department of Agriculture. We acknowledge the Duchossois Family Institute at the University of Chicago (Chicago, IL, USA) for providing *Parabacteroides* sp. TM425 strain to use as internal standard during sequencing. We are grateful to Fatih M. Abasiyanik and Ha-Na Shim in the Single Cell Immunophenotyping Core and Craig DeValk in the Ranganathan lab, both at the University of Chicago, for their guidance on operating the sequencer. We also thank members of the Kuehn lab and Mani lab for helpful discussions.

## Funding

This work was supported by the National Science Foundation Division of Emerging Frontiers EF 2025293 (S.K.) and EF 2025521 (M.M.) and by National Science Foundation PHY 2310746 (M.T.). S.K. acknowledges the Center for the Physics of Evolving Systems at the University of Chicago, National Institute of General Medical Sciences R01GM151538, and support from the National Science Foundation through the Center for Living Systems (grant no. 2317138). S.K. and M.M. acknowledge financial support from the National Institute for Mathematics and Theory in Biology (Simons Foundation award MP-TMPS-00005320 and National Science Foundation award DMS-2235451). MM was supported by The National Science Foundation-Simons Center for Quantitative Biology at Northwestern University and the Simons Foundation grant 597491. MM is a Simons Investigator. This project has been made possible in part by grant number DAF2023-329587 from the Chan Zuckerberg Initiative DAF, an advised fund of the Silicon Valley Community Foundation. Any opinions, findings, conclusions, or recommendations expressed in this material are those of the authors and do not necessarily reflect the views of the National Science Foundation.

## Author contributions

K.K.L., M.M., and S.K. conceptualized the research. K.K.L. and S.K. designed the experiments. K.K.L. conducted field soil sampling in the Cook Agronomy Farm under the supervision of D.H.. K.K.L. performed soil processing, characterization, soil pH perturbation experiments, metabolite assays, gDNA extraction, and sequencing, advised by S.K.. K.K.L., S.L., and M.T. performed statistical analysis of the metabolite dynamics and sequencing data, advised by M.T., M.M., and S.K.. S.L. performed consumer-resource model fits and simulations on the metabolite dynamics, advised by M.M. and S.K.. K.C. performed the monoculture experiments to recapitulate linear nitrate dynamics. K.K.L. performed the nutrient amendment experiments in soils. K.K.L. and S.K. wrote the manuscript with input from S.L., M.M., and M.T.

## Competing interests

The authors declare no competing interests.

## Data availability

Data associated with this manuscript will be made publicly available at NCBI BioProject upon publication.

## Code availability

Code associated with this manuscript will be made publicly available at https://github.com/KiseokKeithLee/Soil_pH_perturbation/ upon publication.

## Supplementary Materials

### A detailed description of the three functional regimes: the Acidic death regime (Regime I), Nutrient-limiting regime (Regime II), and Resurgent growth regime (Regime III)

By quantitatively distinguishing the impact of pH on the consumer side (microbial community, x^(0)) and the resource side (growth-limiting nutrient, C^^^(0)), we can ask the mechanism behind functional adaptation during different regimes. Regime II can be called the “Nutrient-limiting” regime. Within this pH range (Fig. 3C), conditions favor the resident population of nitrate reducers; hence it allows a large indigenous nitrate reducer population to perform nitrate reduction (large x^(0)). This specific range of favorable pH levels is determined by the long-term exposure to the native pH of soils (Fig. 3C). In this regime, the increase of nitrate reduction rate is determined by the biomass growth from the available growth-limiting nutrient. Therefore, in Regime II, the adaptive strategy employed by the nitrate-reducing population is to utilize the pre-existing resident species which are rather robust to pH perturbations, and at the same time incrementally increase the resident’s biomass as the resource availability changes with pH perturbations. Going back to the functional dynamics, that is the reason we see a relatively high slope of CHL+ conditions and a slight increase of denitrification rate in CHL- conditions ((b) and (e) of Fig. 3B), demonstrating that the resident nitrate reducers adapt to the new environment in a “nutrient-limiting” manner.

Regime III can be called the “Resurgent growth” regime. As the perturbed pH is increased from Regime II, there comes a critical pH of around 8 where the adaptive mechanism abruptly transitions. When the pH is perturbed beyond the critical point, the previously large biomass of the nitrate reducer population can no longer adapt and perform nitrate reduction (x^(0) *→* 0). On the resource aspect in Regime III, there is a surplus of limiting nutrients, and thus the system is no longer limited by C but limited by nitrate A. These two effects of short-term pH change (both on the consumer and resource aspect) set the stage for the “resurgence”. A rare population, which we will investigate the composition later, appears to have a small biomass initially showing a flat slope in both CHLand CHL+ conditions but later grows exponentially to exponentially deplete the nitrate in Regime III (panel c, f of Fig. 3F). This shows that in Regime III, the adaptive mechanism of the community is to rely on the rare uprising nitrate reducer biomass to rapidly grow in the absence of nutrient limitation.

Regime I can be called the “Acidic death” regime. This regime is at the other end (acidic) of the Nutrient-limiting regime (Regime II). As the perturbed pH is decreased from Regime II, it transitions into a regime where the system fails to adapt. The boundary pH of Regime I and Regime II is influenced by the native pH of the soil, where relatively acidic soils have a lower boundary of pH 3 or 4 and relatively neutral soils have a higher boundary of pH 5 to 6. Similar to what happens when the community enters Regime III, the unfavorable pH diminishes the indigenous nitrate-reducing activity of the soil, indicated by the flat CHL+ dynamics in panels a, d of Fig. 3B. However, unlike Regime III, the perturbed pH does not make the growth-limiting nutrient superfluous but makes it unavailable, making the divergence of CHLand CHL+ dynamics nonexistent (panel a, d of Fig. 3A). These two effects of pH perturbation make it extremely difficult for the community to adapt to the new environment of Regime III, hence the “Acidic death” regime. Another reason we call it that is to highlight the asymmetric effect of acidic and basic perturbations, which has been seldom acknowledged in the literature.

### Detailed mechanism of nutrient release in soils due to change in pH

We turned to soil literature to develop a comprehensive mechanism for the nutrient release mechanism in soil [38, 39]. Soil comprises minerals, organic matter, water, and air. Minerals and organic matter form aggregated clumps of soil particles, categorized by size into sand, silt, and clay. Clay particles, the smallest among them, consist of layers of phyllosilicates. Each layer includes tetrahedral structures of Si^4+^ covalently bonded to four oxygens and octahedral structures of continuous Mg^2+^ or Al^3+^ covalently bonded to six hydroxides [82]. Due to this chemical structure, clay particles possess numerous electrostatically charged sites, including negatively charged (oxygen atom and hydroxide) and positively charged [39] (shown as and + sites in Fig. 4B). The clay particle’s cation exchange sites are negatively charged and form ionic bonds with cations (positively charged ions), while the anion exchange sites are positively charged and bind to anions (negatively charged ions). Both cations and anions can serve as potential nutrient sources. When they are bound to the clay’s exchange sites (brown section in Fig. 4B), they are protected from microbes. However, when they are released and dissolved in the pore water (light blue or pink section in Fig. 4B), they become available to the microbial community. To understand how NaOH or HCl impacts nutrient availability, it’s essential to track whether cations and anions are bound to clay particles or dissolved in the pore water.

The literature on nutrients and pH in soils proposes the following mechanism for nutrient release (Fig. 4B, see more detailed cartoon Fig. S13B) [38, 39]. When NaOH is added to the soil solution, both Na^+^ and OH*^−^* ions (pH-mediated) act to release the anionic nutrients (case 2 of Fig. 4B). First, OH*^−^* deprotonates ion exchange sites in the clay particles increasing the number of cation exchange sites (charge) and decreasing the number of anion exchange sites (+ charged) reducing the capacity of the clay to hold anions. Secondly, Na^+^ can either bind to the clay particle or remain in solution [38]. The (Na^+^) that remains in solution increases the stability of released anions. Overall, increased OH*^−^* increases the anionic nutrients available to the microbial community (Fig. 4B). The converse happens for HCl perturbations (case 1 of Fig. 4B).

### Effect of base cation on nutrient release in soils

Nutrient release is not solely driven by OH*^−^* ions. The base cation plays an important role, and thus whether the base cation prefers to be in the clay particle or the water solution can influence the amount of nutrients available to microbes. It is known that the bigger the size and greater the charge of the cation, the more selectively the cation binds to clay particles. For example, the cation’s binding specificity to the clay particle instead of staying in the solution is in the order of NH^+^ > K^+^ > Na^+^ > Li^+^, divalent cations having greater binding specificity than monovalent cations (e.g., Ca^+^ > K^+^) [39]. Therefore, when the amount of dissolved organic carbon (DOC) was measured after adding Ca(OH)_2_ vs. KOH in equimolar hydroxide, adding KOH resulted in a much higher concentration of dissolved organic carbon (DOC) [38]. Because K^+^ ions less specifically bind to the clay particle and more likely remain in the solution, it would stabilize the released anions better in the solution, presuming that the DOC is mostly anion due to many negatively charged moieties of O*^−^*s. To check if there was a significant difference between monovalent cations, we compared NaOH and KOH treatments for basic perturbations. We found that there was no significant difference in the nitrate utilization rates in the CHL- condition when we added the same concentrations of NaOH and KOH, indicating that the amount of limiting nutrient released was similar (Fig. S12), although the stabilized endpoint pH was different. As a sanity check, we further tested KCl and NaCl treatments and found that K^+^, Na^+^, or even Cl*^−^* (relevant in the HCl addition) ions themselves without OH*^−^* did not affect nutrient release (Fig. S12), which agrees with previous findings [38].

### Recapitulating linear dynamics with monoculture experiments without carbon

Our model and functional dynamics data suggest that the limited carbon leads to a constant rate of nitrate reduction. However, it is difficult to understand the mechanism behind this phenomenon, because if the organic carbon, an electron donor in the electron transport chain, is coupled to the reduction reaction of nitrate (terminal electron acceptor), the depletion of organic carbon will likely stop the nitrate reduction performed by the nitrate reductase enzymes. This will cause the nitrate reduction rate to be close to 0 rather than the observed constant rate. To resolve this contradiction, one hypothesis can be that the cells internally store carbon nutrients (electron donor) to power the electron transport chain (consuming nitrate) without needing to import external carbon nutrients to generate ATPs for the cell’s maintenance energy. To test this hypothesis, we conducted monoculture experiments with a *E. coli* strain and a known denitrifier *Pseudomonas sp.* strain.

### Culturing protocol

Strains were pre-cultured in two stages under aerobic conditions before being transferred to denitrifying (anaerobic) conditions for phenotyping. First, wells of a sterile 24-well plate (Thermo Scientific Nunc Non-Treated Multidishes) were loaded with 1.7 mL of R2B medium. Wells were inoculated with *E. coli* K12 and *Pseudomonas sp.* PDM04 [73] strains from glycerol stocks stored at *−*80 *^◦^*C. The plates were then sealed with a gas-permeable sterile membrane (Breathe-Easier, USA Scientific, 9126-2100). After sealing, the culture was incubated overnight at 0.5 rcf (400 RPM in Fisherbrand Incubating Microplate Shakers 02-217-759, 3 mm orbital radius) and 30 *^◦^*C in aerobic conditions. These cultures reached saturation during this time. Second, wells of a sterile 24-well plate were loaded with 1.7 mL of defined media (15 mM ammonium, 40 mM phosphate buffer with the final medium pH adjusted to 7.3, and trace metals and vitamins, as described in Ref [73]) with 25 mM succinate. Wells were then inoculated with 17 µL of the saturated R2B *E. coli* K12 and *Pseudomonas sp.* PDM04. After sealing, the cultures were incubated at 0.5 rcf and 30 *^◦^*C in aerobic conditions overnight. These cultures reached saturation during this time. Saturated defined media (DM) cultures were washed and normalized to a desired optical density (measured at 600 nm) via dilution into pH 7.4 phosphate-buffered saline (8 g/L H_2_O, 0.2 g/L KCl, 2.68 g/L Na_2_HPO_4_*·*7H_2_O, 0.24 g/L KH_2_PO_4_).

Wells of a sterile 96-deep well plate (Axygen PDW20C) were loaded with carbon-free 1.2 mL DM supplemented with 2mM sodium nitrate which had been allowed to equilibrate in the anaerobic glovebox. These wells were inoculated in the glovebox with 12 µL of OD-normalized aerobic precultures, resulting in starting ODs of 0.1 and 0.01. Additional wells were left blank as no-growth controls. Plates were sealed with a gas-permeable sterile membrane. Cultures were incubated at 30 *^◦^*C and shaken at 950 RPM (Fisherbrand Incubating Microplate Shakers 02-217-759 or Talboys Professional 1000MP, 3 mm orbital radius) for 72 h. Optical densities of initial and endpoint anaerobic pre-cultures were measured using 300 µL of cultures in 96-well optical plates. Nitrate and nitrite concentrations were assayed over time via manual sampling and subsequent Griess assay and vanadium (III) chloride reduction via the protocol described in Ref. [73].

### Linear metabolite dynamics were recapitulated with monoculture experiments

Both *E. coli* K12 and *Pseudomonas sp.* PDM04 strains were able to reduce nitrate even without carbon in the culture media (Fig. S6A). The reduction rate was negligible for *E. coli* strain at a starting OD600 of 0.01 (optical density at 600nm). However, for the denitrifier *Pseudomonas sp.* PDM04 at a starting OD600 of 0.01, not only the rate of nitrate reduction was comparable to what we observed in soils at the Nutrient-limiting regime (Regime II), but the reduction dynamics were strikingly linear. This result directly demonstrates that nitrate reduction can proceed even when carbon is not exogenously available. Our observations are consistent with this hypothesis that the cells can internally store carbon and oxidize this carbon to provide electrons (NADH) to reduce nitrate to nitrite. If the nitrate reduction rate had increased, it would have meant that the functional biomass, or the quantity of nitrate reductase enzyme, increased. The nitrate reduction rates did not increase throughout the experiment (top panel in Fig. S6A). This supports the idea that cells are using nitrate to maintain biomass. Consistently, final OD600 measurements did not detect any significant increase from the initial OD600 as expected. We can now more confidently presume that soil microbial communities are also in the same maintenance state during the linear nitrate dynamics. In sum, these results suggest that the functional biomass in soils can utilize nitrate at a constant rate, even after external carbon is no longer available. Note our model assumes this to be the case (Fig. 2).

In the monoculture experiment, we observed biphasic behavior in the high initial OD600 condition. This phenomenon is challenging to interpret. In the starting OD600 of 0.1, the initial slopes of nitrate reduction dynamics are constant, then after some time, the rates decrease and remain constant (we will call this “late slope”) until the end of the experiment (bottom panel of Fig. S6A). The linear dynamics observed in the “late slope” again still recapitulate the linear dynamics we observed in the soils in the Nutrient-limiting regime, where the microbes could be using nitrate and internally stored carbon to generate maintenance energy. In the soil experiments, the “late slopes” were determined by the increased functional biomass in the model. However, in these monoculture experiments, it was difficult to understand what determines the late slope values. Their biomass had not changed from the beginning according to the endpoint OD600 measurements, hence requiring further investigation of the bacterial physiology. The initial slopes can be roughly explained by their starting biomass (Fig. S6B), where the initial slope for the OD600 0.1 condition was roughly 10 times greater than that for the OD600 0.01 condition, with the fitted initial slopes showing an increase factor ranging between 5 to 19 times.

### Confirming the linear dependence between functional biomass and acid/base input

Although the linear relationship between acid/base input with the total biomass increase during the Nutrient-limiting regime (Regime II) corroborates our proposed nutrient release mechanism, to be more precise, we need to further show that the fold increase of the “functional” biomass is equal to the fold increase of nitrate reduction rate from the nitrate dynamics data. However, when we observe ASVs increasing in absolute abundance from the start to the end of the experiment we cannot assume all ASVs are performing nitrate reduction (for example, some may be growing via fermentation). To address this we filtered out the ASVs that are likely not nitrate reducers by removing ASVs that were enriched in no-nitrate conditions (dark grey NNresponders bar in Fig. S11D, no pH perturbation). To detect the fractional biomass that performs nitrate reduction, we used a differential abundance analysis to statistically determine which amplicon sequence variants (ASVs) were significantly enriched in each pH perturbed condition compared to the CHL+ counterpart serving as a baseline of no growth (see Materials and Methods for details). Then, we summed up the absolute abundance of these ASVs that we inferred as true nitrate-reducing biomass to obtain the functional biomass for each condition (red bar in Fig. S11D). By comparing the fold increase of these functional biomass values (endpoint/initial functional biomass), we showed that indeed the functional biomass increase and nitrate reduction rate increases are similar in different soils (Fig. S11C). While some soils showed very close agreement between the inferred increase in functional biomass and increases in nitrate reduction rate (Soil 11, inset of Fig. S11C), for many soils the relationship was not quantitative. This discrepancy likely arises from the fact that we inferred the taxa that are not nitrate reducers from slurries where the pH was not perturbed. Thus the no-nitrate responders may be distinct as pH is perturbed and this may increase errors in our inference of changes in functional biomass.

### Investigating the taxonomy, pH niche, and the phylogeny of the Resurgent growth strains

#### Determination of peak pH for each family

To elucidate the pH niche of each family in Fig. S18A, we analyzed the relative abundance of the chloramphenicol-untreated (CHL-) conditions of ASVs identified as being enriched in different pH levels (see Differential abundance analysis). Due to the challenge of visualizing a large number of ASVs, we aggregated the relative abundance of ASVs in the same family for each sample, visualizing the data at the family level. To get the representative relative abundance of the family in each perturbed pH level, we took the median of the relative abundance from three biological replicates. To incorporate abundance values at each perturbed pH from all soils with different native pH levels, we placed the perturbed pH values from all native soils from smallest to largest and then binned neighboring 2–4 perturbed pH values (depending on the total number of relative abundance values greater than 0) to compute the median relative abundance of each family within each bin corresponding to its mean pH value. Now, for each family, we have a median relative abundance value assigned to the mean pH of each bin. For each family, we ranked those median relative abundance values across perturbed pH and found the peak pH value with the highest relative abundance, as well as the second peak pH and its corresponding relative abundance. To reduce the number of families to plot, we chose 89 families that have relative abundance values at the second peak pH greater than 0.002 (= 0.2%). After aligning the 89 families with their peak pH from smallest (top of the plot, dark blue color) to largest (bottom of the plot, yellow color), we plotted a ridge plot with the x-axis being perturbed pH level and height corresponding to the median relative abundance of each bin (Fig. S18A). The maximum heights of the ridge for each family were set to the same level by normalizing the maximum relative abundance for each family. Indeed, the family with peak pH over 8 were mostly Firmicutes phylum (Bacillaceae, Clostridiaceae, Paenibacillaceae, Caloramatoraceae, Peptostreptococcaceae, Lachnospiraceae), other than Yersiniaceae family which was Proteobacteria.

#### Constructing a phylogenetic tree with 16S rRNA sequences

To see whether there was phylogenetic convergence among strains with similar pH niches, we used the 16S rRNA sequences of the ASVs to construct a phylogenetic tree (Fig. S18B). To be consistent with the previous peak pH analysis, we used the ASVs that belonged to the 89 families in the previous analysis. We selected one ASV with the largest relative abundance from each family to represent the family and used its 16S rRNA sequence to construct the phylogenetic tree. The phylogenetic tree was constructed by approximating the Maximum likelihood tree with the General time reversible model in FastTree ver. 2.1.9 [83]. The tree was plotted using the plot.phylo function in ape package in R, each node (labeled with the classified genera or species name) colored by its peak pH.

#### Lower-level taxonomy and traits of the Regime III strains

To identify the specific taxa accountable for the emergence of Regime III at a finer taxonomic level, we conducted a differential abundance analysis that statistically determined which Amplicon sequence variants (ASVs) were significantly more abundant in Regime III CHL-samples, compared to CHL+ samples under same perturbed conditions (see Methods). Then, we aggregated the relative abundance of these differential ASVs (i.e., Regime III strains) to assess their contribution to the emergence of Regime III. Notably, their abundance began to rise between pH 7-8 (Fig. S17C), which aligns with or slightly precedes the transition between Regime II and III (Fig. S17D). This increase in relative abundance corresponded with the shift of the nutrient growth parameter γC^^^(0) from zero (Fig. S17E). The analysis revealed that 10 families belonging to Firmicutes (Bacillaceae, Paenibacillaceae, Clostridiaceae, Caloramatoraceae, Peptostreptococcaceae, etc.) and 2 families belonging to Proteobacteria phylum (Legionellaceae and Yersiniaceae) were significantly enriched in Regime III (Fig. S17B). At the genus level, Bacillus, Clostridium, Paenibacillus, and others were identified as the primary contributors to Regime III (Fig. S17A).

We lastly sought to find distinct features of the Regime III strains that differentiated them from other strains to understand why these strains better adapt to perform nitrate reduction in high pH and high carbon conditions. To do so, we classified families by their peak pH obtained by finding the pH level at which its median relative abundance across soils was the highest across different perturbed pH levels (Fig. S17A). We indeed found that the Regime III families had distinct pH niches compared to other strains, having high relative abundance in basic pH (over 8) and in some cases acidic pH (less than 4) but remained rare (< 0.1%) in the mid-range of pH 4-8. One can speculate that their ability to survive and persist in extreme pH perturbations (see Fig. S20A) may be because many taxa in the phylum Firmicutes are spore-forming bacteria species [84]. These strains did not cluster phylogenetically and were dispersed throughout the phylogenetic tree (Fig. S17B, see Methods).

#### pH titration curves and soil’s native pH are shaped by soil’s physicochemical properties

We’ve constructed pH titration curves for the 20 soils from different native pH levels (see Methods, Fig. 6B). Because we titrated both in acid and basic directions with H^+^ and OH*^−^* respectively, we unified the x-axis to OH*^−^* (m mol) by shifting the curves to the right by 0.2 m mol, ensuring each curve starts at 0 m mol OH*^−^*. We then fitted the pH titration curves with a logistic function with 4 parameters (a, x*_mid_*, b, c) as below (visualized in Fig. S23B):

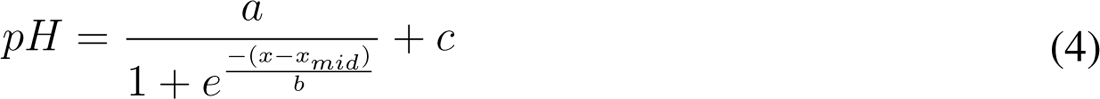

Parameter x*_mid_* strongly correlated with soil’s native pH level (R^2^ = 0.8, Fig. S23D), while parameter c (y-intercept) and a+c (asymptotic y value), scaling parameter (b) stayed mostly constant across soils with different pH levels. This indicates that the titration curve’s general shape is similar for all soils, but the titration curve shifts horizontally depending on the soil’s original pH level.

Soil’s native pH, which determines the horizontal shift of the titration curves, was strongly correlated to the cation exchange capacity (CEC, milliequivalent charge / 100g) (R^2^ = 0.88, Fig. S23D). This was expected because soils with higher CEC will have a greater number of negative charges in the clay particles and hence more likely to adhere to protons. This will result in fewer free protons in the soil pore water and, thus result in more basic pH levels. In the literature, CEC is reported to be determined by soil’s clay particles and its organic matter, because CEC is proportional to how much negative charge the soil has on the aggregate’s surface. However, in our dataset, the percent clay and organic matter did not correlate strongly with CEC (Fig. S23D, see Fig. S23E for percent clay). CEC appeared to be determined by Ca^2+^ ion concentration in the soil and not by other cations (Mg^2+^, K^+^, Na^+^). Soil pH was inversely correlated with S, P, Al, and Fe concentrations, which can either be the cause or result of the soil’s pH (Fig. S23D). In sum, we can attribute the horizontal shift of the pH titration curves to their varying native soil pH levels, which is potentially determined by the CEC and the Ca^2+^ ion concentrations (see summarized diagram in Fig. S23C).

### Evidence for long-term pH adaptation from phyla’s differential response to pH perturbations

We observed that long-term pH variation (different native soil pH) shifts the pH boundaries between functional regimes (Fig. 6). To see if those shifts of pH boundary can be explained by taxa’s differential response to perturbed pH due to long-term pH adaptation, we further asked whether the pH values where the abundances of taxa (Firmicutes, Bacteriodota, and Proteobacteria) exhibit large changes also agree with the boundaries between regimes determined solely by nitrate utilization dynamics (Fig. 3). We observed the growth folds of taxa for the transition from Regime II to Regime III and the survival folds of taxa for the transition from Regime II to Regime I (Fig. S21A). Growth folds were computed by endpoint absolute abundance ratio of Abs*_CHL−_*/Abs*_CHL_*_+_ (chloramphenicol untreated/treated conditions) and survival folds were computed by absolute abundance ratio of Abs*_CHL_*_+_/Abs*_T_*_0_, representing taxa’s endpoint absolute abundance in CHL+ conditions compared to the initial time point (T_0_) for each perturbed pH level.

To understand the transition to the Acidic death regime (Regime I), we observed the survival folds of Proteobacteria and Bacteroidota phyla across perturbed pH levels. Then, we set an identical survival fold threshold for all soils (red lines in Fig. S21) to compute the pH at which the survival fold goes below that threshold during acidic perturbation. We used two distinct definitions to choose a threshold for the survival fold. The first was a definition of “dying” where the taxa’s abundance started to decline in abundance compared to T_0_ (survival fold threshold < 1, red solid lines in Fig. S21A). The second was a definition of “dead” where the taxa’s abundance was close to 0 (survival fold threshold *→* 0, red dashed lines in Fig. S21A). For each of these definitions, the pH transition points were plotted (Fig. S21B with the first “dying” definition and Fig. S21C with the second “dead” definition) and compared to the trends of functional regime boundaries (transition from Regime II to I). Employing the ‘dying’ definition with Proteobacteria, Bacteroidota (Fig. S21B) allowed us to recapitulate the phenomenon observed in the functional data, where the fitted slope of Boundary I-II was less than 1, as shown in Fig. 6A. This suggests that these phyla in the relatively neutral soil are more tolerant of larger ΔpH change until they start to die than those in acidic soils, possibly due to variations in soil titration curves (Fig. 6B). Because the fitted slope is greater than 0, this also means that these phyla in relatively acidic soils can tolerate lower acidic pH conditions than those in neutral soils, which signals long-term pH adaptation. The ‘dead’ definition threshold resulted in a flat slope close to 0. This suggests that, despite long-term adaptation to varying native soil pH levels, these taxa have similar pH thresholds at which complete decimation occurs.

Similarly, to understand the transition to the Resurgent growth regime (Regime III), we observed the growth folds of Firmicutes phylum across perturbed pH levels. Then, we applied an identical growth fold threshold for all soils (red lines in Fig. S21A) to compute the pH at which the growth fold goes above the threshold during basic perturbations. These pH transition points were plotted (Fig. S21B&C) and compared to the trends of functional regime boundaries (transition from Regime II to III). Consistent with the trend of functional regime boundary II-III (Fig. 6A), the abundance of Firmicutes began to increase at higher pH values as the native soil pH increased. Since the NaOH amount, and consequently, the carbon nutrient level, remains constant at the pH boundary of II-III (Fig. S24), the reason Firmicutes increases at higher pH values is not linked to the amount of nutrients available. Therefore, this is another signal for taxa adaptation to long-term pH variation.

## Supplementary Tables

**Table S1:** Varying experimental conditions in previous studies.

| Paper | Soil/source | Carbon | Chloramphenicol | Water content | Measurement (time) |
| --- | --- | --- | --- | --- | --- |
| Wijler & Delwiche (1954) | 1 soil's pH modified by H<sub>2</sub>SO<sub>4</sub> or KOH | Dried alfalfa added | Untreated | Added water and dried | N<sub>2</sub>O/N<sub>2</sub> gas (17 days) |
| Nömmik (1956) | 6 soils (pH 5.2-7.6), each pH modified by HCl or KOH | Untreated, wheat straw, glucose conditions | Untreated | 100% water saturation | N<sub>2</sub>O/N<sub>2</sub> gas, NO<sub>3</sub>-, NO<sub>2</sub>-, NH<sub>4</sub><sup>+</sup> (12 days) |
| Bremner & Shaw (1958) | 8 soils (pH 3.6-8.6) and 2 of those soils (pH 3.6-4.1)'s pH modified by CaOH | Glucose | Untreated | Waterlogged (slurry) | Loss of N (30 days) |
| Valera & Alexander (1961) | 5 pure cultures of denitrifiers tested in buffered media of different pH levels | Glucose and yeast extract | Untreated | Defined buffered medium | N<sub>2</sub> gas |
| Burford & Bremner (1975) | 17 soils (pH 5.8-7.8) | Untreated | Untreated | Waterlogged (slurry) | Nitrous oxide/NO/N<sub>2</sub> gas, NO<sub>3</sub>-, NO<sub>2</sub>-, NH<sub>4</sub><sup>+</sup> (7 days) |
| Van Cleemput et al. (1975) | 1 soil's pH modified by HCl or NaOH | Dried ground plant material | Untreated | Waterlogged (slurry) | N<sub>2</sub>O/N<sub>2</sub> gas, NO<sub>3</sub>-, NO<sub>2</sub>- (100 hrs) |
| Smith et al. (1978) | 4 soils (pH 6.4-7.6) | Untreated | Chloramphenicol and untreated | Waterlogged (slurry) | N<sub>2</sub>O while acetylene inhibition (10 hrs) |
| Smith & Tiedje (1979) | 3 soils and pure cultures of denitrifiers | Succinate or glucose added and untreated | Chloramphenicol and untreated | Waterlogged (slurry) | N<sub>2</sub>O while acetylene inhibition (10 hrs) |
| Koskinen & Keeney (1982) | 4 soils (pH 4.6-6.9) | Untreated | Untreated | 20% water content | N<sub>2</sub>O/NO/N<sub>2</sub> gas, NO<sub>3</sub>-, NO<sub>2</sub>-, NH<sub>4</sub><sup>+</sup> (16 days) |
| Waring & Gilliam (1983) | 15 acidic soils (pH 3.42-5.54) and 5 of those soils limed to increase pH | Glucose and untreated | Untreated | Waterlogged (slurry) | NO<sub>3</sub>-, NO<sub>2</sub>- (20 days) |
| Parkin et al. (1985) | 2 soils (pH 4, 6), each pH modified by H<sub>2</sub>SO<sub>4</sub> or NaOH | Glucose for short-term (DEA), untreated for long-term (DP) | Chloramphenicol (DEA) and untreated (DP) | Waterlogged (slurry) for DEA | N<sub>2</sub>O while acetylene inhibition. Denitrifying enzyme activity (DEA) (2 hrs), Denitrification potential (DP) (5 days) |
| Nägele & Conrad (1990) | 3 soils (pH 4-7.8), each pH modified by HCl or NaOH | Glucose | Chloramphenicol and untreated | Waterlogged (slurry) | N<sub>2</sub>O/NO/N<sub>2</sub> gas, NO<sub>3</sub><sup>-</sup>, NO<sub>2</sub><sup>-</sup>, NH<sub>4</sub><sup>+</sup> (20 hrs) |
| Drury et al. (1991) | 13 soils (pH 4.88-7.79) | Glucose for denitrification potential, untreated for background denitrification rate | Untreated | Field capacity | N<sub>2</sub>O while acetylene inhibition (75 hrs) |
| Bandibas et al. (1994) | 18 soils (pH 4-7.8) | Untreated | Untreated | Field capacity, saturation, waterlogged | N<sub>2</sub>O/N<sub>2</sub> gas, NO<sub>3</sub><sup>-</sup>, NO<sub>2</sub><sup>-</sup>, NH<sub>4</sub><sup>+</sup> (20 days) |
| Yamulki et al. (1997) | 3 soils (pH 3.9, 5.9, 7.6) and the acid soil (pH 3.9)'s pH modified by NaOH | Untreated | Untreated | Unadjusted | N<sub>2</sub>O/NO/N<sub>2</sub> gas, NO<sub>3</sub><sup>-</sup>, NO<sub>2</sub><sup>-</sup> (6hrs), field measurements (12 months) |
| Ellis et al. (1998) | 4 soils (pH 3.3-6.1) | Untreated | Chloramphenicol and untreated | Waterlogged (slurry) for anaerobic condition and aerobic condition | N<sub>2</sub>O with and without acetylene inhibition, NO<sub>3</sub><sup>-</sup>, NO<sub>2</sub><sup>-</sup>, NH<sub>4</sub><sup>+</sup> (48 hrs) |
| Šimek et al. (2000) | 13 soils (pH 5.62-7.77) | Glucose | Chloramphenicol (DEA) and untreated (DP) | Waterlogged (slurry) | N<sub>2</sub>O while acetylene inhibition. Denitrifying enzyme activity (DEA) (30-60 min), Denitrification potential (DP) (48 hrs) |
| Simek et al. (2002) | 5 soils (pH 4.9-7.9), each pH modified by H<sub>2</sub>SO<sub>4</sub> or NaOH | Glucose | Chloramphenicol (DEA) and untreated (DP) | Waterlogged (slurry) | N<sub>2</sub>O while acetylene inhibition. Denitrifying enzyme activity (DEA) (30 min-3 hrs), Denitrification potential (DP) (48 hrs) |
| Liu et al. (2010) | 3 soils (pH 3-4), each pH modified by long-term liming | Glutamate and untreated | Untreated | Unadjusted (1st run), flooding and draining (2nd run) | N<sub>2</sub>O/N<sub>2</sub> gas (21 hrs) |
| Cuhel et al. (2010) | 3 soils (pH 5.5-7.7), each pH modified by 10-month application of H<sub>2</sub>SO<sub>4</sub> and KOH | Glucose | Chloramphenicol for short-term (DEA), untreated for in-situ field chamber measurements | Waterlogged (slurry) for DEA | N<sub>2</sub>O/N<sub>2</sub> gas, NO<sub>3</sub>-, NO<sub>2</sub>-, NH<sub>4</sub>+. In-situ field chamber measurements (74 hrs), Denitrifying enzyme activity (DEA) (30-60 min) |
| Bergaust et al. (2010) | 1 pure culture in 5 pH levels (pH 6.0-7.5) | Succinate | Untreated | Defined buffered medium | N<sub>2</sub>O/NO/N<sub>2</sub> gas, NO<sub>3</sub>-, NO<sub>2</sub>- (72 hrs) |
| Dörsch et al. (2012) | Cells extracted from 3 soils (pH 5.4-7.1) perturbed to pH 5.4 and 7.1 | Glutamate | Untreated | Defined buffered medium | N<sub>2</sub>O/NO/N<sub>2</sub> gas, NO<sub>3</sub>-, NO<sub>2</sub>- (130 hrs) |
| Samad et al. (2016) | 13 soils (pH 5.57-7.03) | Untreated | Untreated | Flooded and drained | N<sub>2</sub>O/NO/N<sub>2</sub> gas (200 hrs) |

**Table S2:**
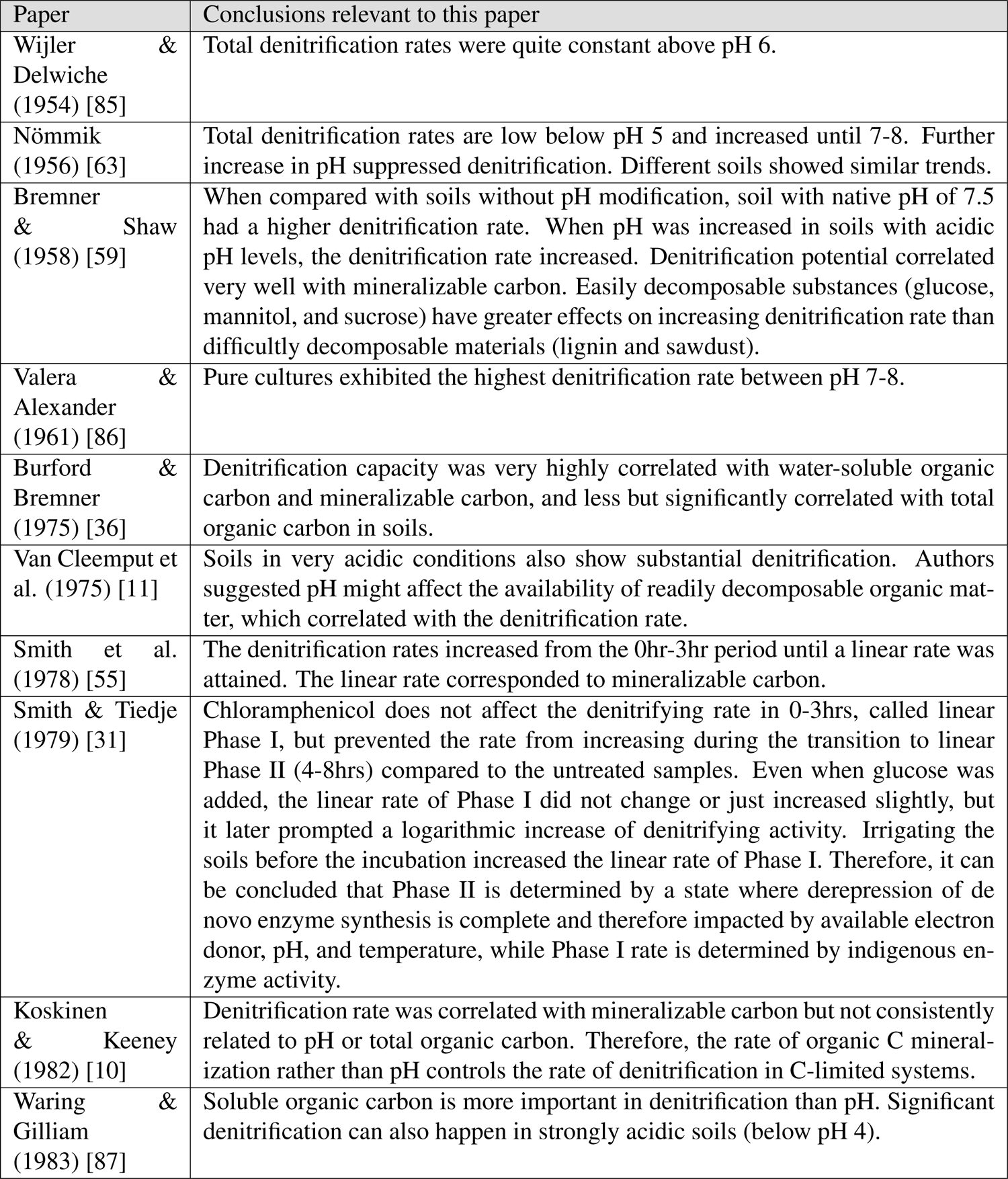
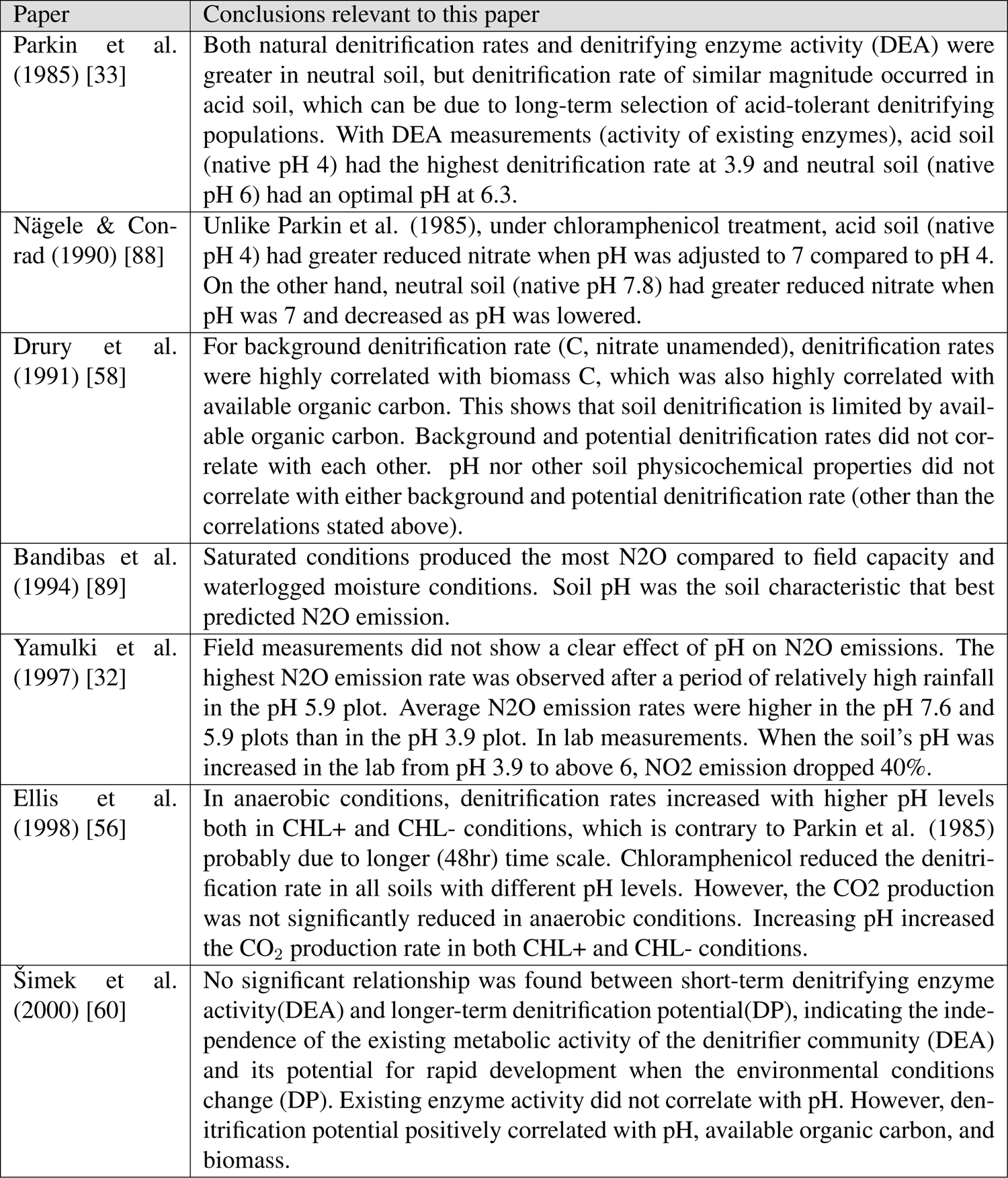
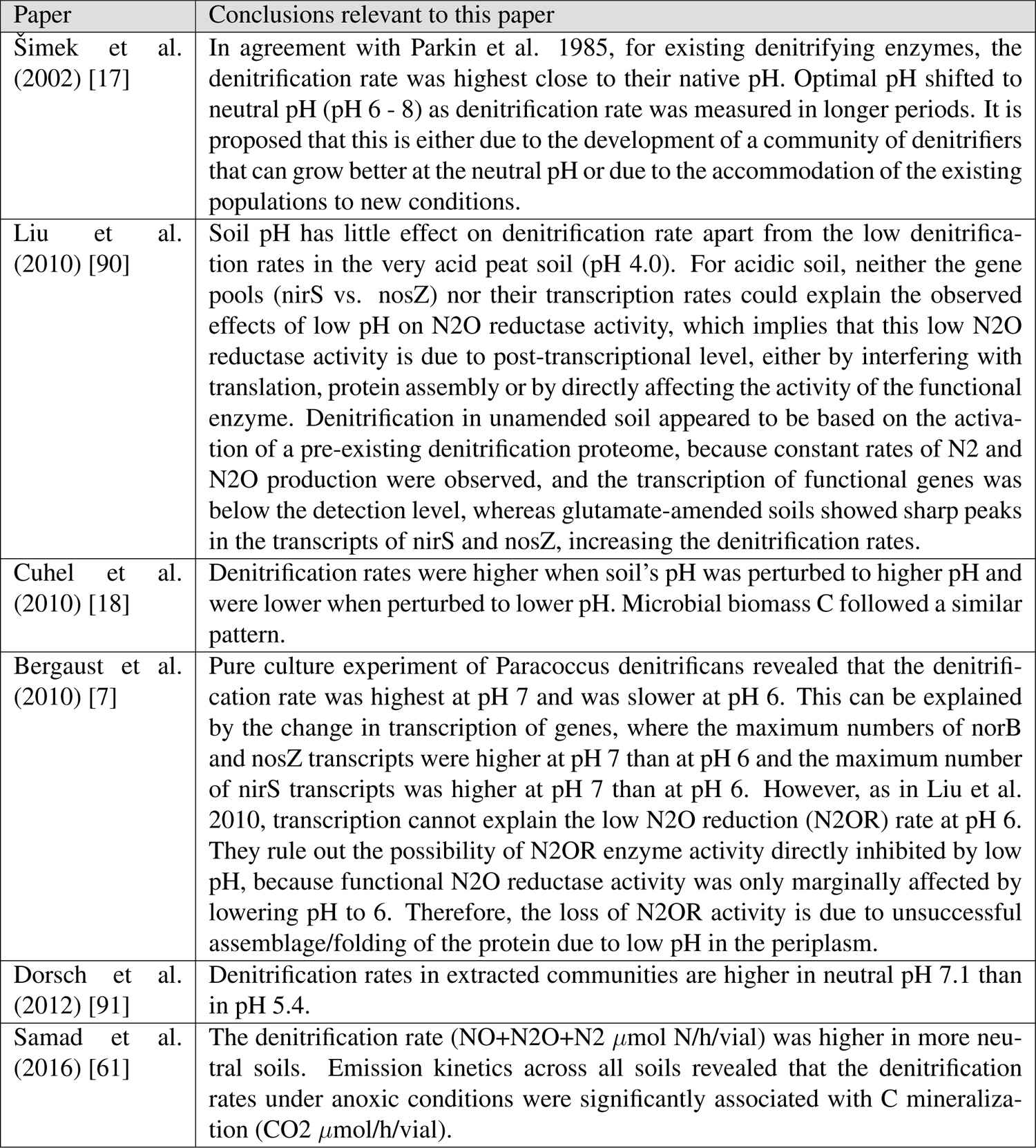
Relevant conclusions from previous studies.

**Table S3:** Physical and chemical characteristics of soil samples.

| Soil No. | Soil ID | $pH_{H_2O}$ | Latitude | Longitude | Sand (%) | Silt (%) | Clay (%) | Total N (%) | Total C (%) | C:N ratio | Depth | Date |
| --- | --- | --- | --- | --- | --- | --- | --- | --- | --- | --- | --- | --- |
| Soil1 | Acidic4 | 4.703 | 46.785703 | -117.079245 | 0.00 | 58.15 | 41.85 | 0.13 | 1.61 | 12.79 | 0-10cm | 9/10/22 |
| Soil2 | Acidic12 | 5.094 | 46.781117 | -117.080513 | 0.00 | 63.80 | 36.20 | 0.15 | 1.97 | 13.19 | 0-10cm | 9/12/22 |
| Soil3 | CE239 | 4.987 | 46.781153 | -117.080455 | 0.00 | 61.20 | 38.80 | 0.11 | 1.35 | 12.27 | 10-20cm | 9/8/22 |
| Soil4 | SE56b | 5.277 | 46.778744 | -117.082738 | 0.00 | 57.40 | 42.60 | 0.10 | 1.26 | 13.31 | 10-20cm | 9/8/22 |
| Soil5 | CE201 | 5.324 | 46.781049 | -117.086242 | 0.00 | 58.70 | 41.30 | 0.12 | 1.57 | 13.07 | 10-20cm | 9/8/22 |
| Soil6 | CE73 | 5.405 | 46.779637 | -117.086172 | 0.00 | 61.20 | 38.80 | 0.13 | 1.65 | 12.97 | 10-20cm | 9/8/22 |
| Soil7 | CE153 | 5.514 | 46.7805 | -117.085431 | 0.00 | 58.70 | 41.30 | 0.13 | 1.76 | 13.82 | 10-20cm | 9/8/22 |
| Soil8 | CE56a | 5.552 | 46.778855 | -117.082968 | 0.00 | 58.70 | 41.30 | 0.10 | 1.35 | 13.64 | 10-20cm | 9/8/22 |
| Soil9 | CE277 | 5.822 | 46.781883 | -117.0835 | 0.00 | 61.20 | 38.80 | 0.09 | 1.19 | 12.65 | 10-20cm | 9/8/22 |
| Soil10 | CE253 | 5.975 | 46.781534 | -117.084192 | 0.00 | 62.50 | 37.50 | 0.08 | 0.98 | 12.00 | 10-20cm | 9/8/22 |
| Soil11 | CE234 | 6.186 | 46.781301 | -117.082533 | 0.00 | 62.50 | 37.50 | 0.13 | 1.88 | 14.04 | 10-20cm | 9/8/22 |
| Soil12 | CE229 | 6.255 | 46.781206 | -117.084623 | 0.00 | 58.80 | 41.20 | 0.09 | 1.20 | 12.73 | 10-20cm | 9/8/22 |
| Soil13 | Neutral7 | 6.435 | 46.781523 | -117.084533 | 0.00 | 60.00 | 40.00 | 0.08 | 0.99 | 12.06 | 10-20cm | 9/11/22 |
| Soil14 | Neutral2 | 6.545 | 46.781308 | -117.084696 | 0.00 | 58.80 | 41.20 | 0.11 | 1.28 | 12.06 | 10-20cm | 9/11/22 |
| Soil15 | Neutral5 | 6.789 | 46.781416 | -117.084818 | 0.00 | 63.70 | 36.30 | 0.10 | 1.31 | 13.79 | 10-20cm | 9/11/22 |
| Soil16 | Neutral6 | 6.860 | 46.781524 | -117.084694 | 0.00 | 61.20 | 38.80 | 0.09 | 1.07 | 12.30 | 10-20cm | 9/11/22 |
| Soil17 | Neutral3 | 7.052 | 46.781194 | -117.084732 | 0.00 | 60.00 | 40.00 | 0.09 | 1.09 | 12.56 | 10-20cm | 9/11/22 |
| Soil18 | Neutral1 | 7.681 | 46.781354 | -117.084812 | 0.00 | 63.70 | 36.30 | 0.10 | 1.21 | 12.37 | 10-20cm | 9/11/22 |
| Soil19 | Neutral4 | 8.232 | 46.781222 | -117.084882 | 0.00 | 62.50 | 37.50 | 0.08 | 1.05 | 13.78 | 10-20cm | 9/11/22 |
| Soil20 | CE251 | 8.323 | 46.781492 | -117.085028 | 0.00 | 67.50 | 32.50 | 0.09 | 1.47 | 16.14 | 10-20cm | 9/8/22 |

## Supplementary Figures

**Figure S1:**
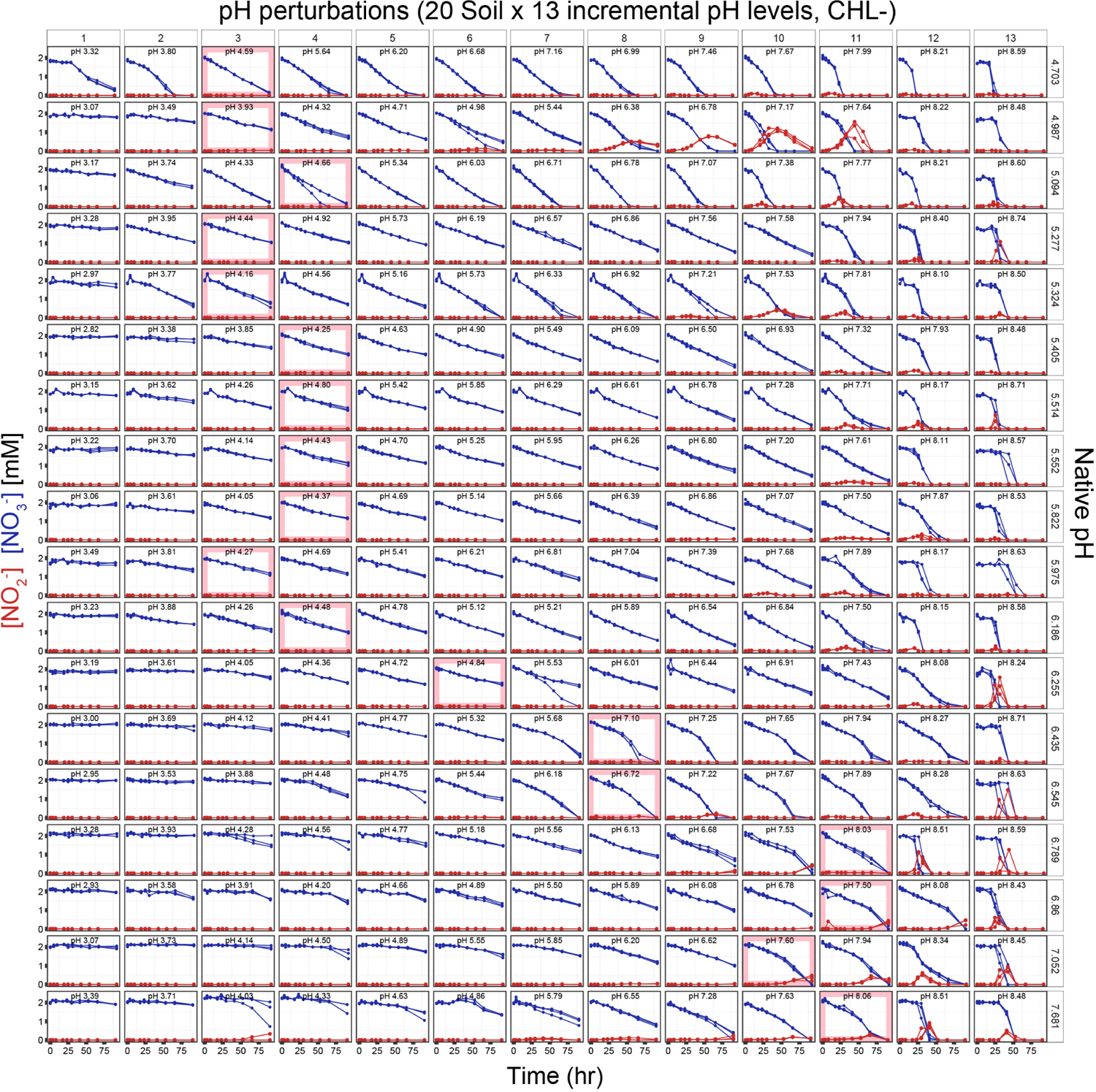
Flux dynamics of nitrate and nitrite of the dataset. Time series measurements of nitrate (blue points) and nitrite (red points) across 4 days are shown. Each row is from the identical soil sample of a native pH level (pH*_H_*_2_*_O_*), indicated at the right end of each row in the order of most acidic (top) to most basic (bottom). Each row has 13 columns which are the 13 different levels of short-term pH perturbations. The targeted perturbed pH levels were determined by constructing a soil pH titration curve before the experiment and computing how much acid (HCl) or base (NaOH) to add to the slurries. Perturbed pH levels are indicated inside each panel, which are obtained by measuring the stabilized pH values at the endpoint of the experiment (see Methods). Each line connects the point of measurements of a replicate, constituting the 3 replicates per perturbed condition. The pink-colored box for each row indicates the condition without any acid/base addition, where the pH of these conditions also changes with incubation. Soil 19 and Soil 20 are not shown due to having different numbers of perturbed pH levels (7 and 3, respectively).

**Figure S2:**
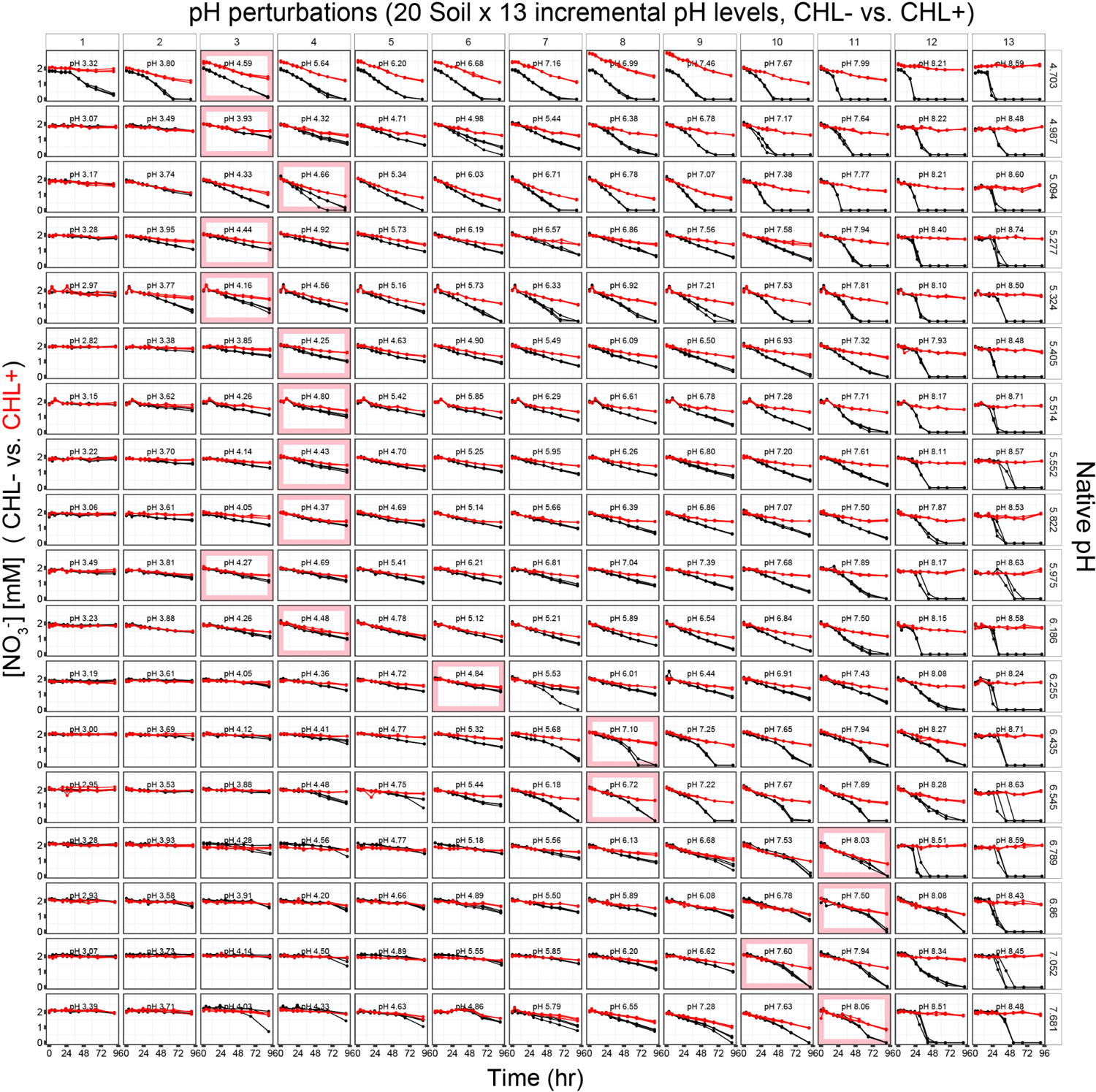
Nitrate dynamics of chloramphenicol untreated (CHL-) and treated (CHL+) conditions in the dataset. Time series measurements of nitrate in chloramphenicol-untreated (CHL-, black points) and treated (CHL+, red points) across 4 days are shown. Each row is from the identical soil sample of a native pH level (pH*_H_*_2_*_O_*), indicated at the right end of each row in the order of most acidic (top) to most basic (bottom). Each row has 13 columns which are the 13 different levels of short-term pH perturbations. The targeted perturbed pH levels were determined by constructing a soil pH titration curve before the experiment and computing how much acid (HCl) or base (NaOH) to add to the slurries. Perturbed pH levels are indicated inside each panel, which are obtained by measuring the stabilized pH values at the endpoint of the experiment (see Methods). Each line connects the point of measurements of a replicate, constituting the 3 replicates per perturbed condition. The pink-colored box for each row indicates the condition without any acid/base addition, where the pH of these conditions also changes with incubation. Soil19 and Soil20 are not shown due to having different numbers of perturbed pH levels (7 and 3, respectively).

**Figure S3:**
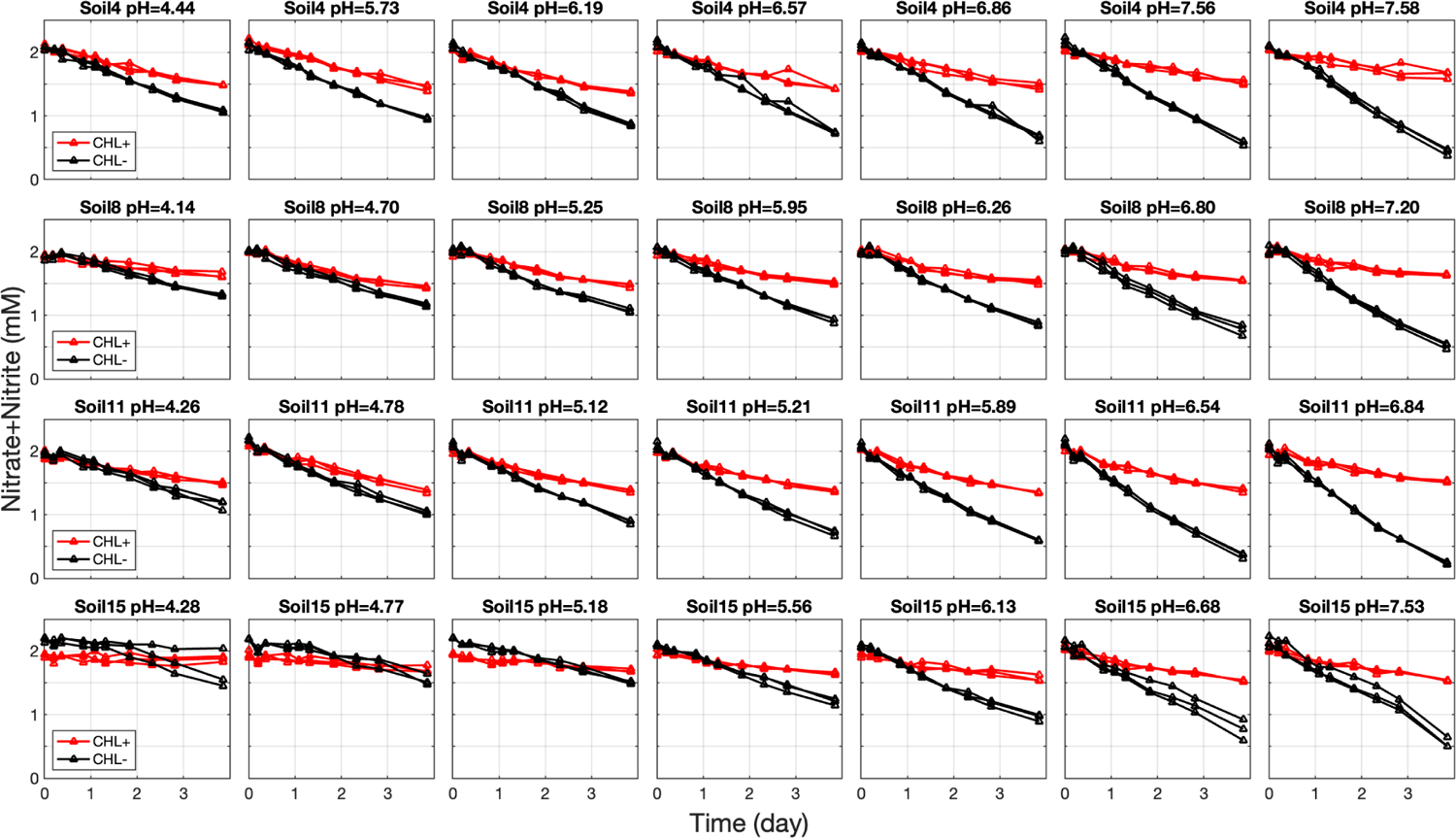
Nitrate + nitrite concentration dynamics to show constant nitrite reduction rates. The points indicate the time-series measurement of the sum of nitrate and nitrite concentrations. Concentrations from chloramphenicol-treated (CHL+) samples are in red and untreated (CHL-) samples are in black, with lines connecting each of the three biological replicates. A subset of pH perturbed conditions (each row is from the same native pH soil, with varying perturbed pH levels) is shown. Nitrate + nitrite dynamics (in CHL- conditions) are linear, indicating that the community’s nitrite consumption is constant across time.

**Figure S4:**
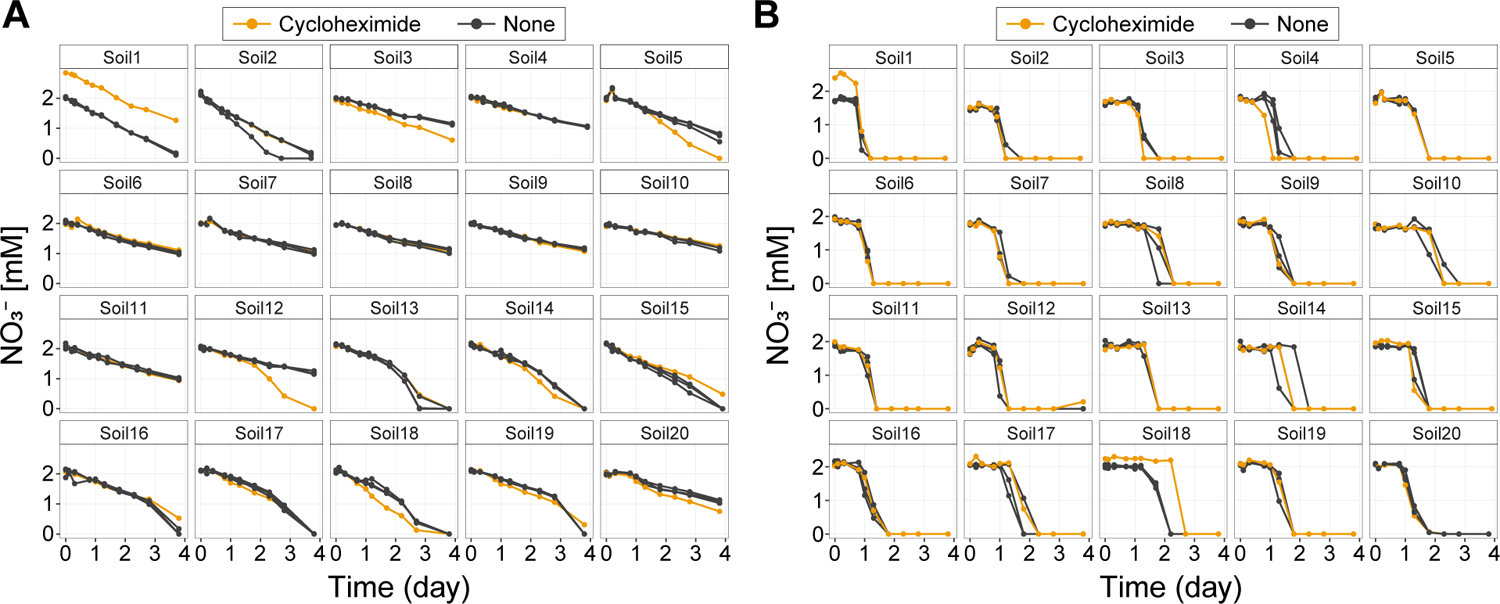
Cycloheximide antifungal controls suggest a minimal role for fungi in nitrate utilization dynamics. Nitrate dynamics across a 4-day anaerobic incubation with and without cycloheximide treatment for all 20 soils. Panel **(A)** shows the nitrate dynamics of pH-unperturbed samples with (orange data points, 1 replicate) and without cycloheximide (black data points, 3 biological replicates) treatment, while **(B)** illustrates the nitrate dynamics for basic-perturbed samples, also with (orange, 1 replicate) and without cycloheximide (black, 3 biological replicates) treatment. Most of the dynamics were not affected by the application of 200 ppm cycloheximide. Only 1 case out of 40 samples (Soil 18 in **B**) showed delayed nitrate reduction when the antifungal was treated. This means that fungi do not play a significant role during nitrate reduction.

**Figure S5:**
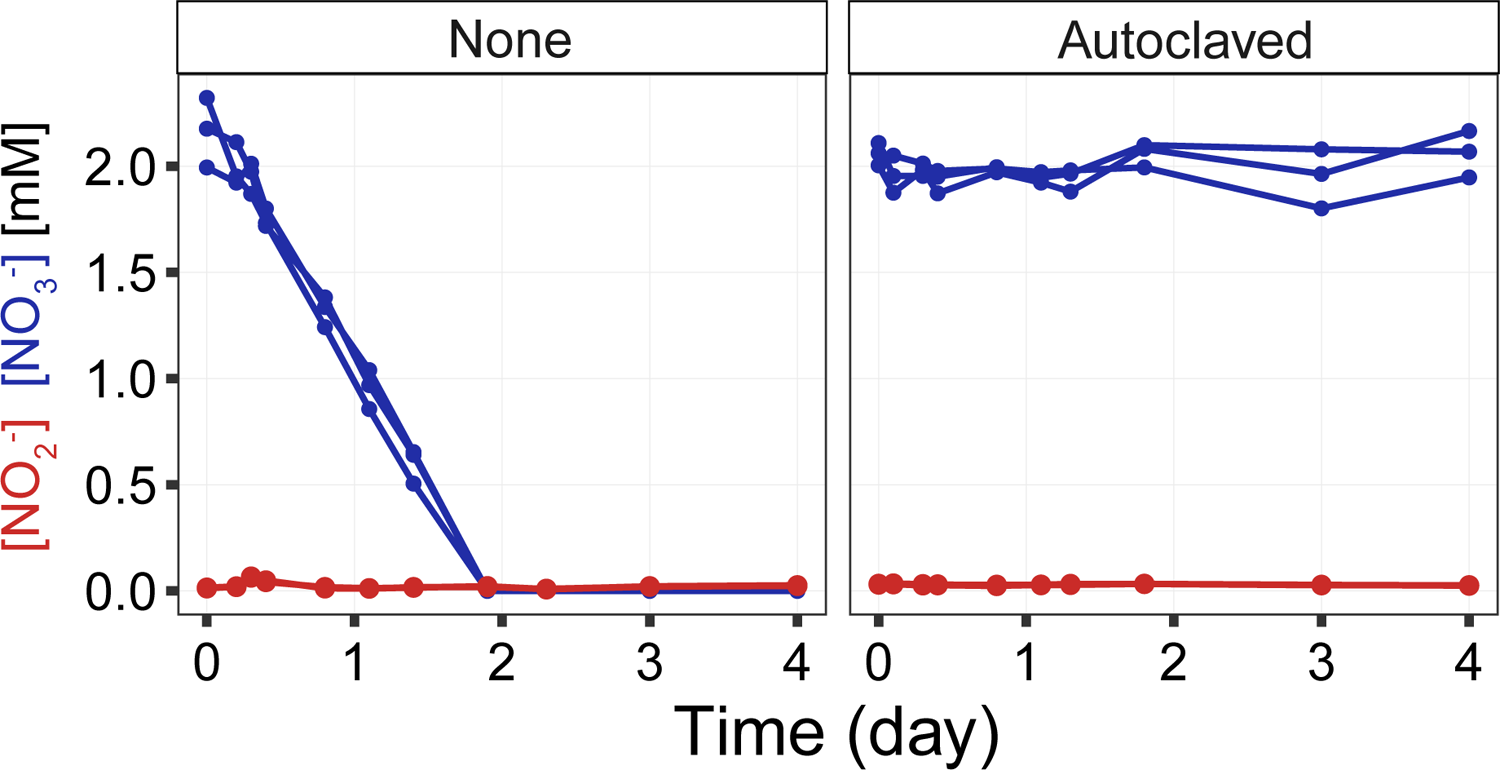
The results from autoclaving soil suggest the absence of abiotic (chemical) nitrate reduction. Nitrate (blue data points) and nitrite (red data points) dynamics of a soil sample with (right, Autoclaved) and without (left, None) autoclaving procedure. The autoclaving was performed at 120 *^◦^*C for 99 minutes and repeated five times at two-day intervals. The soil used in this experiment was collected from LaBagh Woods (latitude 41.977855, longitude −87.742585), Sauganash Prairie, Chicago, IL, USA, on January 18, 2022. Contrary to the soil without the sterilization, nitrate reduction did not occur in the soil with the sterilization process (autoclaving). This rules out the possibility of abiotic (chemical) nitrate reduction occurring in soils.

**Figure S6:**
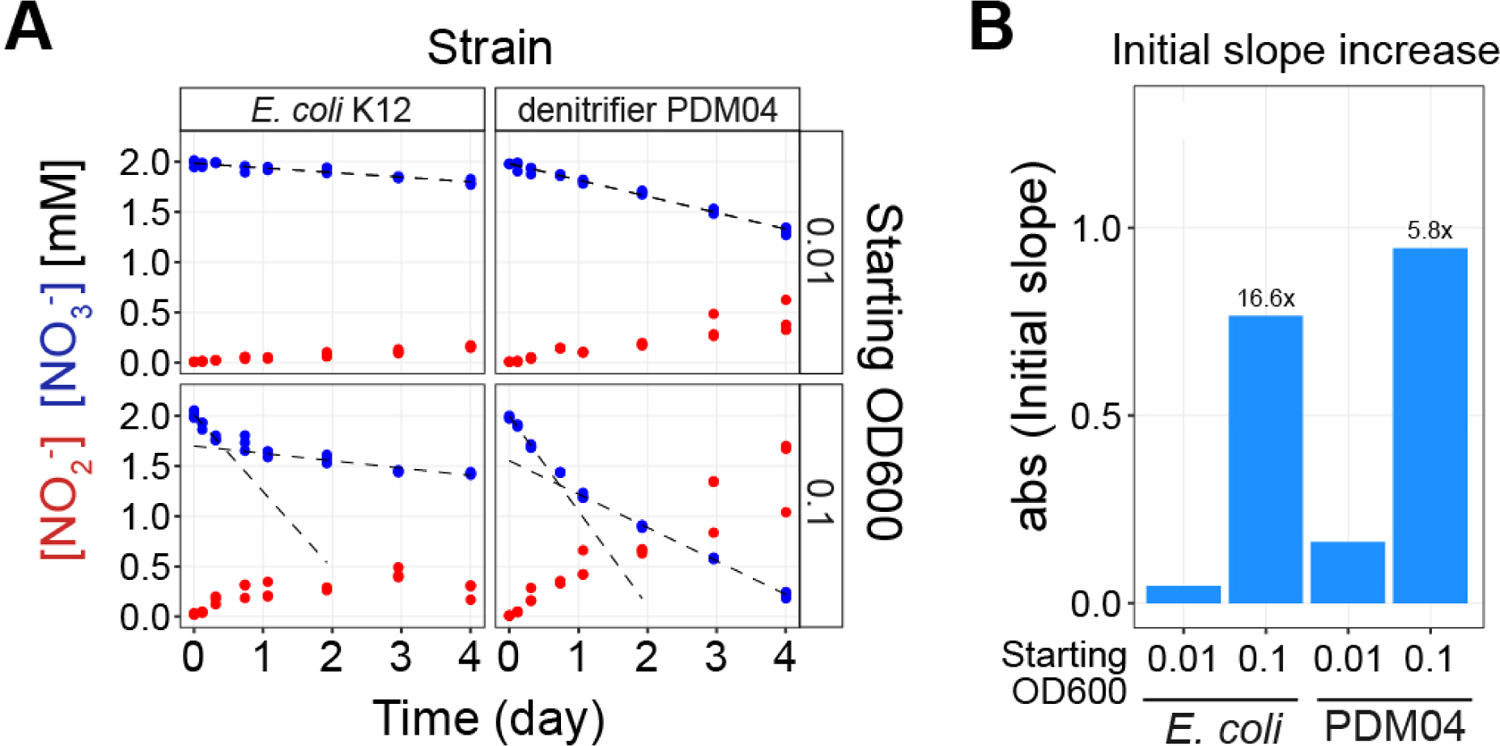
Linear metabolite dynamics recapitulated from monoculture experiments in the absence of external carbon sources. **(A)** Nitrate and nitrite dynamics of monoculture experiments using *E. coli* K12 and the denitrifier *Pseudomonas sp.* PDM04 strains over 4 days with no external carbon provided in the culture media (see SM for detailed experimental methods). The top two panels have a starting OD600 (optical density at 600nm) of 0.01 and the bottom panels have a starting OD600 (optical density at 600nm) of 0.1. The x-axis represents time in days, and the y-axis represents the concentration of nitrate (blue points) or nitrite (red points) reduced (mM), each condition having three biological replicates. The linear dynamics demonstrate that nitrate reduction can occur even in the absence of external carbon, resolving the previous contradiction about the necessity of carbon for this process. The dashed lines represent linear regression of the dynamics: for the top panels, linear regression used all data points, while in the bottom panels, initial slopes were derived from fitting the first three points, and late slopes were calculated using the last four points. **(B)** The fitted initial slope values in nitrate reduction (using initial slopes in **A**). The different bars indicate the initial slopes from different conditions of starting OD600 values (0.01 and 0.1) and two strains. The plot underscores the effect of starting biomass on initial nitrate reduction rates, with factor increase of initial slopes annotated on top of the bars.

**Figure S7:**
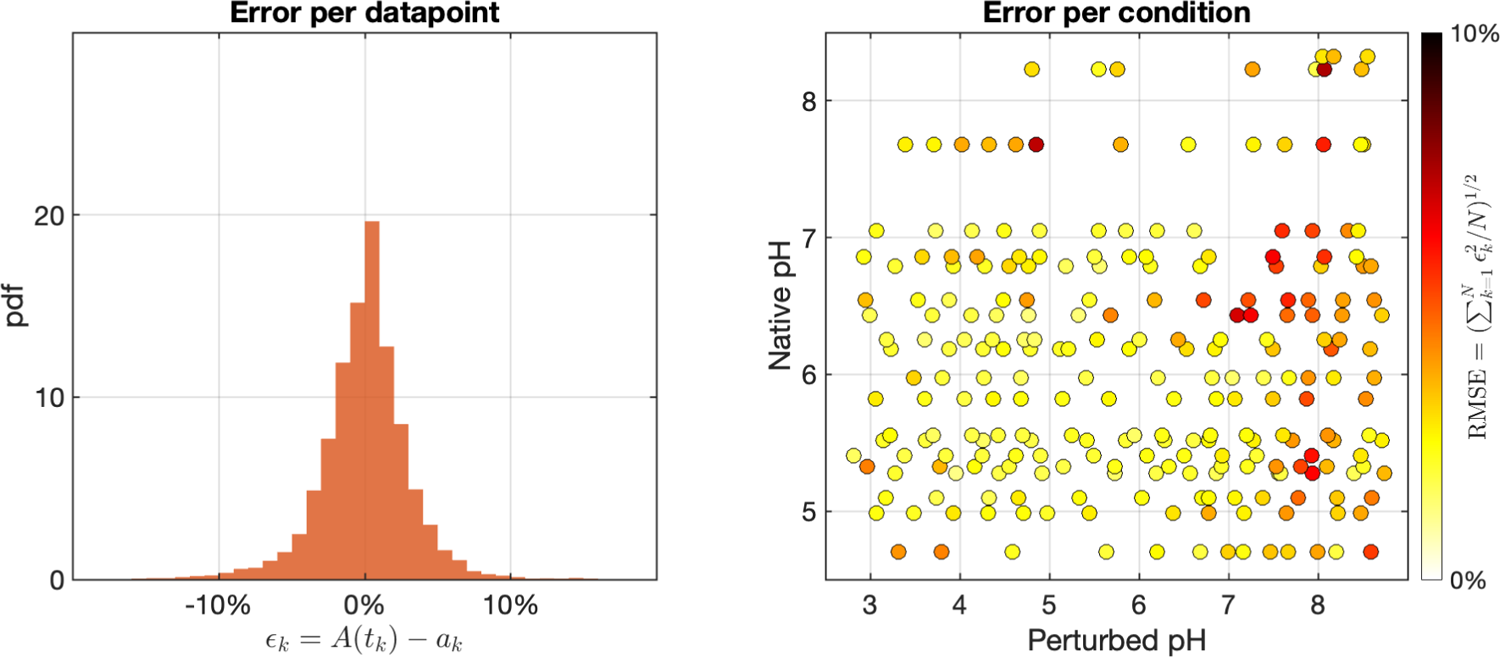
Quantification of error in model fitting. Error per data point (Left panel): The probability density function (pdf) represents the distribution of errors for individual data points of nitrate measurements at time point *k*. Errors are calculated as the difference between the model’s predicted nitrate concentration *A*(*t_k_*) and the observed nitrate amounts *a_k_* for either the chloramphenicol-untreated(CHL-) or treated(CHL+) conditions, normalized by dividing by the input nitrate concentration (2mM) to be expressed as a percentage. Error per condition (right panel): Each dot represents the error for a specific experimental condition (triplicates), with the native pH of the sample on the y-axis and the perturbed pH on the x-axis. The error per condition, indicated by the color of each point, is the square root of the mean-squared error (MSD) loss function minimized during parameter optimization of both CHL-/+ conditions of triplicates, normalized by the input nitrate concentration (2mM) to be expressed as a percentage (refer to Methods for the error computation).

**Figure S8:**
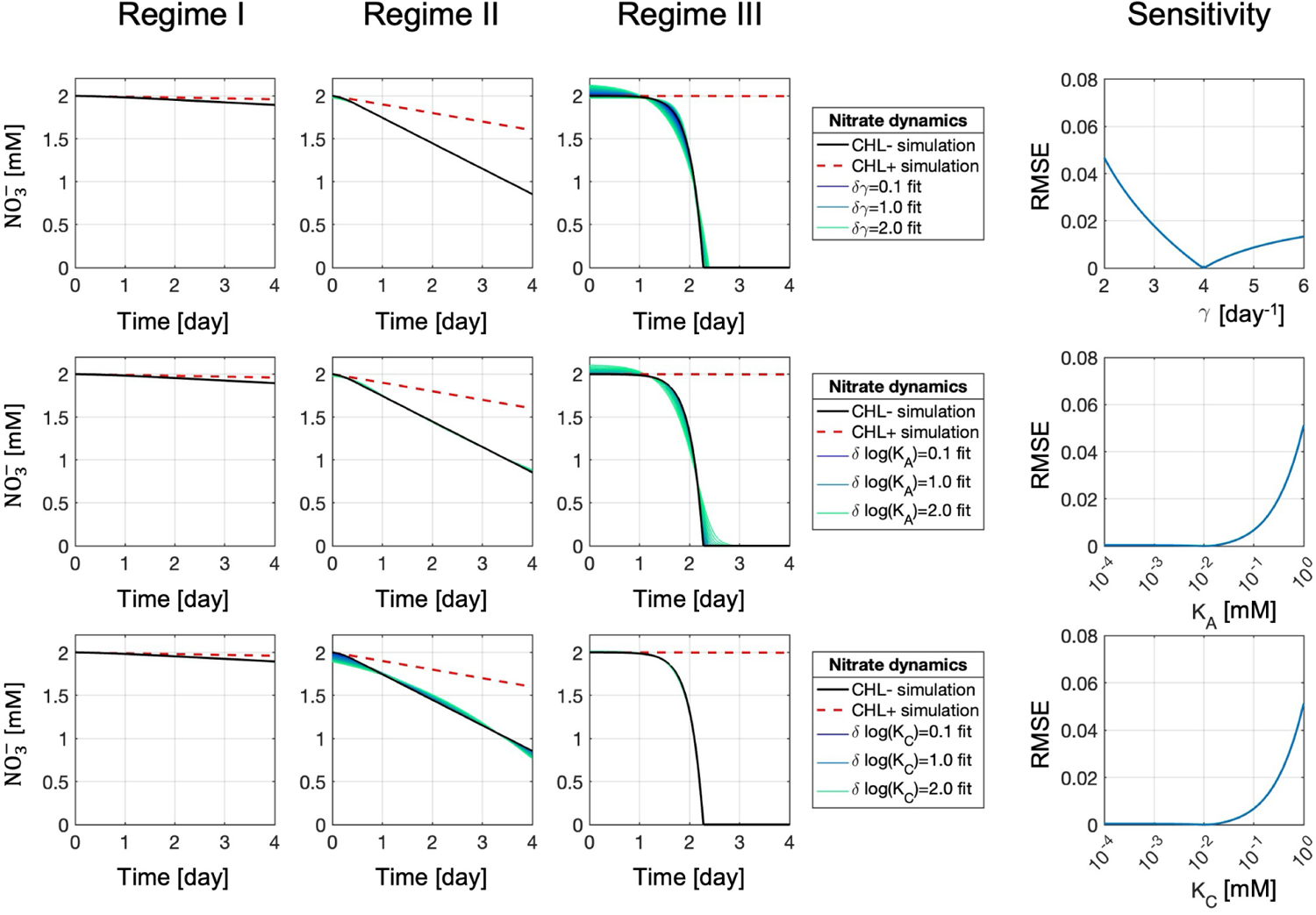
Sensitivity analysis on model parameters. *γ*, *K_A_***, and** *K*^^^*_C_***to justify fixing these parameters** To justify the fixed parameters in the fitting scheme, we analyzed the sensitivity of *γ*, *K_A_*, and *K*^^^*_C_*by simulating dynamic data. To reflect the three typical dynamics (regimes) observed from the measurement, we simulated three nitrate curves by setting up the initial conditions to be *x*^(0) = 0.01, 0.1, 0.001*mM/day* and *C*^^^(0) = 0.005, 0.05, 2*mM*, respectively. Other parameters are given by *A*_0_ = *A^c^* = 2*mM*, *K_A_* =*K*^^^*_C_*= 0.01*mM*, *γ* = 4*day^−^*^1^. Black curves indicate the simulated nitrate dynamics from the chloramphenicol-untreated (CHL-) conditions, and red dashed lines indicate the simulated nitrate dynamics from the chloramphenicol-treated (CHL+) conditions. We then used different fixed values of parameters to fit the three examples. In the first row, we used different fixed *γ* values from *γ* = 2*day^−^*^1^ to *γ* = 6*day^−^*^1^ - to fit three simulations. The square root of the mean-squared error (RMSE) is computed by the loss function (mean-squared difference of predicted and observed nitrate concentration for both CHL-/+ conditions) minimized during parameter optimization, normalized by the input nitrate concentration (2mM) to be expressed as a percentage (refer to Methods for loss function). We demonstrate very small mismatches (RMSE*<* 5%) from these variations of parameter values, which are almost invisible in Regime I and Regime II fittings (purple lines indicate fitted results from *γ* = 4 *±* 0.1 *day^−^*^1^, blue lines indicate fitted results from *γ* = 4 *±* 1 *day^−^*^1^, green lines indicate fitted results from *γ* = 4 *±* 2 *day^−^*^1^). In the second and the third row, we used different fixed *K_A_* and *K*^^^*_C_*values from 10*^−^*^4^*mM* to 1*mM* to fit three simulations. When *K_A_ <* 0.1*mM* or *K*^^^*_C_<* 0.1*mM*, the mismatches were again very small (RMSE *<* 1%) and invisible (purple lines indicate fitted results from *K_A,C_* = 10*^−^*^2^*^±^*^0.1^*mM*, blue lines indicate fitted results from *K_A,C_*= 10*^−^*^2^*^±^*^1^*mM*, and green lines indicate fitted results from *K_A,C_* = 10*^−^*^2^*^±^*^2^*mM*). These results indicate that the fixed values of *γ*, *K_A_* and *K*^^^*_C_*are insensitive in large ranges.

**Figure S9:**
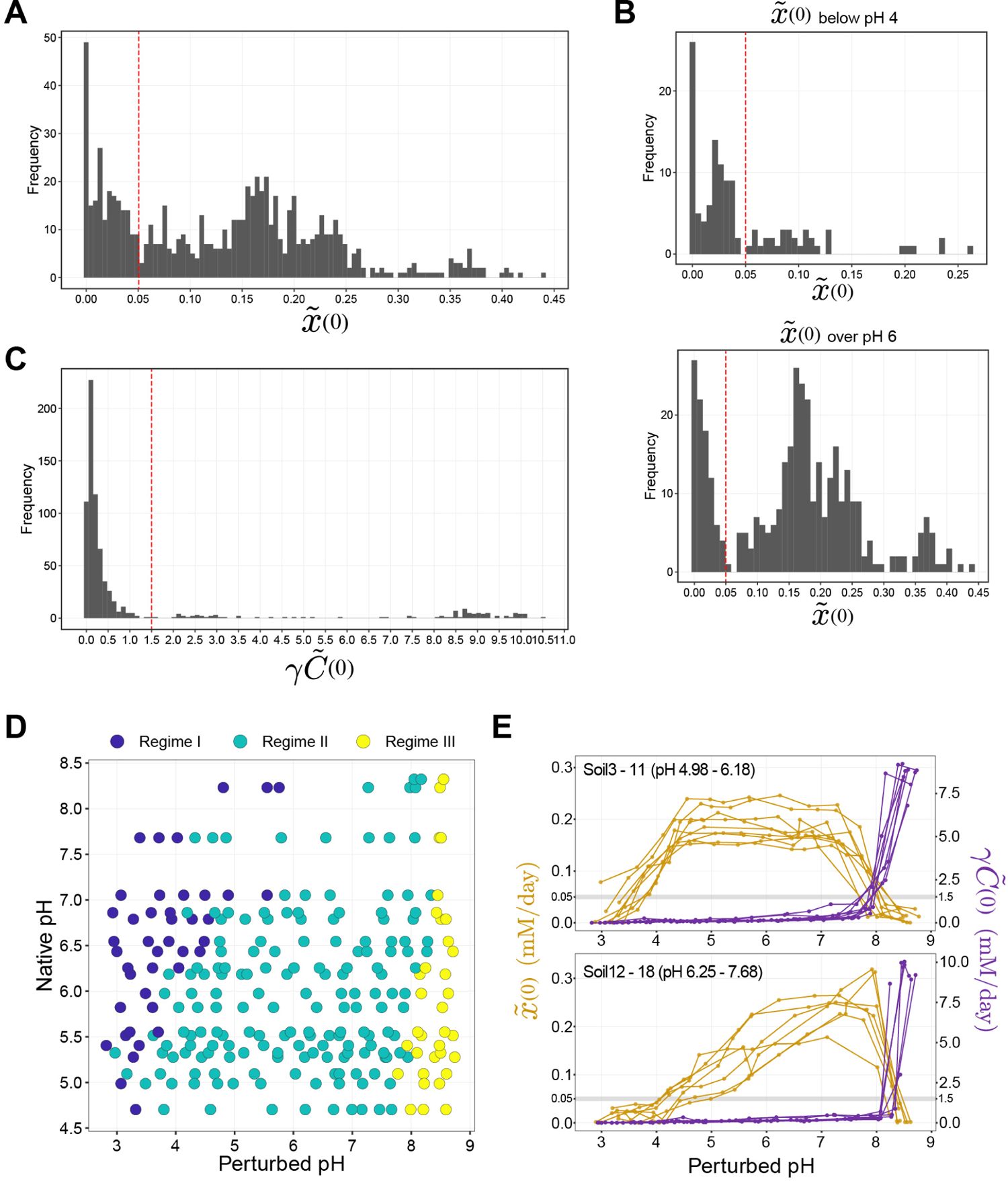
Determining regime boundary thresholds with distributions of the parameters. *x*^(0) **and** *γC*^^^(0) To determine the regime boundaries, we examined the distributions of parameters fitted to the functional data for *x*^(0) and *γC*^^^(0). **(A)** x^(0) had a bimodal frequency distribution, having two peaks. **(B)** This bi-modality becomes more evident when we separately observe its distribution from the left half (perturbed pH < 4) and right half (perturbed pH > 6) of the parameter space displayed in the perturbed pH vs. native pH grid in Figure 3C. We set the threshold for the x^(0) boundary where these two modes are separated (x^(0) = 0.05). **(C)** γC^^^(0) showed an uni-modal frequency distribution. We set the threshold (γC^^^(0) = 1.5) at the tail of the distribution, where the γC^^^(0) threshold also separated the Regime III samples in the top-left quadrant of the x^(0) vs. γC^^^(0) scatter plot (Fig. 3A). The separation of Regime I and Regime II data points may not be clear cut in the x^(0) vs. γC^^^(0) scatter plot (Fig. 3A). However, when we plot x^(0) of different soils (dark yellow colored lines) by grouping them into relatively acidic (Soil3–11 (pH 4.98-6.18), **(E)** top panel) and neutral soils (Soil12–18 (pH 6.25-7.68), **(E)** bottom panel), the transition from Regime II (large x^(0)) to Regime I (small x^(0)) is evident going towards more acidic pH perturbations, especially in the naturally acidic soils (top panel), because the large x^(0) levels are sustained over a wide pH range in Regime II. **(D)** With these thresholds of two parameters, we can define the three different regimes of adaptive behavior across native pH and perturbed conditions (colored differently by regimes).

**Figure S10:**
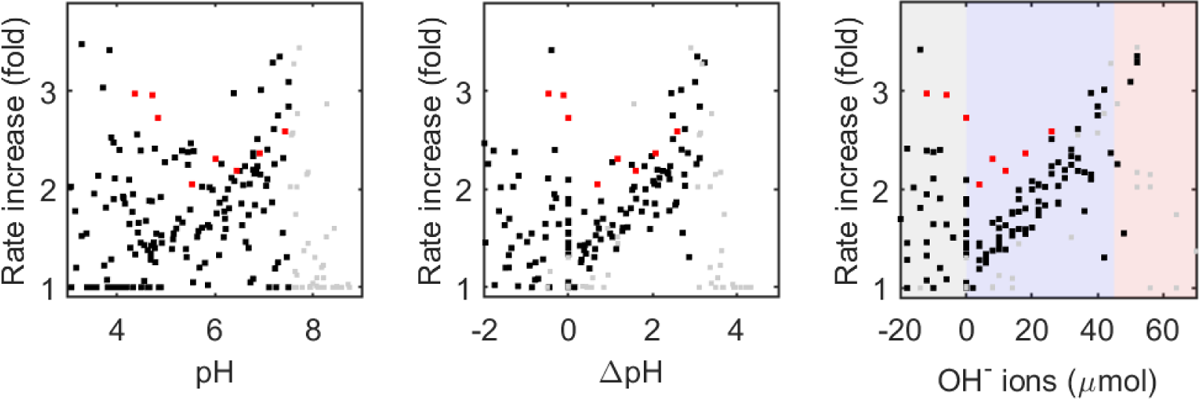
NaOH input has a more consistent linear relationship with rate increase than pH or delta pH. To provide additional evidence that the NO*^−^* reduction rate increase (fold) has a linear relationship with the added base, we calculated the rate increase (y-axis) independently from the model. To do so, we performed linear regression on the linear nitrate dynamics of chloramphenicol treated (CHL+) and untreated (CHL-) conditions, determining the slope ratio (CHL-/CHL+). On the leftmost plot, the rate increase is plotted against the perturbed pH (x-axis), against delta pH (= perturbed pH native pH) in the central plot, and against the added amount of OH*^−^* ions (in *µ* moles, negative values indicate the amount of H^+^ ions) input on the rightmost plot. We are using perturbed samples from all soils with varying native pH levels. As we progress from left to right plots, we observe a greater collapse of data into a linear relationship with the rate increase. This confirms that NaOH is the most reliable descriptor for consistently explaining the growth due to nutrient release across soils with various native pH levels. Data points from Soil12 are colored in red owing to its slope being different from the collapsed slope of other soils (black points). Treatments with pH greater than 7.5 were colored gray, as they predominantly belong to the Resurgent growth regime (Regime III), while linearity is expected to only hold in the Nuitrient-limiting regime (Regime II). The blue background in the rightmost plot is a guide for the eye, indicating the range of perturbations that typically remain within the Nutrient-limiting regime (Regime II).

**Figure S11:**
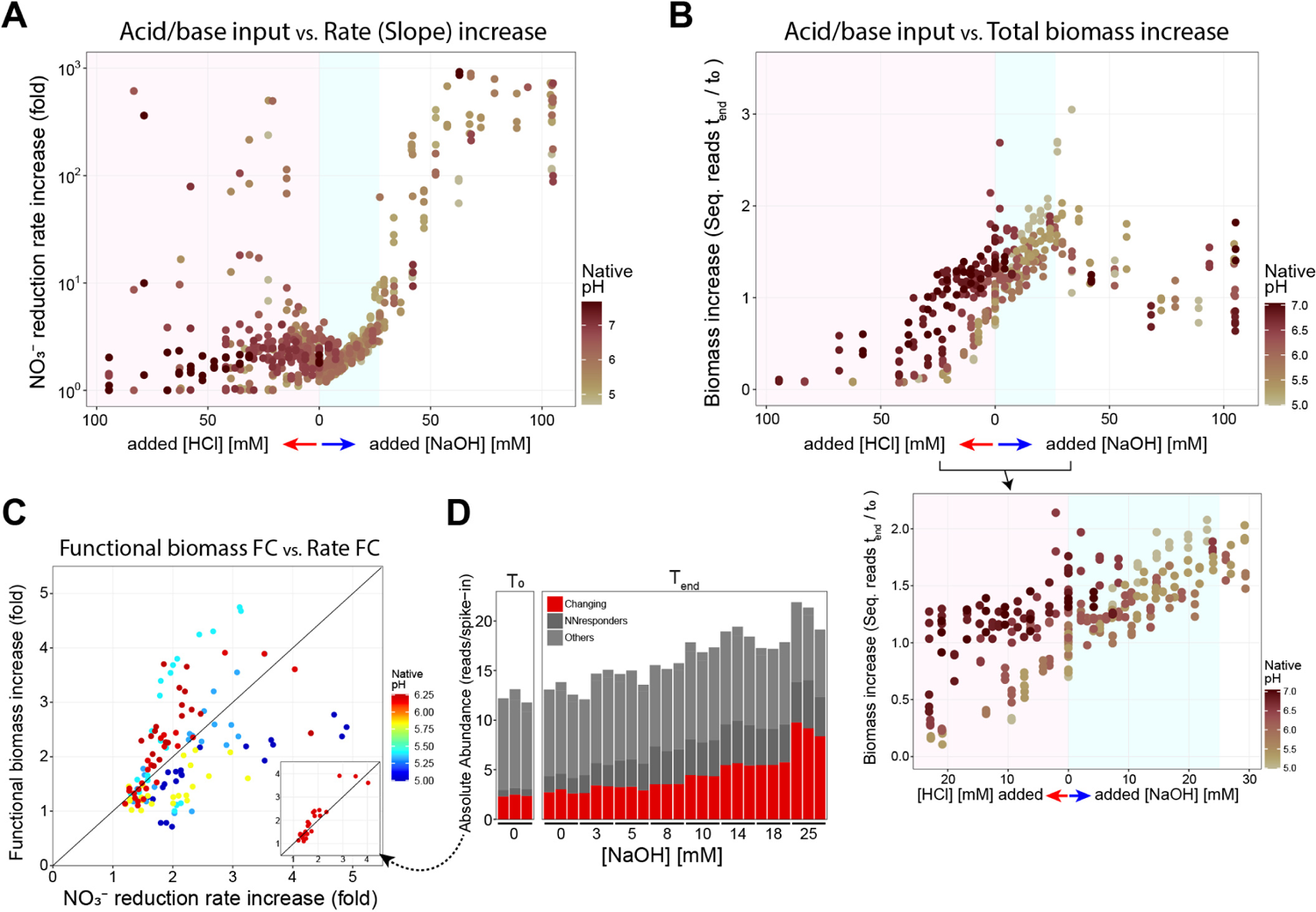
Confirming the linear dependence between functional biomass and acid/base added. A more detailed analysis, accounting for individual Amplicon sequence variant) ASVs **(C, D)**that responded to the amendment of nitrate, further confirmed the linear dependence between biomass and acid/base added. **(A)** Showing the full range of acid/base input (x-axis) against the NO*^−^* reduction rate increase (fold) (y-axis) in chloramphenicol-untreated (CHL-) conditions compared to treated conditions (CHL+) for all soils from different native pH (color gradient of data points). The rate fold increase is computed from the fitted model parameters (1 + *γC*^^^_0_*/x*^_0_). This linear relationship is observed within the range of NaOH addition from 0mM to 25mM, which belongs to the Nutrient-limiting regime (Regime II) (light blue background in **A**). This was not the case for acidic perturbations (*>* 0mM HCl addition) and basic perturbations beyond 25mM NaOH addition. Therefore, the fitted model parameter suggests that the addition of NaOH causes the release of limiting nutrients in the soil, increasing biomass growth. **(B)** Showing the full range of acid/base input against biomass growth measured by the sequencing data. Biomass increase (fold) was computed with the ratio of the total absolute abundance of initial and end time points samples (*T_end_/T*_0_). We plotted an inset to highlight a zoomed-in range (*<* 25mM HCl, *<* 25mM NaOH). In this range, the amount of biomass growth evidently increases with the addition of NaOH (light blue background) and decreases with the addition of HCl (pink background) for all soils from different native pH levels (color gradient of data points). Although this linear relationship corroborates our proposed nutrient release mechanism, to be more precise, we need to prove further that the factor increase of the “functional” biomass equals the factor increase of nitrate reduction linear rate from the flux dynamics data. This is because not all biomass performs NO*^−^* reduction. To detect the fractional biomass that performs denitrification, we used a differential abundance analysis to statistically determine which ASVs were significantly enriched in each pH perturbed condition compared to the CHL+ counterpart serving as a baseline of no growth (see Methods). We filtered out the ASVs that could be false-positive nitrate reducers by removing ASVs that were statistically enriched in no-nitrate conditions (dark grey NNresponders bar in **(D)**). Then, we summed up the absolute abundance of these ASVs that we inferred as true nitrate reducer biomass to obtain the functional biomass for each condition (red bar in **(D)**). **(C)** By comparing the factor increase of these functional biomass values (endpoint/initial functional biomass), we showed that indeed the functional biomass increase and denitrification rate increase are aligned in different soils (color spectrum in soils with different native pH). Some soils had these two values lie very close to the 1:1 diagonal line (Soil11, inset of **(C)**), which validates our inference procedure.

**Figure S12:**
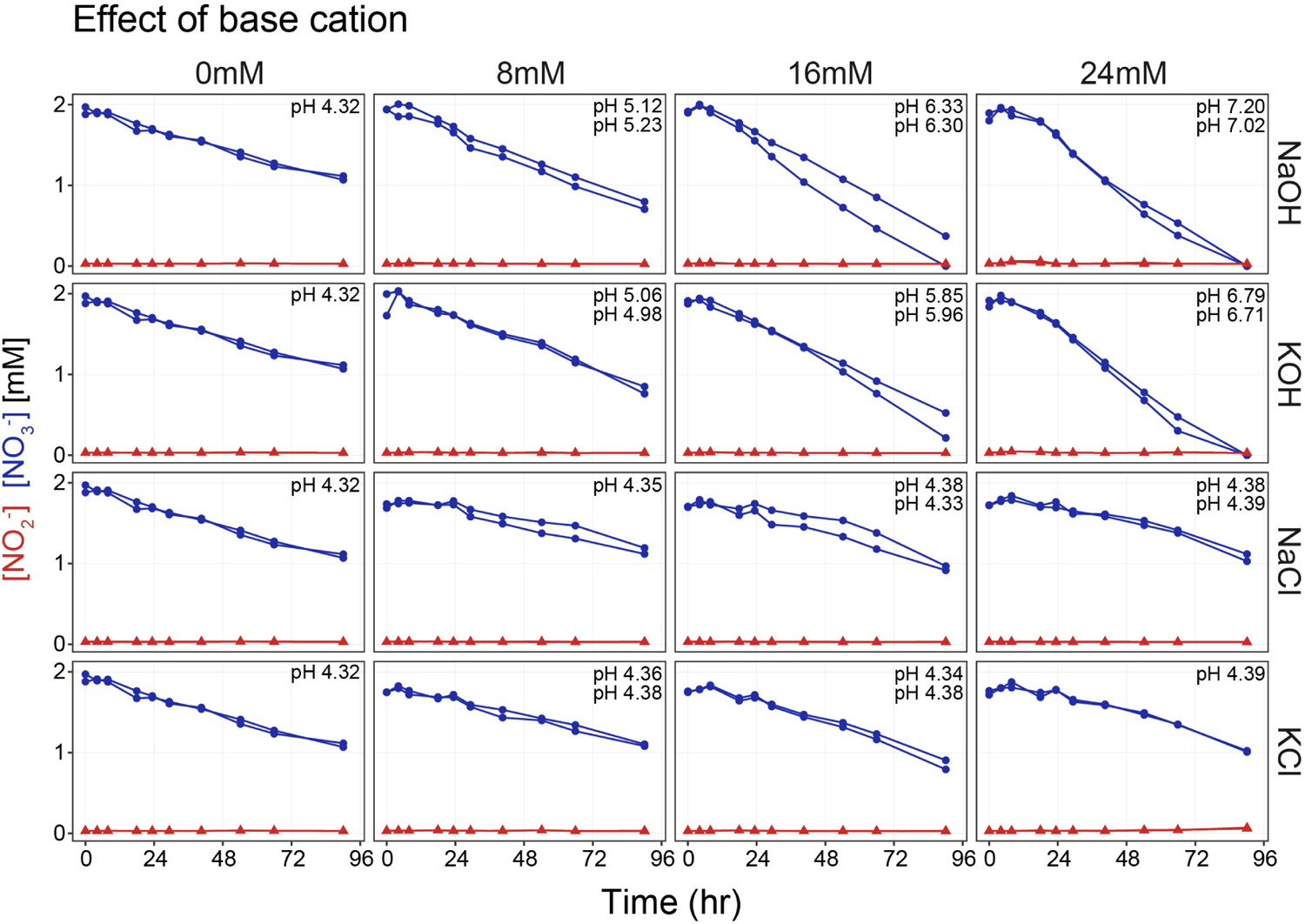
Testing the effect of different bases and salts on nutrient release. To see the effects of different bases (NaOH and KOH) on nitrate reduction dynamics, we added different concentrations of NaOH and KOH (final concentration of 0, 8, 16, 24mM in the slurry), following the same protocol previously described (without chloramphenicol), to measure the nitrate and nitrite dynamics using Soil6 (Table S3). In addition, to test the effects of Na^+^, K^+^, and Cl*^−^* separately, we added different concentrations of salts (NaCl, KCl) (without chloramphenicol and without adding any acid/base) and measured the metabolite dynamics. Blue points denote nitrate measurements and red points denote nitrite measurements. The lines connect the data points of two biological replicates. Each panel displays the stabilized endpoint pH (1M KCl method). Identical pH values across biological replicates are noted once.

**Figure S13:**
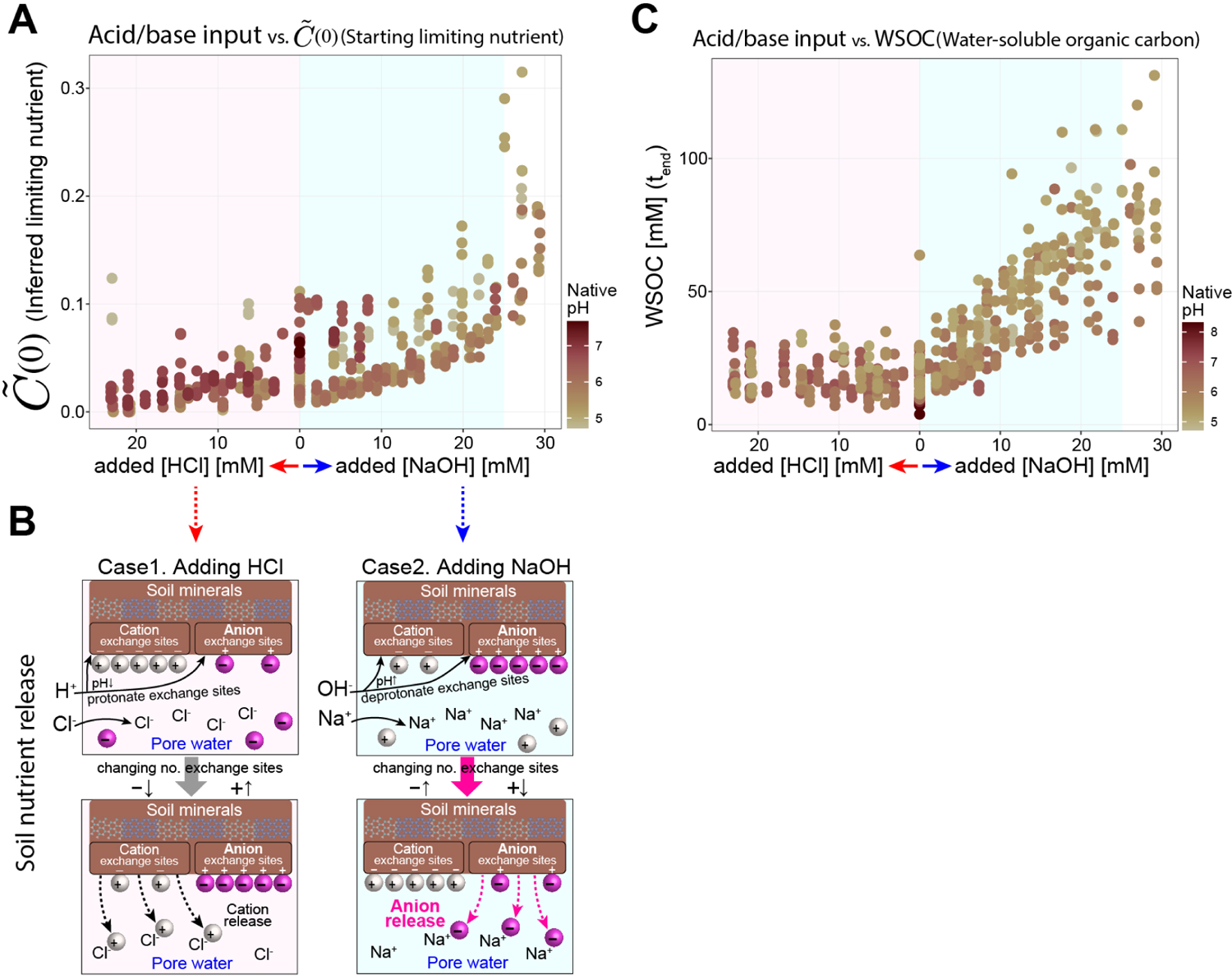
Water-soluble organic carbon (WSOC) measurement aligns with the nutrient release hypothesis in the Nutrient-limiting regime (Regime II) **(B)** Cartoon illustrating the soil nutrient release hypothesis; NaOH results in the release of anion nutrients from soil clay particles (brown region), while the addition of HCl releases cation nutrients and adsorbs anion nutrients. Microbes cannot access the nutrients adsorbed in the soil particles but can access the nutrients dissolved in soil pore water. Added OH*^−^* ions deprotonate both cation and anion exchange sites, hence decreasing the number of anion exchange sites in the soil particles and increasing the number of cation exchange sites. This releases anion nutrients from the clay particles to the pore water, while cations in the pore water are adsorbed to the clay particles. In concert, added Na^+^ ions stabilize the released anions in the pore water facilitating the release. On the other hand, during HCl addition, Added H^+^ protonates both cation and anion exchange sites, hence increasing the number of anion exchange sites in the soil particles and decreasing the number of cation exchange sites. This releases cations from the clay particles to the pore water, while anion nutrients in the pore water are adsorbed to the clay particles no longer available to the microbes. In concert, added Cl*^−^* ions stabilize the released cation in the pore water. **(A)** With this proposed mechanism of nutrient release by NaOH and HCl, we can further specify the type of growth-limiting nutrient by observing the change of the fitted model parameter of C^^^(0) (starting limiting nutrient). In natively acidic soils, increasing NaOH concentrations linearly increased the C^^^(0) (light blue region), which indicated that the limiting nutrient is negatively charged (anion nutrient). In natively neutral soils, increasing HCl concentration linearly decreased the C^^^(0) (light pink region). This is congruent with our statement that the growth-limiting nutrients are anions, because when HCl is added, anions are sequestered to the clay particles becoming unavailable to the microbes (purple spheres in **B**). **(C)** Coincidentally, adding NaOH linearly increased the water-soluble organic carbon (WSOC) concentrations present in the slurry at the endpoint, while adding HCl did not. This suggests two aspects related to our nutrient release hypothesis. Firstly, it appears that most water-soluble organic carbon (WSOC) may be negatively charged (anion). Secondly, the growth-limiting nutrient might be either the WSOC itself or another nutrient that is concomitantly released with organic carbon in the form of organic matter, including all carbon (C), nitrogen (N), sulfur (S), and phosphorus (P). If the limiting nutrient were WSOC, only a fraction of WSOC would be used as nutrient, because while C^^^(0) and WSOC are well correlated, the concentration of released WSOC is disproportionately higher (*≈*20-75 C mM) than the amount of limiting nutrient needed to deplete all 2mM NO*^−^* in the system, as shown in Fig. 4D).

**Figure S14:**
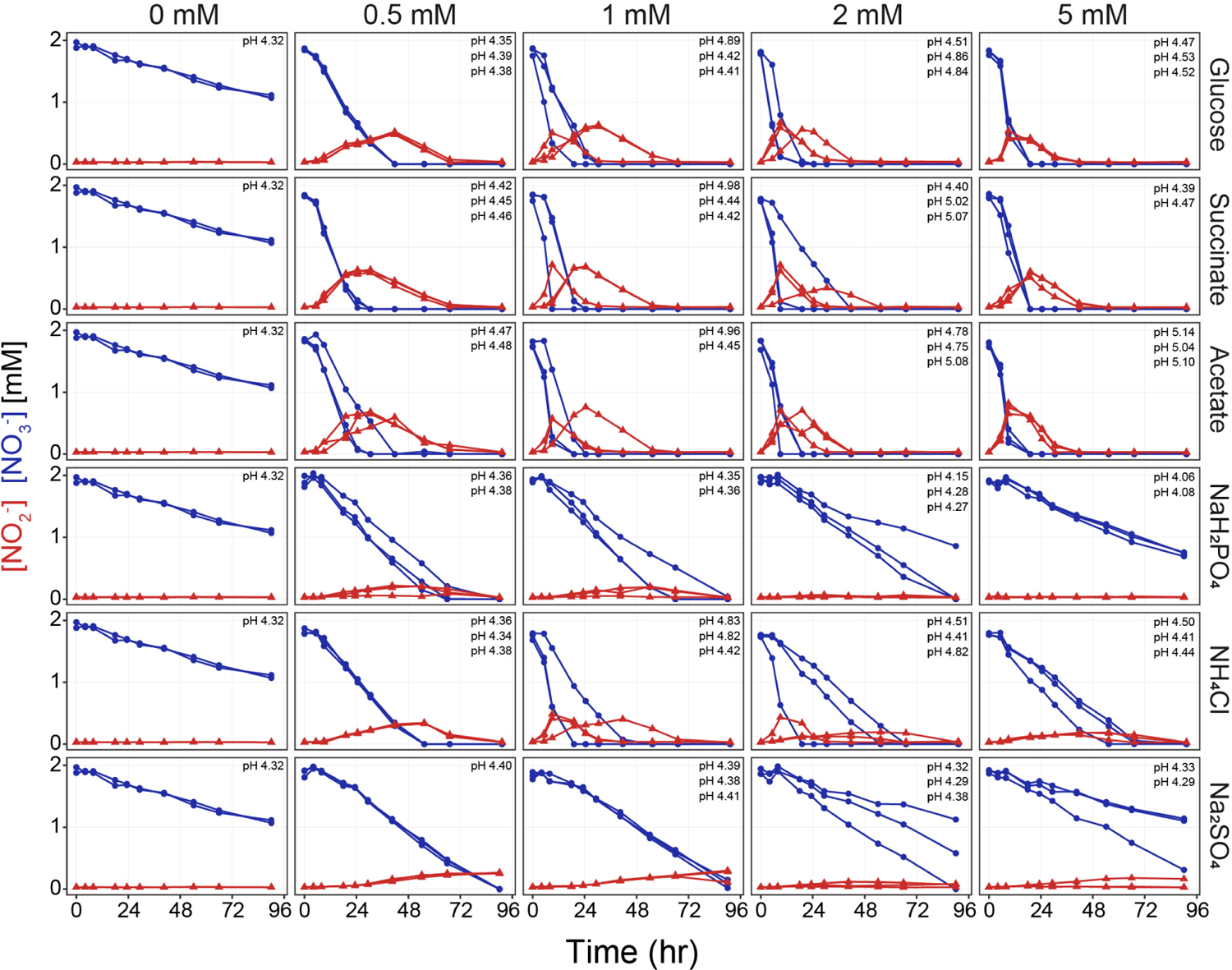
Nitrate and nitrite dynamics of soils amended with different nutrients. To experimentally determine what nutrient was limiting growth in the Nutrient-limiting regime (Regime II), we conducted nutrient amendment experiments respectively with varying concentrations of glucose, succinate, sodium acetate, ammonium chloride (NH_4_Cl), monosodium phosphate (NaH_2_PO_4_), and sodium sulfate (Na_2_SO_4_). Nitrate dynamics (blue) and nitrite dynamics were measured following the same protocol with 2mM NO*^−^* (see Methods) using Soil6 (Table S3) without chloramphenicol and not adding any acid/base. Columns in the plot are different concentrations of C mM, N mM, S mM, or P mM in final concentrations in the slurry varying from 0 to 5 mM, each with biological replicates. 0mM amendment conditions are the same for all nutrients. Rows in the plots are different nutrients: C source (glucose, succinate, acetate), P source (phosphate), N source (ammonium), and S source (Sulfate). Each panel displays the stabilized endpoint pH (1M KCl method), and identical pH values across biological replicates were noted once. Among them, succinate (pK*_a_* = 4.21 and 5.64, 25 *^◦^*C), acetate (pK*_a_* = 4.76, 25 *^◦^*C), and phosphate (pK*_a_* = 2.2, 7.2, and 12.4, 25 *^◦^*C) were strong candidates for the limiting nutrient according to our soil nutrient release hypothesis, due to their anionic nature in mid-range pH (5-7). Because we have previously tested the effect of Na^+^ and Cl*^−^* to be negligible in nitrate dynamics, the effect of these amendments can be attributed solely to C/N/S/P nutrients other than Na^+^ and Cl*^−^*. We observed a transition from linear dynamics to exponential depletion of nitrate, when we amended the soil with a carbon source. Ammonium also made the nitrate consumption dynamics exponential in 1 mM amendment, but not in other amendment concentrations.

**Figure S15:**
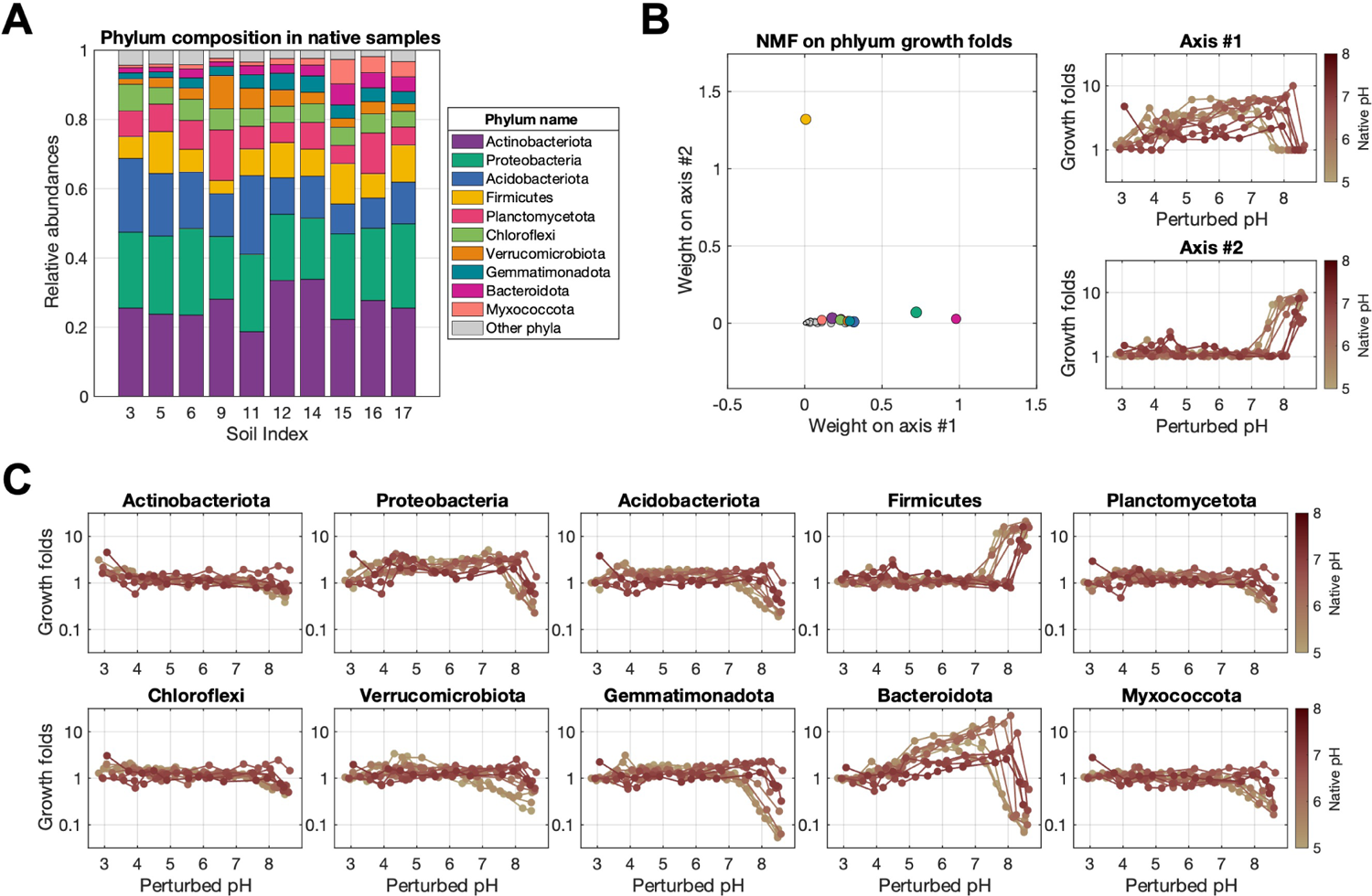
NMF (Non-negative matrix factorization) reveals low-dimensional shifts in growth at the phylum level. **(A)** Initial community composition (*T*_0_) of native soils in the phylum level. The x-axis indicates soils with different native pH levels (Soil 3, 5, 6, 9, 11, 12, 14, 15, 16, 17, see Table S3 for their properties). The y-axis represents the relative abundance (summed to 1) of the top 10 phyla out of 40, with the cumulative abundance of the remaining phyla depicted in gray as ‘Other phyla’. **(C)** By using the absolute abundance of each taxon in chloramphenicol-treated (CHL+) conditions as a baseline value for growth in each perturbed pH condition, we computed the fold increase of each taxon’s absolute abundance in chloramphenicol-untreated (CHL-) conditions, which we call growth fold (*Abs_CHL−_/Abs_CHL_*_+_). Ten different phyla showed idiosyncratic patterns of growth response along the varying perturbed pH. Soils with different native pH, indicated by the line color, showed relatively conserved growth trends in each phylum. **(B)** To systematically identify the underlying lower-dimensional growth response to pH, we used non-negative matrix factorization (NMF) on the growth fold values to decompose the growth response of all phyla into two modes (Axis #1 and Axis #2 in **B**, see Methods for details). Intriguingly, these two response patterns across pH matched the trend of functional parameters fitted with our consumer-resource model respectively for *C*^^^(0) and *x*^(0). The growth folds of each phylum are the linear combination of two modes whose weights are plotted on the left panel of **(B)** (points are colored by phylum as in **(A)**). Firmicutes phylum is mainly composed of mode #2, while other phyla are mainly composed of mode #1. Proteobacteria and Bacteroidota have higher weight #1 than other phyla. Therefore, this enabled us to focus our analysis on these phyla to explain the transition from the Nutrient-limiting regime (Regime II) to the Resurgent growth regime (Regime III).

**Figure S16:**
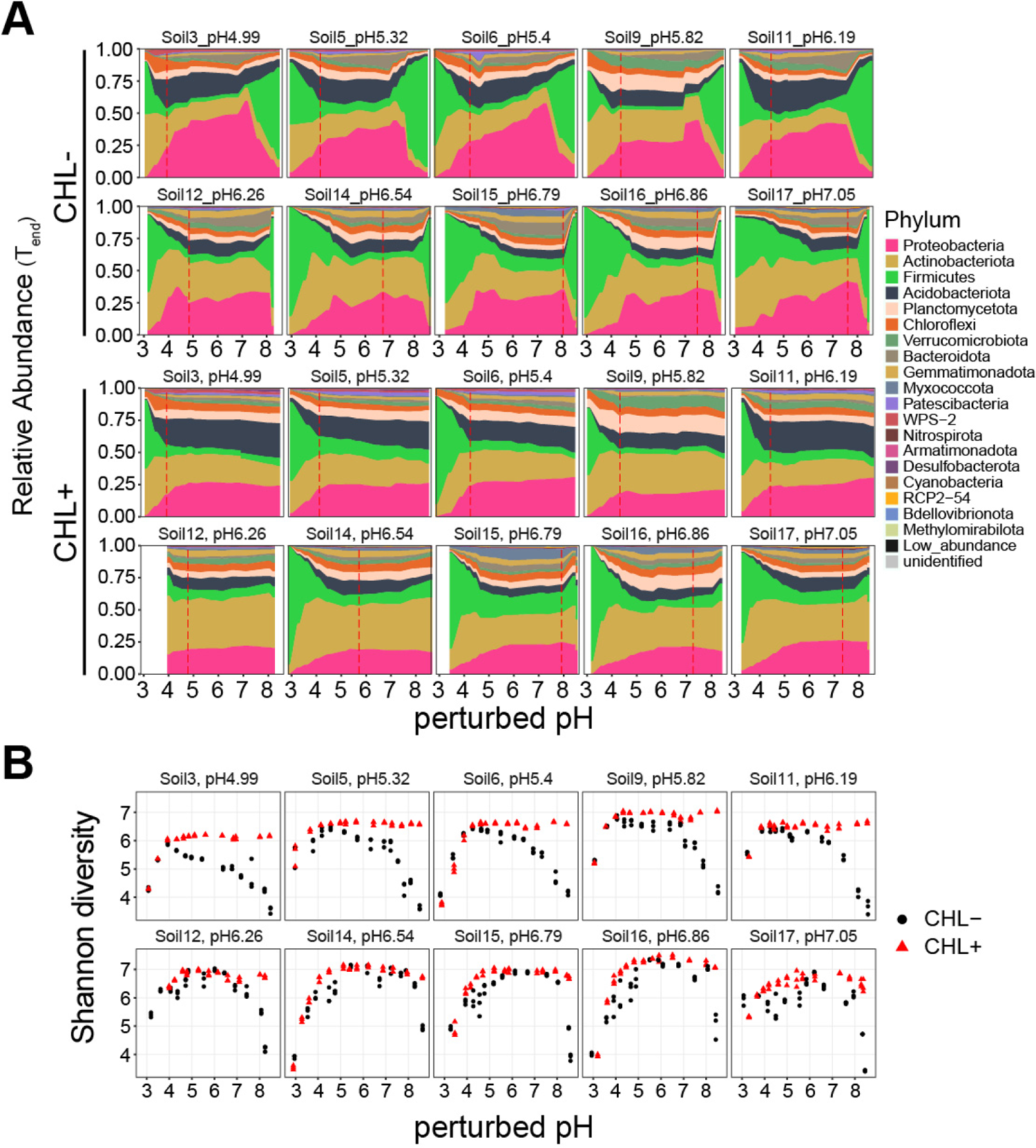
Relative abundance and diversity of different soils across perturbed pH levels. **(A)** Endpoint relative abundance in the phylum level is plotted across the perturbed pH for ten different soils. CHL-indicates endpoint samples without chloramphenicol treatment. CHL+ indicates endpoint samples with chloramphenicol treatment. The alluvial plots were constructed by connecting the relative abundance values of 13 different pH perturbed levels. Red vertical dashed lines indicate the stabilized endpoint pH (1M KCl method) of the unperturbed samples. **(B)** Shannon diversity of the endpoint community is plotted across the perturbed pH for ten different soils in chloramphenicol-untreated(CHL-) and chloramphenicol-treated(CHL+) conditions.

**Figure S17:**
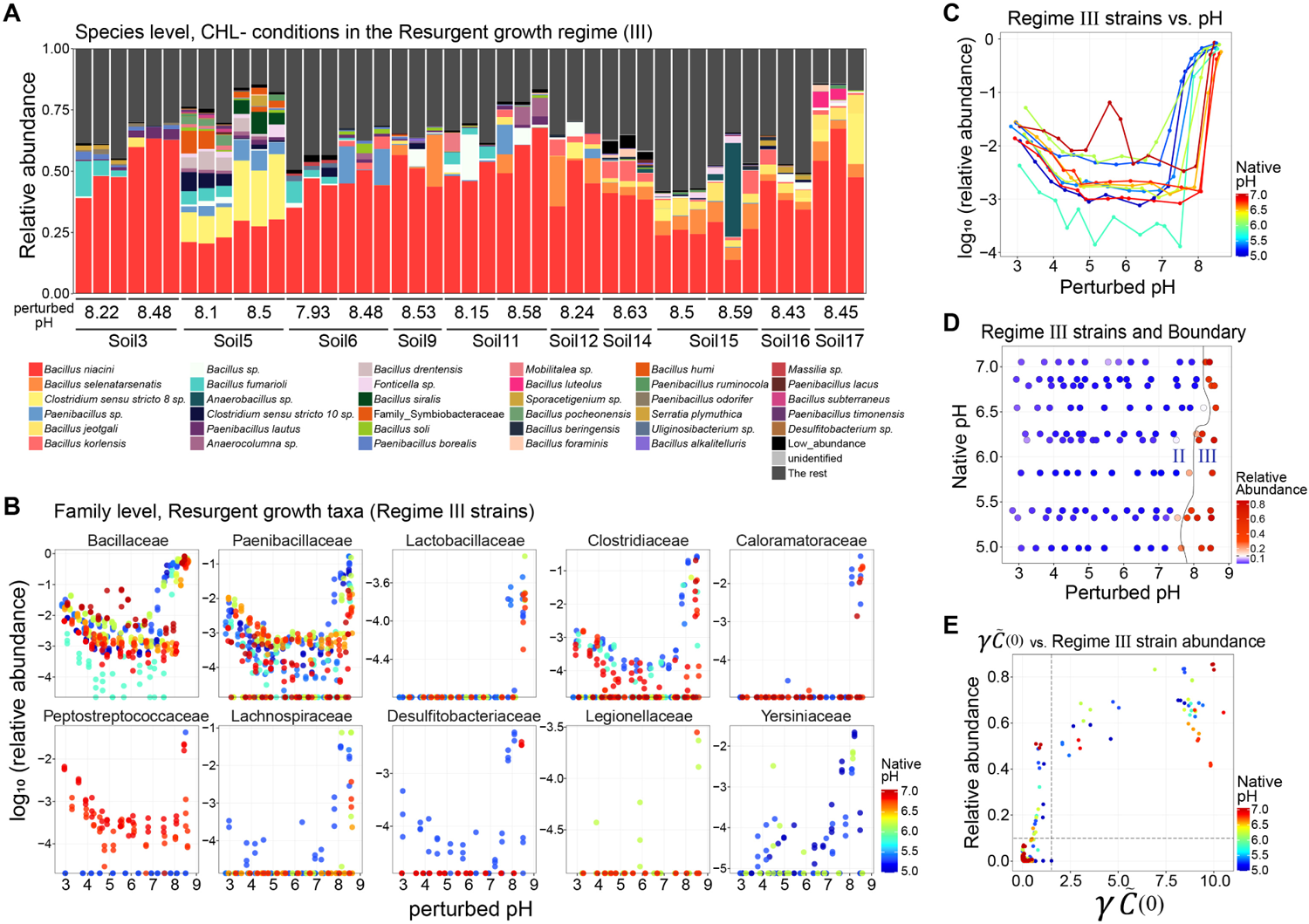
Taxonomy of the identified Resurgent growth strains (Regime III strains) and their abundance agreeing with the functional Regime III. To identify the specific taxa accountable for the emergence of Regime III at a finer taxonomic level, we conducted a differential abundance analysis that statistically determined which Amplicon sequence variants (ASVs) were significantly more abundant in the Regime III CHL- samples, compared to CHL+ samples under same perturbed pH conditions (see Methods). **(A)** The relative abundance of the ASVs in all Regime III samples is highlighted and colored by their assigned species level. The ASVs not significantly enriched in Regime III samples are colored dark gray and labeled as “The rest”. At the genus level, Bacillus, Clostridium, Paenibacillus, and others were identified as the primary contributors to the Resurgent growth regime (Regime III) as plotted. **(B)** The analysis revealed that 10 families belonging to Firmicutes (Bacillaceae, Paenibacillaceae, Clostridiaceae, Caloramatoraceae, Peptostreptococcaceae, etc.) and 2 families belonging to Proteobacteria phylum (Legionellaceae and Yersiniaceae) significantly enriched in the Resurgent growth regime (Regime III). Their relative abundance (log_10_ scale) increases at basic perturbed pH levels, patterns differing in soils with different native pH levels. Their relative abundance also slightly increases in Regime I, due to their high tolerance to pH perturbations. **(C-D)** Then, we aggregated the relative abundance of these differential ASVs (i.e., Regime III strains) to assess their contribution to the emergence of Regime III. Notably, their abundances (log_10_ scale) rise between pH 7-8, which aligns with or slightly precedes the transition between Regime II and III. **(E)** This increase in relative abundance corresponded with the rise of the nutrient growth parameter *γC*^^^_0_from zero.

**Figure S18:**
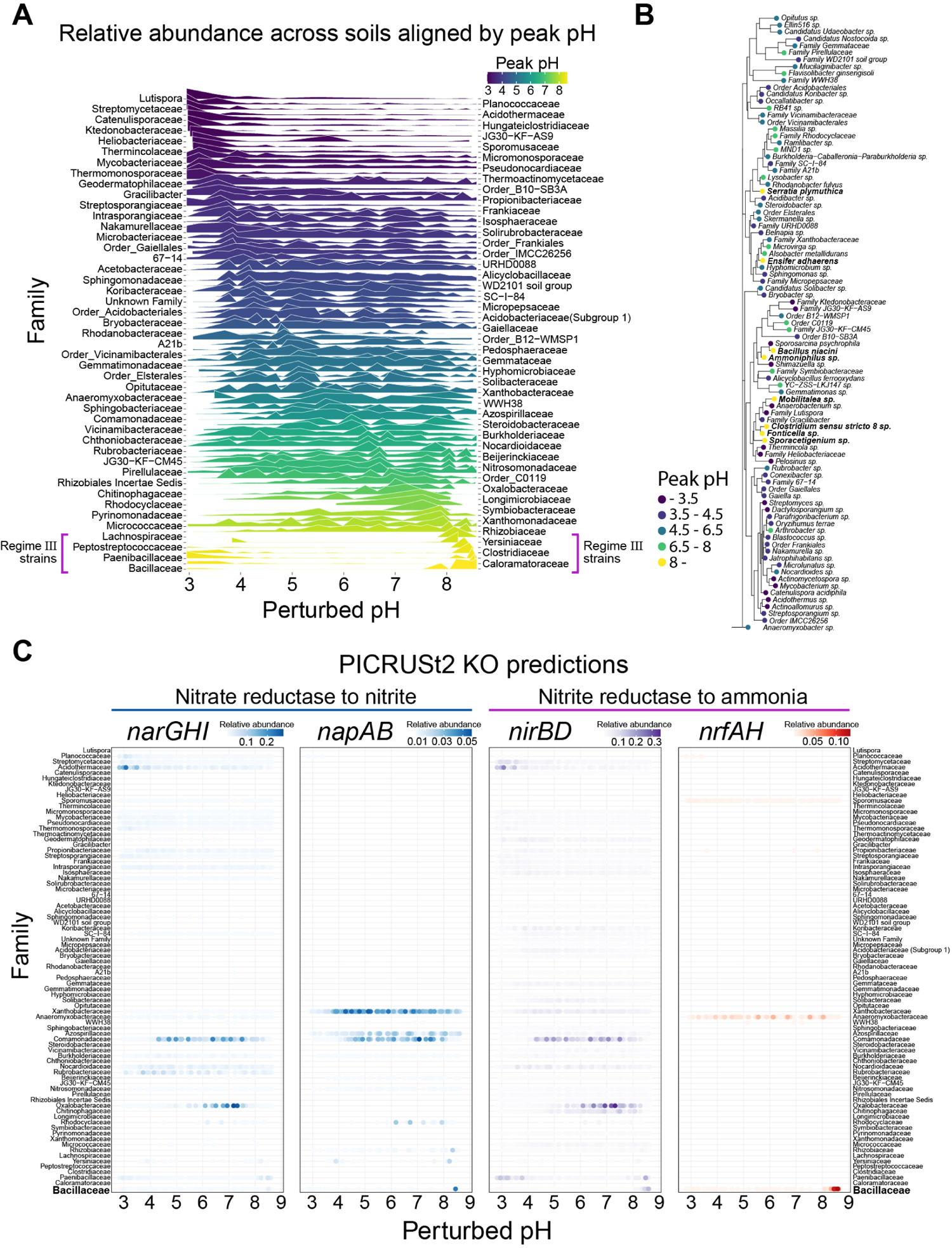
Traits of the Resurgent growth strains (Regime III strains) are analyzed through pH niche, phylogenetic distance, and gene predictions from PICRUSt2. **(A)** To elucidate the pH niche of all taxa, we analyzed the relative abundance of amplicon sequence variants (ASVs) identified as being enriched in different pH levels (see Differential abundance analysis in Methods). We aggregated the relative abundance of ASVs in the same family for each sample, and then computed the median relative abundance of samples across different soils belonging to each pH bin (see Methods). The families were ordered by their peak pH (acidic peak pH on top with dark blue color and basic peak pH on the bottom with yellow color), which was the pH corresponding to the pH bin with the highest median relative abundance for each family. Indeed, the families belonging to the Resurgent growth strains had a peak pH over 8: Bacillaceae, Clostridiaceae, Paenibacillaceae, Caloramatoraceae, Peptostreptococcaceae, Lachnospiraceae (Firmicutes phylum), and Yersiniaceae (Proteobacteria phylum). **(B)** To see whether there was phylogenetic convergence among strains with similar pH niches, we used the 16S rRNA sequences of the ASVs to construct a phylogenetic tree. We selected one representative ASV with the largest relative abundance from each family to represent the family and used its 16S rRNA V34 region sequence to construct the phylogenetic tree. (see Method). Each node is labeled by the genus or species name and colored by its peak pH. The Resurgent growth strains (yellow-colored) did not cluster phylogenetically and were dispersed throughout the phylogenetic tree. **(C)** To infer genotypes of the Resurgent growth strains (Regime III strains), we used PICRUSt2 to predict the KEGG ortholog (KO) gene abundance from the 16S rRNA sequence of each ASV (see Methods). We focused on KOs/genes related to denitrification and Dissimilatory Nitrate Reduction to Ammonium (DNRA): nitrate reductase in denitrification (*narG*, *narH*, *narI*, *napA*, *napB*) and nitrite reductase to ammonium (*nirB*, *nirD*, *nrfA*, *nrfH*). To track which KOs were enriched at which pH in the 89 families used in the peak pH analysis in **A**, we summed the relative abundance (reads / total reads of each perturbed pH level in CHL-samples) of the ASVs belonging to each family that possessed at least 1 predicted gene respectively for *narGHI*, *napAB*, *nirBD*, and *nrfAH*. Then, we plotted their relative abundance values across pH for all soils, indicated by the intensity of the point’s colors. For the Resurgent growth strain, Bacilli family exhibited a notable enrichment in *nrfAH* genes (indicated by red points), which are DNRA-related genes producing ammonium from nitrite.

**Figure S19:**
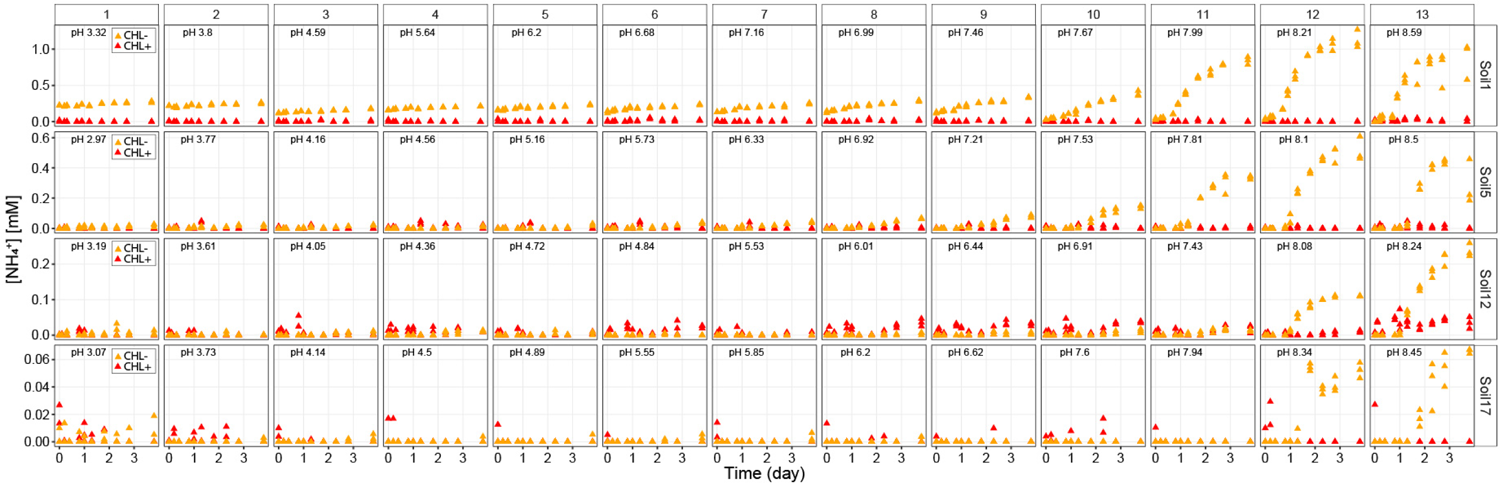
Ammonium dynamics in chloramphenicol-untreated (CHL-) and treated (CHL+) samples in 4 soils with different native pH levels. For Soil 1, 5, 12, and 17 (for details of soils, refer to Table S3), the ammonium dynamics were measured colorimetrically using the Salicylate-hypochlorite assay [75]. Ammonium dynamics show that varying levels of NH^+^ (from 3% (Soil17) to 50% (Soil1) of the provided 2mM NO_3_*^−^* converted to NH_4_^+^) accumulated in the Resurgent growth regime (Regime III). This indicates the activation of dissimilatory nitrate to ammonia (DNRA) pathway by a subset of strains responsible for the Resurgent growth regime. Chloramphenicol treatments in the samples (CHL+) led to consistent detection of 0.5mM NH_4_^+^ due to its N-H moiety. The NaOH concentration in perturbed samples also impacted NH_4_^+^ measurements, because the Salicylate-hypochlorite assay includes a step that OCl*^−^* reacts with the N-H moiety resulting in N-Cl and OH*^−^*, higher NaOH concentrations resulting in slightly lower detection of chloramphenicol in the CHL+ samples (0.45mM NH^+^ in 100mM NaOH perturbations). Taking advantage of these control measurements, we used the constant NH_4_^+^ levels in the controls without 2mM NO_3_*^−^* (No-Nitrate controls) in the CHL+ conditions for each soil to offset the NaOH effect in the CHL- samples (by computing the conversion factor ratio of NH_4_^+^ levels of No-Nitrate controls in CHL+ conditions to the initial NH_4_^+^ levels of each condition with different NaOH additions in CHL+ samples) and to subtract NH^+^ levels caused by chloramphenicol in CHL+ samples. pH indicated inside each panel is the endpoint stabilized pH (see Methods).

**Figure S20:**
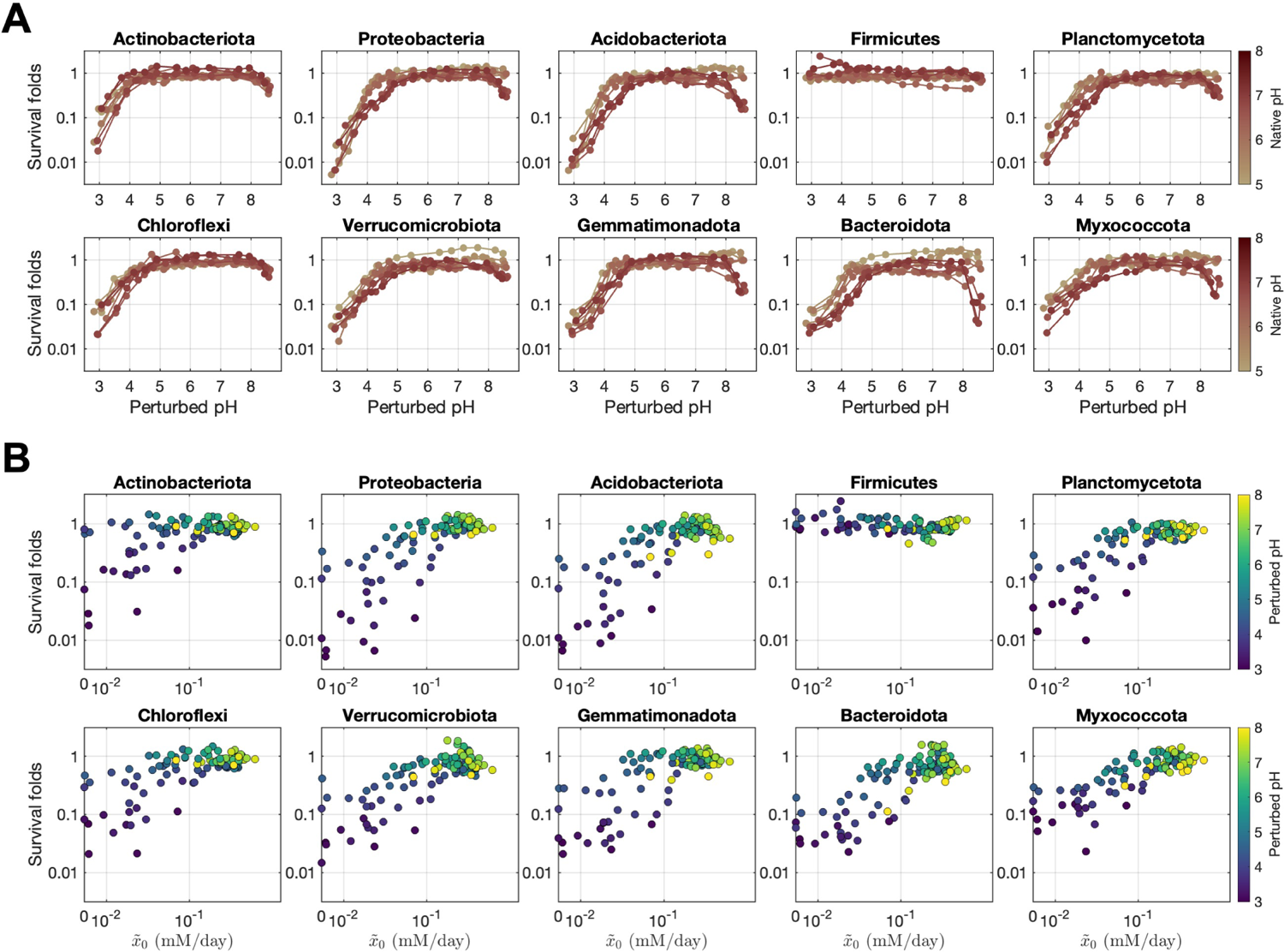
Death of phyla during acidic pH perturbations explains Acidic death regime (Regime I) **(A)** To infer death, the survival folds of the top 10 phyla (relative abundance-wise) among 40 taxa were determined for each perturbed pH condition by computing the fold difference in the endpoint absolute abundance of each Phylum under CHL+ conditions, relative to their baseline levels at the initial time point (*Abs_CHL_*_+_*/Abs_T_*_0_). We used the abundance in CHL+ conditions to rule out growth and only compute the death effect of pH. A consistent drop of survival folds during acidic perturbation was observed across all phyla except for the Firmicutes phylum. **(B)** To check if the sequencing data supports our model parameter fitted from functional dynamics, we plotted the survival folds against the fitted *x*^(0) parameter (indigenous biomass activity) for each perturbed pH level denoted by the color gradient. We removed Regime III data points to focus on the Acidic death regime (Regime I). Indeed, during acidic perturbations (dark blue points), the survival folds decreased with the indigenous biomass activity *x*^(0) parameter fitted with functional data, except for only the Firmicutes. These results suggest that the impacts of acidic and basic pH perturbations on death are asymmetric rather than symmetric: while acidic conditions cause widespread death, death from basic conditions is less prominent.

**Figure S21:**
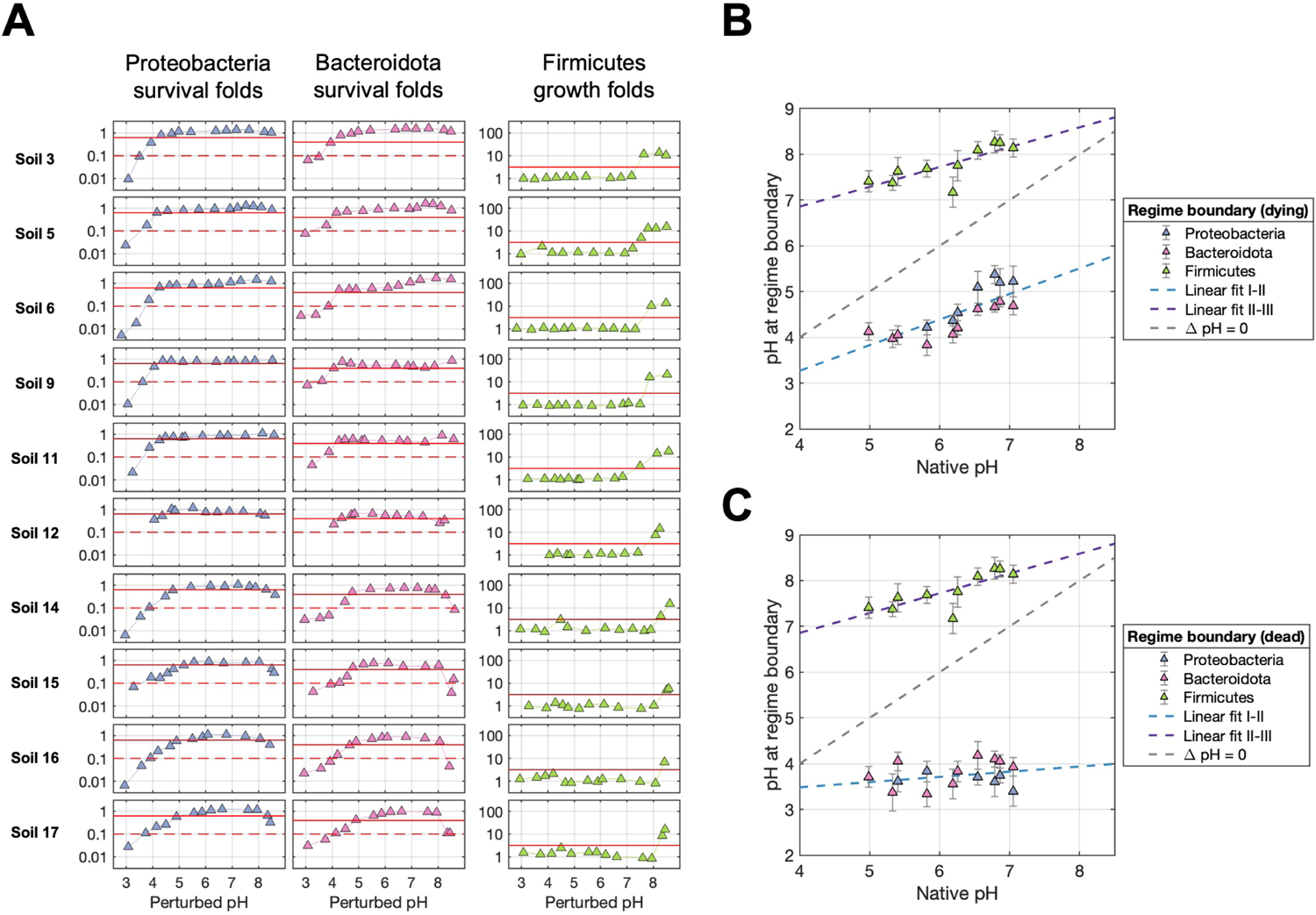
Inferring regime boundary with phyla abundance dynamics. To understand the transition to the Acidic death regime (Regime I) with sequencing data, we computed the survival folds of Proteobacteria and Bacteroidota phyla across perturbed pH levels, which is the absolute abundance ratio of *Abs_CHL_*_+_*/Abs_T_*_0_ (endpoint absolute abundance in CHL+ conditions compared to the initial time point (*T*_0_) for each perturbed pH level) (blue and pink data points in the left two panels). We set an identical survival fold threshold for all soils (red lines) to compute the pH at which the survival fold goes below that threshold during acidic perturbation. We used two distinct definitions to choose a threshold for the survival fold; (1) the “dying” definition where the taxa’s abundance started to decline in abundance compared to *T*_0_ (survival fold threshold *<* 1, red solid lines), and (2) the “dead” definition where the taxa’s abundance was close to 0 (survival fold threshold *→* 0, red dashed lines). To understand the transition to the Resurgent growth regime (Regime III) with sequencing data, we computed the growth folds of the Firmicutes phylum (green data point in the rightmost panel) by endpoint absolute abundance ratio of *Abs_CHL−_/Abs_CHL_*_+_ (chloramphenicol untreated/treated conditions), and similarly computed the boundary pH at which the growth folds started increasing (threshold = 3, red solid lines). **(B)** These pH transition points were plotted against the native soil pH level. For the pH transition points in Proteobacteria and Bacteroidota, we used the “dying” definition. Employing the ‘dying’ definition with Proteobacteria, Bacteroidota allowed us to recapitulate the phenomenon observed in the functional data, where the fitted slope of Boundary I-II was less than 1, as shown in Fig. 6A. This suggests that these phyla in the relatively neutral soil are more tolerant of larger ΔpH change until they start to die than those in acidic soils, possibly due to variations in soil titration curves (Fig. 6B). Because the fitted slope is greater than 0, this also means that these phyla in relatively acidic soils can tolerate lower acidic pH conditions than those in neutral soils, which signals long-term pH adaptation. **(C)** The pH transition points were plotted against the native soil pH level using the “dead” definition for Proteobacteria and Bacteroidota. The “dead” definition threshold resulted in a flat slope close to 0. This suggests that, despite long-term adaptation to varying native soil pH levels, these taxa have similar pH thresholds at which complete decimation occurs. For **B and C**, error bars represent the pH difference between the two samples neighboring the survival or growth fold threshold. The points indicate the mid-point pH value of these boundary samples. The linear fit was determined using a least squares method (blue and purple dashed lines). The grey dashed line represents y = x, indicating hypothetical points where there is no difference between native and perturbed pH values (ΔpH = pH*_perturbed_ −*pH*_native_* = 0).

**Figure S22:**
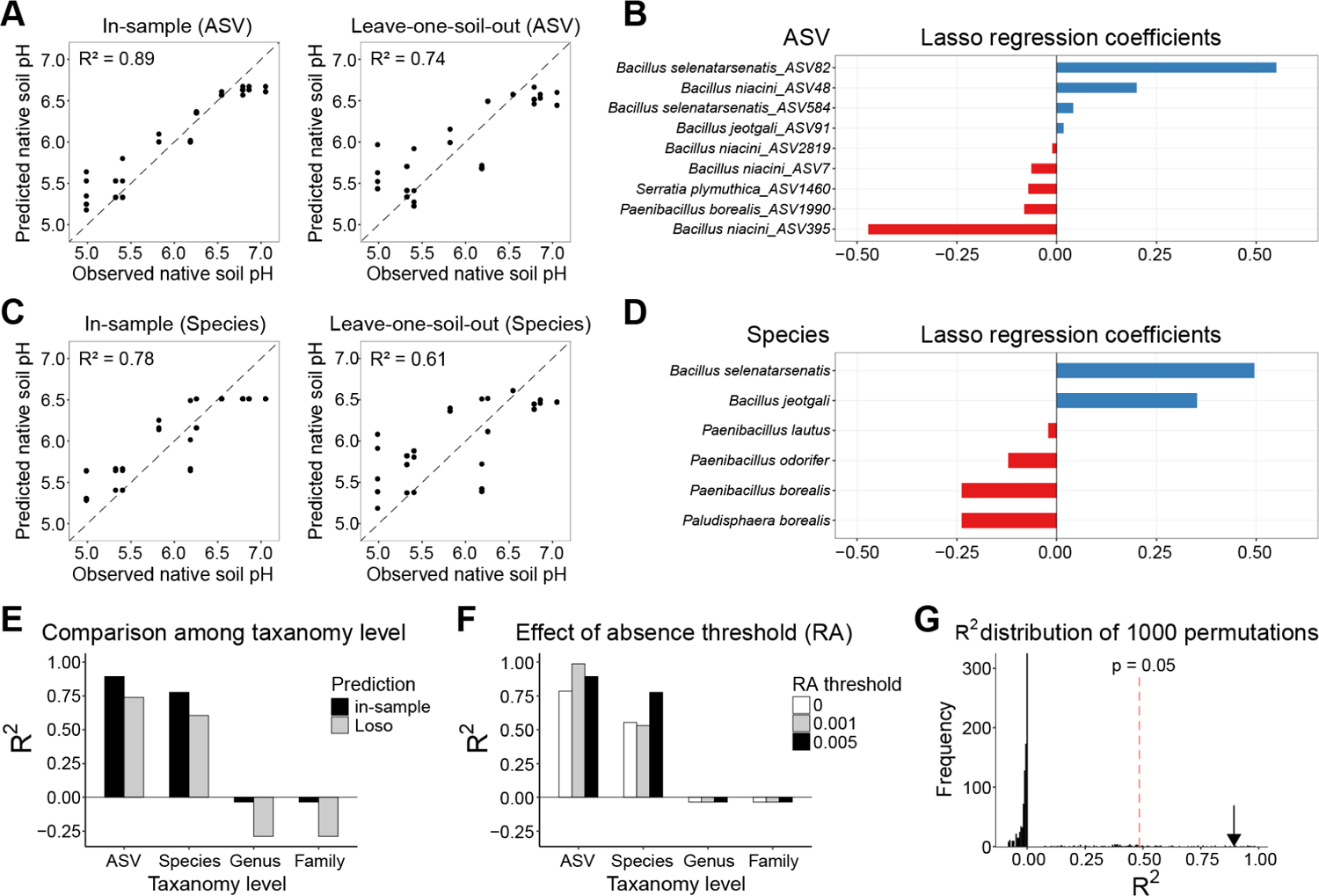
Strain (ASV) and species-level variation of the Resurgent growth strains arise from the soil’s natural pH environment. To investigate whether the taxonomic identity of Resurgent growth (Regime III) strains is determined by long-term adaptation to or selection from different natural pH environments, we performed a regularized regression analysis to see if we can predict the native pH level of the source soil from the presence or absence of Resurgent growth strains across different taxonomic levels (Amplicon sequence variant (ASV), Species, Genus, Family, or higher taxonomic levels) (see Methods). **(A)** Predicted and observed native soil pH with the Lasso regression from the presence and absence of the ASVs of Resurgent growth strains, using 0.005 (out of the relative abundance of 1) as a threshold for presence (see Methods). Left is the in-sample predictions, and the right is ‘Leave-one-soil-out’ (Loso) predictions where we leave out samples from one soil when we fit the regression model and then use the left-out samples to make predictions of their native pH as shown in the scatter plot. The prediction quality (R^2^) was computed using the mean predicted and mean observed native pH levels for each soil. **(B)** Bar plots indicate the regression coefficients of all ASVs with non-zero coefficients from in-sample predictions in **A**. **(C)** Predicted and observed native soil pH from the Lasso regression from the presence and absence of the Resurgent growth strains in the species level. **(D)** Bar plots indicate the regression coefficients of all species with non-zero coefficients from in-sample predictions in **C**. **(E)** In-sample and Loso predictions are good only until the ASV and species level. From the genus level or higher, the predictions are worse than random (negative R^2^ values). A relative abundance (RA) threshold of 0.005 was used for the presence and absence. **(F)** Effect of RA threshold values for the presence and absence (0, 0.001, and 0.005 out of 1). In the case of in-sample predictions, imposing the RA threshold improved the predictions at the ASV and species level. **(G)** To ascertain our prediction quality is not an artifact, we randomly permuted the native pH values 1000 times, and then predicted in-sample the native pH, showing that the R^2^ value of 0.89 from in-sample predictions from **A** is greater than the top 50th R^2^ value (0.475) out of 1000 shuffled predictions (p = 0.05).

**Figure S23:**
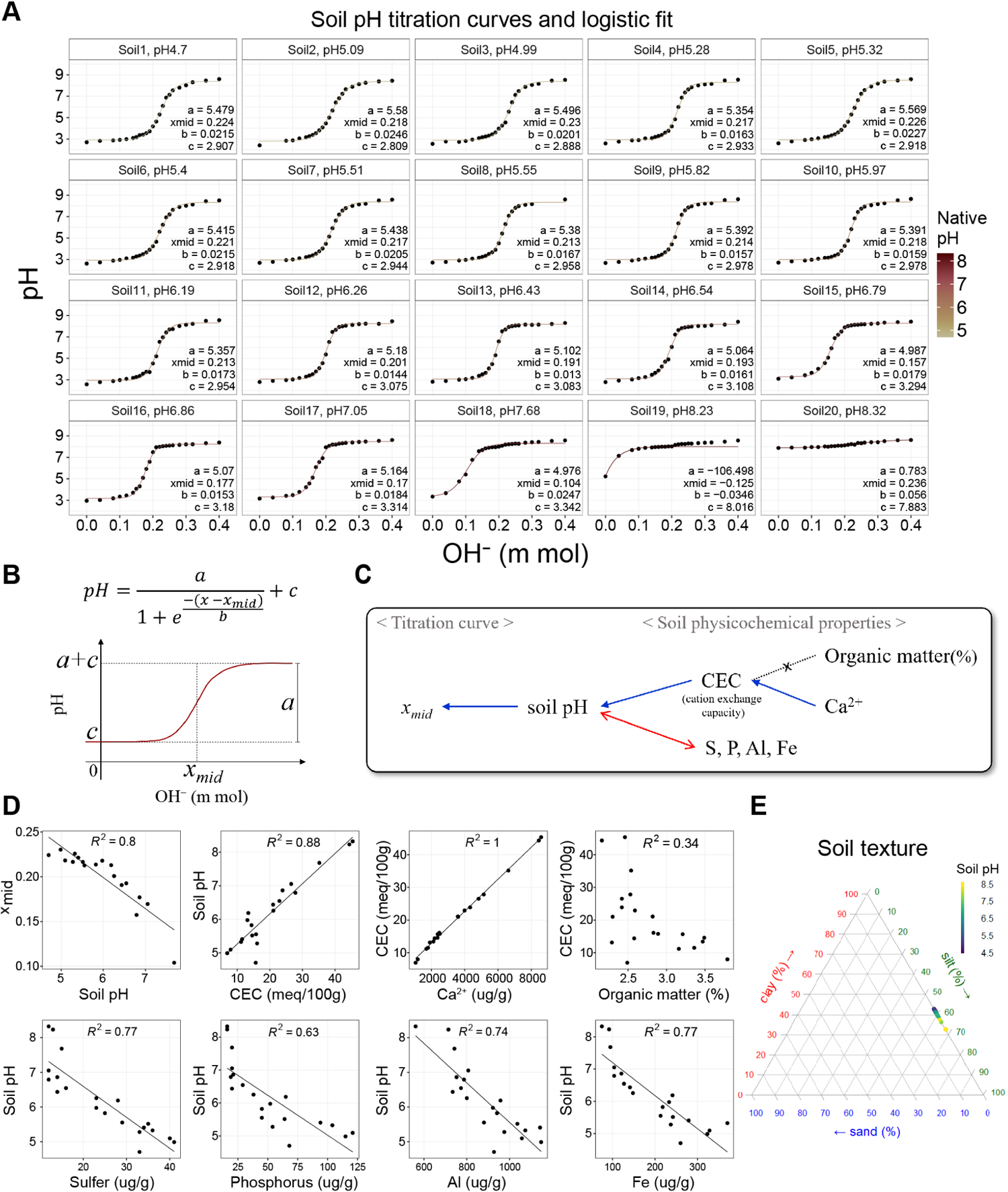
Relationship between pH titration curves and soil physicochemical properties. **(A)** Fitting logistic function to pH titration curves of the 20 soils from different native pH levels (see Methods). **(B)** Logistic function and parameters. **(C)** Summary of how soil’s physicochemical properties can influence the soil pH titration curves. We can attribute the horizontal shift of the pH titration curves to their varying native soil pH levels, which are potentially determined by the Cation exchange capacity (CEC) and the *Ca*^2+^ ion concentrations. **(D)** The correlations that support the claim from the summary diagram **C** are shown with R^2^ values. **(E)** Soil texture (sand, silt, clay percentage composition) was mostly identical for different soils, thereby not explaining the difference in soil pH levels and the titration curves.

**Figure S24:**
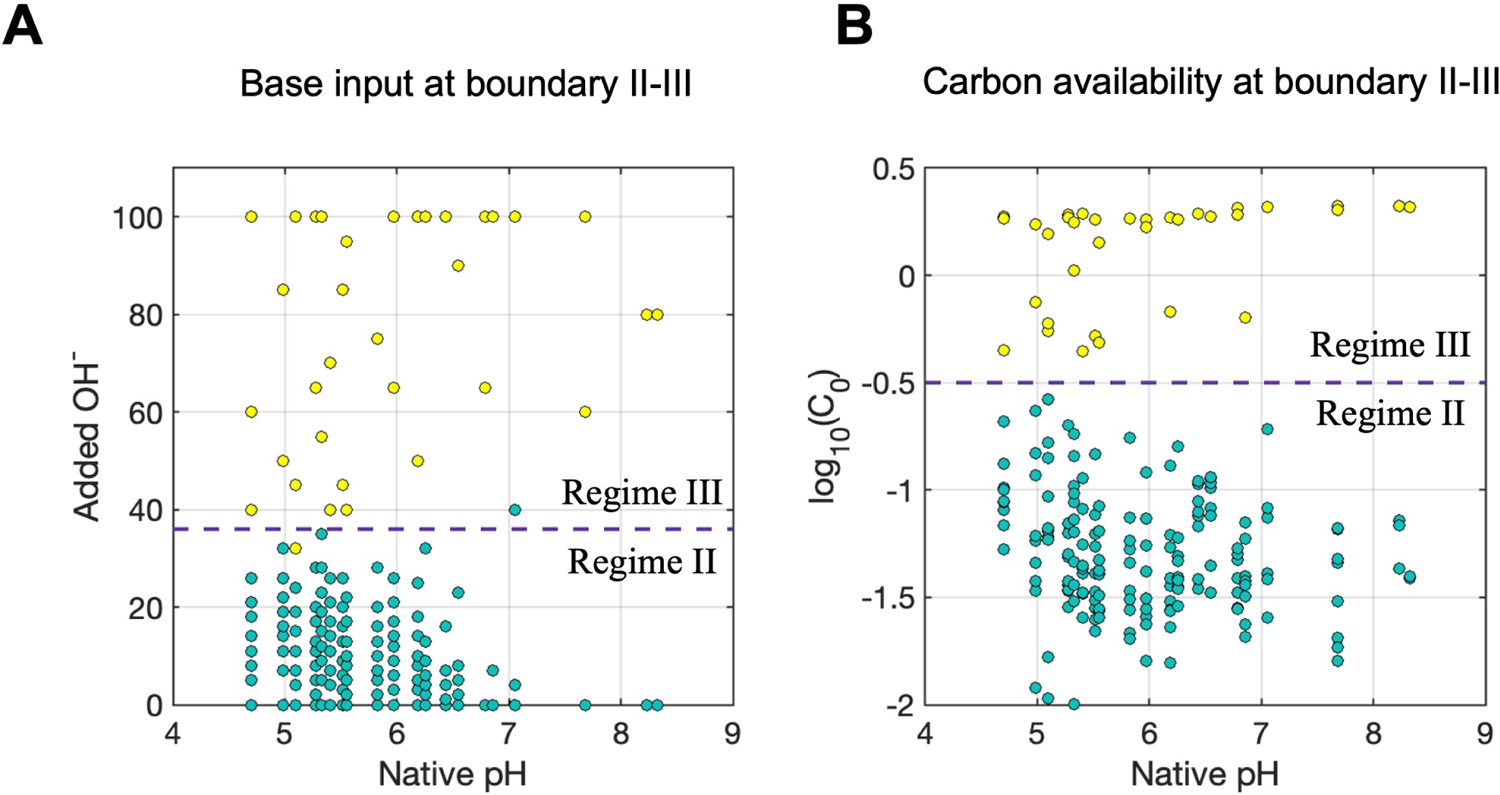
Functional regime boundary II-III is dictated by the amount of NaOH and the available carbon nutrients. **(A)** The relationship of the native soil pH and the amount of NaOH input (y-axis in mM) in Regime II samples (green data points) and Regime III (yellow data points). The dashed purple horizontal line indicates the NaOH input required to transition from Regime II to Regime III. The rather flat slope of pH boundary II-III vs. native pH in Fig. 6A can be explained by the fixed amount of NaOH input (dashed purple line in Fig. 6B). From our previous results, the amount of available carbon corresponds to the NaOH input. **(B)** Therefore, we plotted the relationship of the native soil pH and the fitted *C*^^^(0) values (log scale). We indeed observe the consistent amount of available carbon required to transition from Regime II to Regime III (dashed purple horizontal line).

**Figure S25:**
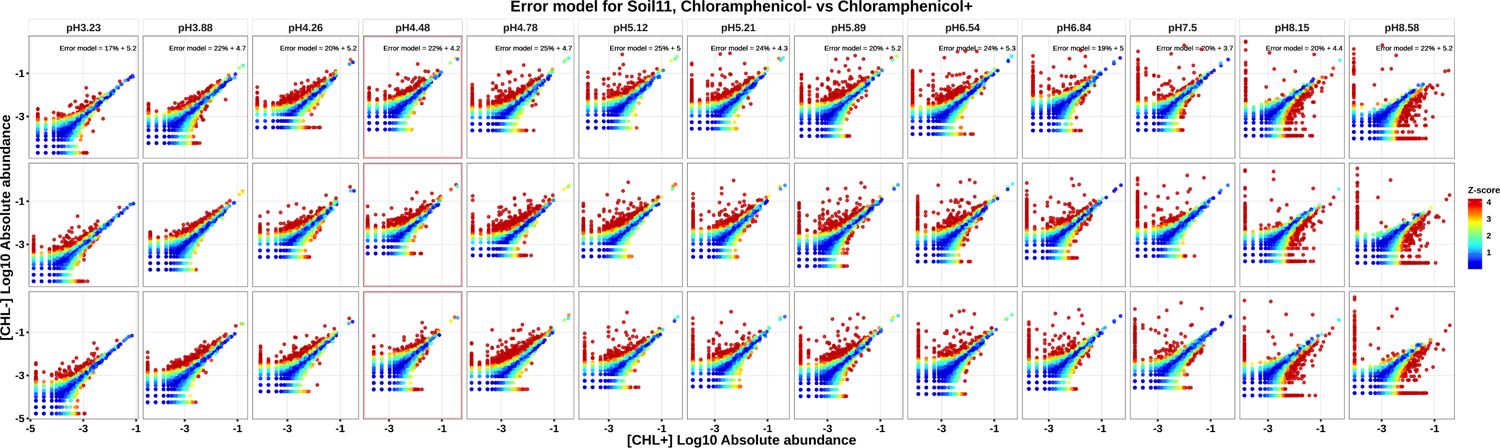
Error model z score. To identify the ASVs enriched for each perturbed pH level, we empirically constructed a null model with the three biological replicates (see Methods). For each soil (Soil11 shown in the plot) and for each perturbed pH level (pH indicated in the header of each panel), we plot the log-scale absolute abundance of each ASV in chloramphenicol-treated samples (CHL+, x-axis) against the absolute abundance in chloramphenicol-untreated samples (CHL-, y-axis). Three rows of the panel indicate three biological replicate pairs. The deviations of replicate-replicate comparisons from 1:1 line are well-described by an effective model combining two independent contributions, a Gaussian noise of fractional magnitude *c*_frac_ and a constant Gaussian noise of magnitude *c*_0_ reads, such that repeated measurements (over biological r<u>eplicates) of</u> an ASV with mean abundance *n* counts are approximately Gaussian-distributed with a standard deviation of *σ*(*c*_0_*, c*_frac_) = (*c*_frac_*n*)^2^ + *c*^2^ counts. In this expression, *c*_frac_ was estimated from moderate-abundance ASVs (*>* 50 counts) for which the other noise term is negligible; and *c*_0_ was then determined as the value for which 67% of replicate-replicate comparisons are within *±σ*(*c*_0_*, c*_frac_) of each other, as expected for 1-sigma deviations. This noise model was inferred separately for each soil and each perturbed pH level, as the corresponding samples were processed independently in different sequencing runs. The noise parameters are denoted inside the panel. The points are colored by the z-score computed from the Gaussian-distributed error model. The pink peripheral box indicates the condition without any acid/base addition.

## Notes

### Competing Interest Statement

The authors have declared no competing interest.

### Summary of Updates

Updated Title, Abstract, Introduction, and Discussion.

